# Gene expression heterogeneity during brain development and aging: temporal changes and functional consequences

**DOI:** 10.1101/595249

**Authors:** Ulaş Işıldak, Mehmet Somel, Janet M. Thornton, Handan Melike Dönertaş

## Abstract

Cells in largely non-mitotic tissues such as the brain are prone to stochastic (epi-)genetic alterations that may cause increased variability between cells and individuals over time. Although increased inter-individual heterogeneity in gene expression was previously reported, whether this process starts during development or if it is restricted to the aging period has not yet been studied. The regulatory dynamics and functional significance of putative aging-related heterogeneity are also unknown. Here we address these by a meta-analysis of 19 transcriptome datasets from diverse human brain regions. We observed a significant increase in inter-individual heterogeneity during aging (20+ years) compared to postnatal development (0 to 20 years). Increased heterogeneity during aging was consistent among different brain regions at the gene level and associated with lifespan regulation and neuronal functions. Overall, our results show that increased expression heterogeneity is a characteristic of aging human brain, and may influence aging-related changes in brain functions.

---

Aging is a complex process characterized by a gradual decline in maintenance and repair mechanisms, accompanied by an increase in genetic and epigenetic mutations, and oxidative damage to nucleic acids, protein and lipids^1,2^. The human brain experiences dramatic structural and functional changes in the course of aging. These include decline in gray matter and white matter volumes^3^, increase in axonal bouton dynamics^4^ and reduced synaptic plasticity, all processes that may be associated with decline in cognitive functions^5^. Changes during brain aging are suggested to be a result of stochastic processes, unlike changes associated with postnatal neuronal development that are known to be primarily controlled by adaptive regulatory processes^6–8^. The molecular mechanisms underlying age-related alteration of regulatory processes and eventually leading to aging-related phenotypes, however, are little understood.

Over the past decade, a number of transcriptome studies focusing on age-related changes in human brain gene expression profiles were published^2,9–12^. These studies report aging-related differential expression patterns in many functions, including synaptic functions, energy metabolism, inflammation, stress response, and DNA repair. By analyzing age-related change in gene expression profiles in diverse brain regions, we previously showed that for many genes, gene expression changes occur in opposite directions during postnatal development (pre-20 years of age) and aging (post-20 years of age), which may be associated with aging-related phenotypes in healthy brain aging^13^. While different brain regions are associated with specific, and often independent, gene expression profiles^9,10,12^, these studies also show that age-related alteration of gene expression profiles during aging is a widespread effect across different brain regions.

One of the suggested effects of aging is increased variability between individuals and somatic cells, which has been previously reported by several studies. Some of these studies find an increase in age-related heterogeneity in heart, lung and white blood cells of mice^14–16^, *Caenorhabditis elegans*^17^, and human twins^18^. A study analysing microarray datasets from different tissues of humans and rats also reported an increase in age-related heterogeneity in expression as a general trend^19^, although this study found no significant consistency across datasets, nor any significant enrichment in functional gene groups. That said, the generality of increase in expression heterogeneity remains unresolved. For instance, Viñuela et al. find more decrease than an increase in heterogeneity in human twins^20^ and Ximerakis et al. show the direction of the heterogeneity change depends on cell type in aging mice brain^21^. Using GTEx data covering different brain regions (20 to 70 years of age), Brinkmeyer-Langford et al. identify a set of differentially variable genes between age groups, but they do not observe increased heterogeneity at old age^22^. Meanwhile, another study performing single-cell RNA sequencing of human pancreatic cells, identifies an increase in transcriptional heterogeneity and somatic mutations with age^23^. A meta-analysis also suggested more shared expression patterns during development than in aging, implying an increase in inter-individual variability^13^. Likewise, a prefrontal cortex transcriptome analysis we recently conducted revealed a weak increase in age-dependent heterogeneity at the gene, transcriptome and pathway levels, irrespective of the preprocessing methods^24^.

Whether age-related increase in heterogeneity is a universal phenomenon thus remains contentious. Furthermore, where it can be detected, whether this is a time-dependent process that starts at the beginning of life or whether this increase and its functional consequences are only seen after developmental processes are completed, have not yet been explored. In this study, we retrieved transcriptome data from independent studies covering the whole lifespan, including data from diverse brain regions, and conducted a comprehensive analysis to identify the prevalence of age-related heterogeneity changes in human brain aging compared with those observed during postnatal development. We confirmed that increased age-related heterogeneity is a consistent trend in the human brain transcriptome during aging but not during development, and it is associated with the pathways and biological functions that are related to longevity and neuronal function.

## Results

To investigate how heterogeneity in gene expression changes with age, we used 19 published microarray datasets from three independent studies. Datasets included 1,010 samples from 17 different brain regions of 298 individuals whose ages ranged from 0 to 98 years (Supplementary Table S1, Fig. S1). In order to analyze the age-related change in gene expression heterogeneity during aging compared to the change in development, we divided datasets into two subsets as development (0 to 20 years of age, *n* = 441) and aging (20 to 98 years of age, *n* = 569). We used the age of 20 to separate pre-adulthood and adulthood based on commonly used age intervals in earlier studies (see Methods). For the analysis, we focused only on the genes for which we have a measurement across all datasets (*n* = 11,137).

### Age-related change in gene expression levels

To quantify age-related changes in gene expression, we used a linear model between gene expression levels and age (see Methods, Supplementary Fig. S2). We transformed the ages to the fourth root scale before fitting the model as it provides relatively uniform distribution of sample ages across the lifespan, but we also confirmed that different age scales yield quantitatively similar results (see Supplementary Fig. S3). We quantified expression change of each gene in aging and development periods separately and considered regression coefficients from the linear model (*β* values) as a measure of age-related expression change (Supplementary Fig. S4).

We first analyzed similarity in age-related expression changes across datasets by calculating pairwise Spearman’s correlation coefficients among the *β* values (Figure 1a). Both development (median correlation coefficient = 0.56, permutation test *p* < 0.001, Supplementary Fig. S5) and aging datasets (median correlation coefficient = 0.43, permutation test *p* = 0.003, Supplementary Fig. S5) showed moderate correlation with the datasets within the same period. Although the difference between dataset correlations within development and aging datasets was not significant (permutation test *p* = 0.1, Supplementary Fig. S6), weaker consistency during aging may reflect the stochastic nature of aging, causing increased heterogeneity between aging datasets.

**Figure 1.**
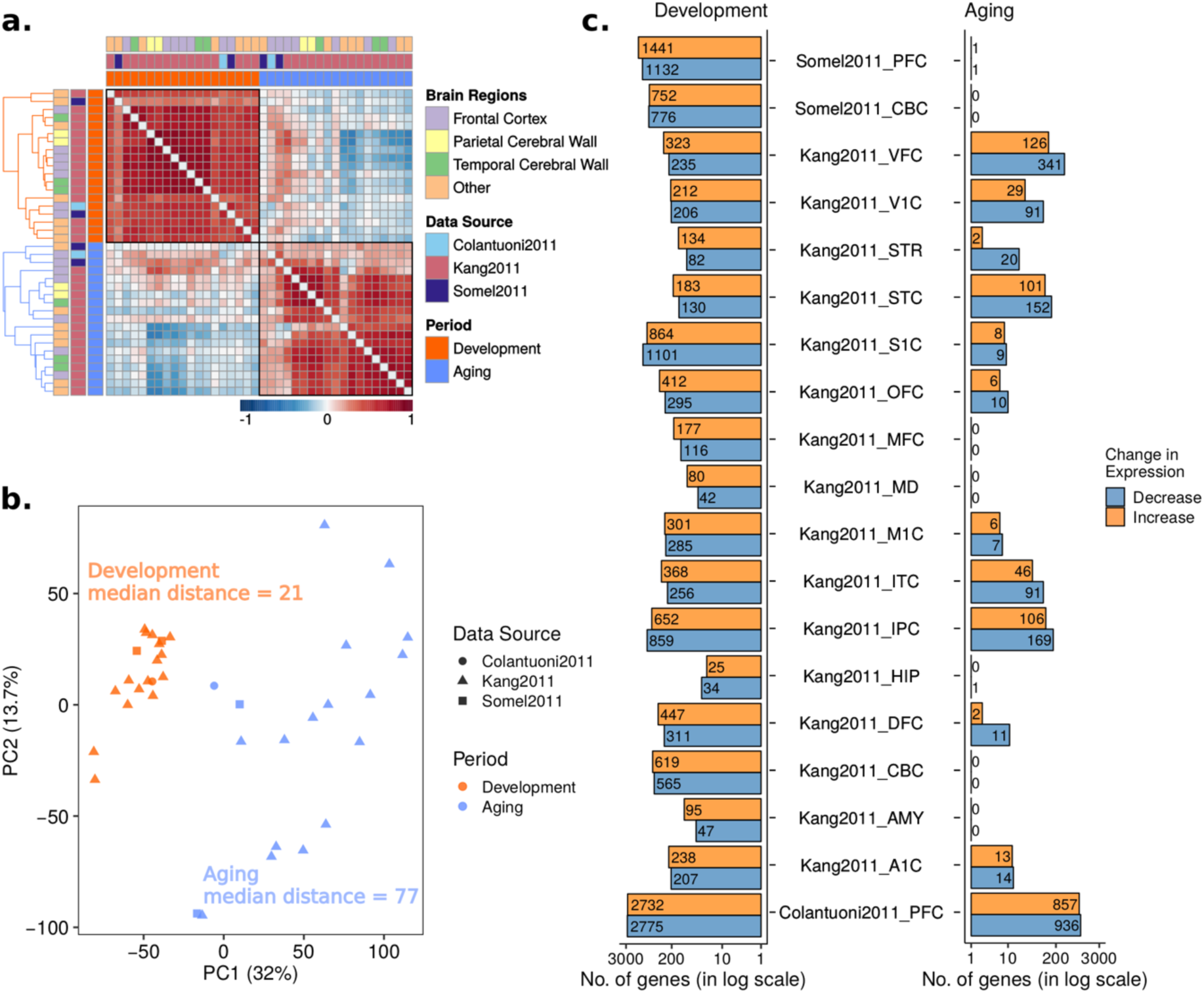
Age-related change in gene expression during postnatal development and aging. (a) Spearman correlations among age-related expression changes (β values) across datasets. The color of the squares indicates if the correlation between the corresponding pair of datasets (across β values of 11,137 common genes) is positive (red) or negative (blue), while darker color specifies a stronger correlation. Diagonal values were removed in order to enhance visuality. Annotation rows and columns indicate data source, brain region and period of each dataset. Hierarchical clustering was performed for each period separately (color of the dendrogram indicates periods) to determine the order of datasets. (b) Principal component analysis (PCA) of age-related expression changes during aging and development. The analysis was performed on age-related expression change values of 11,137 common genes among all 38 datasets. The values of the first principal component on the x-axis and second principal component on the y-axis were drawn, where the values in the parenthesis indicate the variation explained by the corresponding principal component. Median Euclidean pairwise distances among development and aging datasets calculated using PC1 and PC2 were annotated on the figure. Different shapes show different data sources and colors show development (dark orange) and aging (blue) (c) Number of significant (FDR corrected p < 0.05) gene expression changes in development (left panel) and aging (right panel). The x-axis shows the number of genes in the log scale. The color of the bars shows the direction of change, decrease (steel gray), and increase (orange). The exact number of genes are also displayed on the plot.

The principal component analysis (PCA) of age-related expression changes (*β*) revealed distinct clusters of development and aging datasets (Figure 1b). Moreover, aging datasets were more dispersed than development datasets (median pairwise Euclidean distances between PC1 and PC2 were 77 for aging and 21 for development), which may again reflect stochasticity in gene expression change during aging and can indicate more heterogeneity among different brain regions or datasets during aging than in development.

We next identified genes showing significant age-related expression change (FDR-corrected *p* < 0.05), for development and aging datasets separately (Figure 1c). Development datasets showed more significant changes compared to aging (permutation test *p* = 0.003, Supplementary Fig. S6), which may again indicate higher expression variability among individuals during aging. The direction of change in development was mostly positive (14 datasets with more positive and 5 with more negative), whereas in aging datasets, we observed more genes with a decrease in expression level (13 datasets with more genes decreasing expression and 5 with no significant change, and 1 with an equal number of positive and negative changes).

### Age-related change in gene expression heterogeneity

To assess age-related change in heterogeneity, we obtained the unexplained variance (residuals) from the linear models used to calculate the change in gene expression level. For each gene in each dataset, we separately calculated Spearman’s correlation coefficient (*ρ*) between the absolute value of residuals and age, irrespective of whether the gene shows a significant change in expression (see Methods, Supplementary Fig. S2). We considered *ρ* values as a measure of heterogeneity change, where positive values mean an increase in heterogeneity with age. We also repeated this approach using loess regression instead of a linear model between expression level and age, and found high correspondence between *ρ* values based on linear and loess regression models (Supplementary Fig. S7). Still, loess regression was more sensitive to the changes in sample sizes and parameters and we therefore continued downstream analyses with the *ρ* estimates based on the residuals from the linear model.

We next asked if datasets show similar *ρ, i.e.* age-related changes in heterogeneity, by calculating pairwise Spearman’s correlation between pairs of datasets, across shared genes (Figure 2a). Unlike the correlations among expression level changes, *ρ* values did not show a higher consistency during development. In fact, although the difference is not significant (permutation test *p* = 0.2, Supplementary Fig. S6), the median value of the correlation coefficients was higher in aging (median correlation coefficient = 0.21, permutation test *p* = 0.24, Supplementary Fig. S5), than in development (median correlation coefficient = 0.11, permutation test *p* = 0.25, Supplementary Fig. S5).

**Figure 2.**
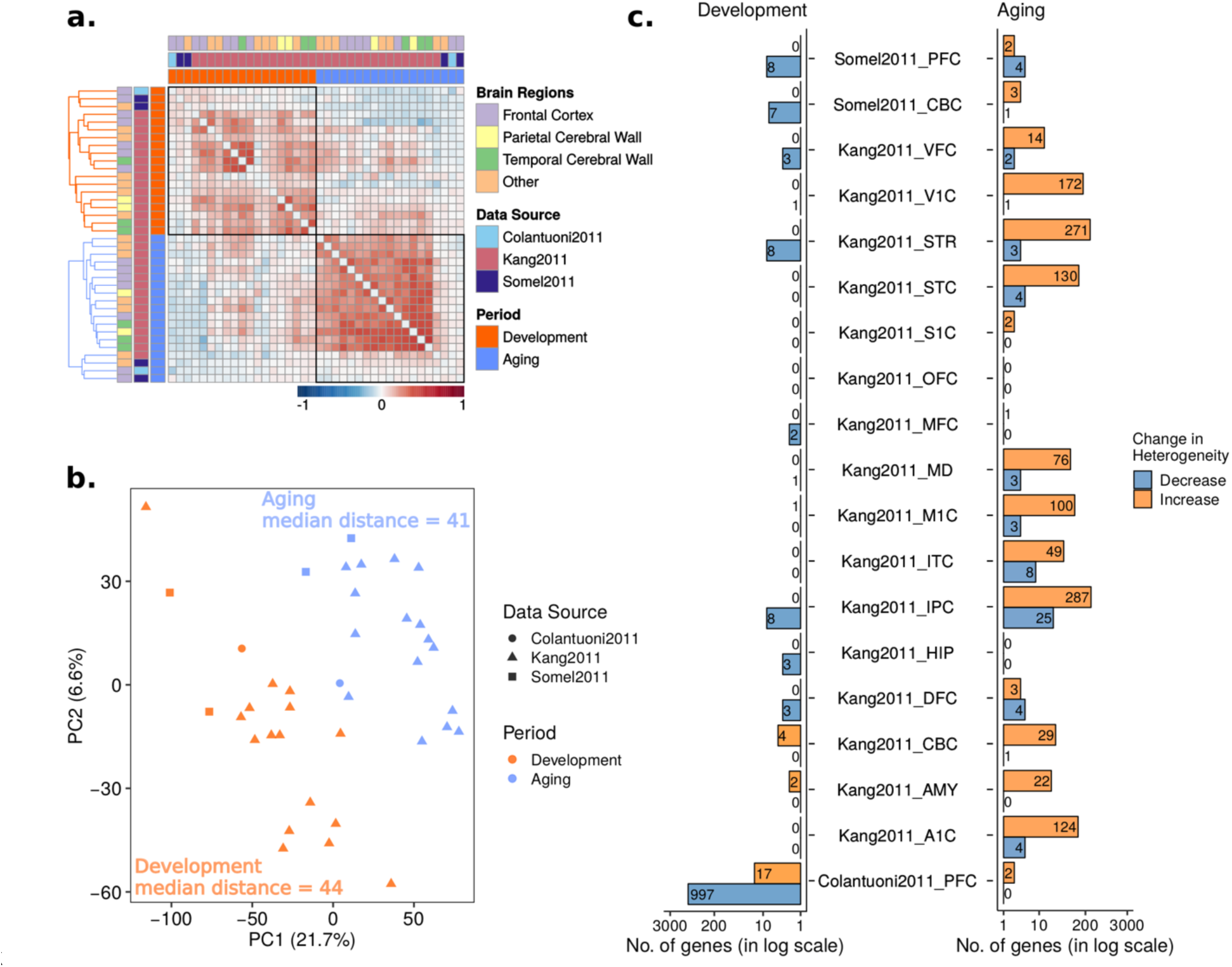
Age-related change in gene expression heterogeneity during development and aging. The procedures are similar to those in Figure 1, except, age-related heterogeneity changes (ρ values) were used instead of expression changes (β values). (a) Spearman correlations among age-related heterogeneity changes (ρ values) across datasets. (b) Principal component analysis (PCA) of heterogeneity change with age. (c) The number of genes showing significant heterogeneity change in aging and development.

A principal component analysis (PCA) showed that, like expression change, heterogeneity change with age can also differentiate aging datasets from development (Figure 2b). Similar to the pairwise correlations (Figure 2a), aging datasets clustered more closely than development datasets (median pairwise Euclidean distances between PC1 and PC2 are 41 and 44 for aging and development, respectively). Both observations imply more similar changes in heterogeneity during aging.

Using the *p*-values from Spearman’s correlation between age and the absolute value of residuals for each gene, we then investigated the genes showing a significant change in heterogeneity during aging and development (FDR corrected *p*-value < 0.05). We found almost no significant change in heterogeneity during development, except for the Colantuoni2011 dataset, for which we have high statistical power due to its large sample size. In aging datasets, on the other hand, we observed more genes with significant changes in heterogeneity (permutation test *p* = 0.06, Supplementary Fig. S6) and the majority of the genes with significant changes in heterogeneity tended to increase in heterogeneity (Figure 2c). However, the genes showing a significant change did not overlap across aging datasets (Supplementary Fig. S8).

Nevertheless, our analyses indicated relatively more consistent heterogeneity changes among datasets in aging compared to development, implying that heterogeneity change could be a characteristic linked to aging (see Discussion).

### Consistent increase in heterogeneity during aging

As our previous analyses suggested age-related changes in heterogeneity can differentiate development from aging and show more similarity during aging, we sought to characterize the genes displaying such changes. Since the significance of the changes is highly dependent on the sample size, instead of focusing on significant genes identified within individual datasets, we leveraged upon the availability of multiple datasets and focused on their shared trends, capturing weak but reproducible trends across multiple datasets. Consequently, we used the level of consistency in age-related heterogeneity change across datasets to sort genes.

We first examined profiles of age-related heterogeneity change in aging and development. Among aging datasets 18/19 showed more increase than decrease in heterogeneity with age (median *ρ* > 0, *i.e.* higher numbers of genes with increase), while the median heterogeneity change in one dataset was zero. In development, on the other hand, only 5/19 datasets showed more increase in heterogeneity, while the remaining 14/19 datasets showed more decrease with age (median *ρ* < 0) (Figure 3a). The age-related change in heterogeneity during aging was significantly higher than during development (permutation test *p* < 0.001, Supplementary Fig. S6). We also checked if there is a relationship between changes in heterogeneity during development and during aging (*e.g.* if those genes that decrease in heterogeneity tend to increase in heterogeneity during aging) but did not find any significant trend (Supplementary Fig. S9).

**Figure 3.**
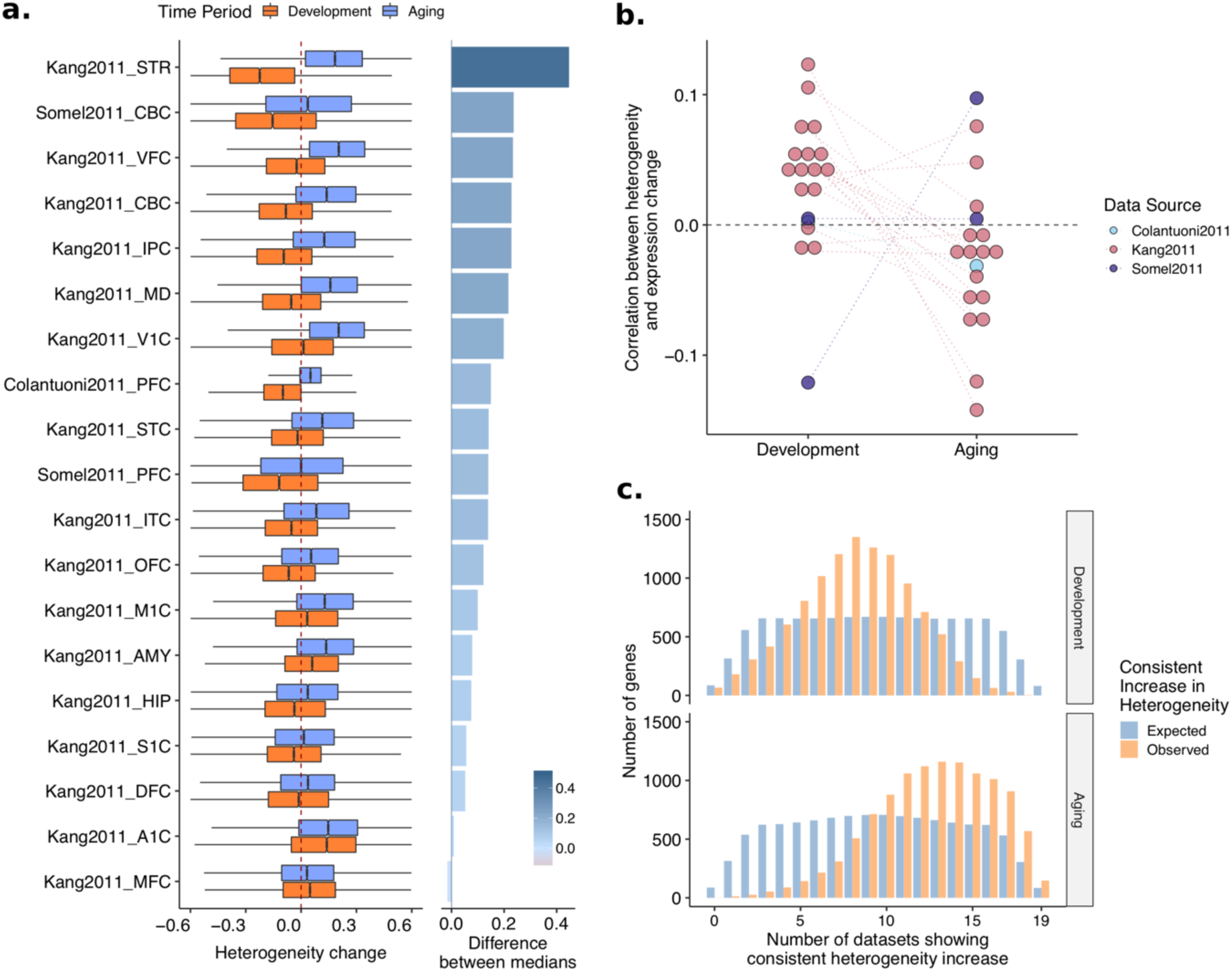
(a) Boxplots, showing distributions of age-related heterogeneity changes (ρ values) of 11,1137 common genes for each dataset and period separately. The dotted red line (vertical line at x = 0) reflects no change in heterogeneity. The difference between median heterogeneity change in aging and development is given as a bar plot on the right panel. Datasets are ordered by the differences in median heterogeneity changes in aging and development. (b) The relationship between expression and heterogeneity change with age. Spearman correlation analysis was performed between age-related expression changes (β values) and age-related heterogeneity changes (ρ values) of 11,137 common genes, separately for each dataset. The dotted gray line at y = 0 reflects no correlation between expression and heterogeneity. (c) Expected and observed consistency in the heterogeneity change across datasets in development and aging. There is a significant shift toward heterogeneity increase in aging (permutation test p<10^-7^) (lower panel), while there is no significant consistency in either direction in development (upper panel). The expected distribution is constructed using a permutation scheme that accounts for the dependence among datasets and is more stringent than random permutations (see Supplementary Fig. S10 for details).

A potential explanation why we see different patterns of heterogeneity change with age in development and aging could be the accompanying changes in the expression levels, as it is challenging to remove dependence between the mean and variance. To address this possibility, we first calculated Spearman’s correlation coefficient between the changes in heterogeneity (*ρ* values) and expression (*β* values), for each dataset. Overall, all datasets had values close to zero, suggesting the association is not strong. Surprisingly, we saw an opposing profile for development and aging; while the change in heterogeneity and expression were positively correlated in development, they showed a negative correlation in aging (Figure 3b).

Having observed both a tendency to increase and a higher consistency in heterogeneity change during aging, we asked which genes show consistent increase in heterogeneity across datasets in aging and development. We therefore calculated the number of datasets with an increase in heterogeneity during development and aging for each gene (Figure 3c). To calculate significance and expected consistency, while controlling for dataset dependence, we performed 1,000 random permutations of individuals’ ages and re-calculated the heterogeneity changes (see Methods). In development, there was no significant consistency in heterogeneity change in either increase or decrease. During aging, however, there was a significant signal of consistent heterogeneity increase, *i.e.* more genes showed consistent heterogeneity increase across aging datasets than randomly expected (Figure 3c, lower panel). We identified 147 common genes with a significant increase in heterogeneity across all aging datasets (permutation test *p* < 0.001, Supplementary Table S2). Based on our permutations, we estimated that 84/147 genes could be expected to have consistent increase just by chance, suggesting only ∼40% true positives. In development, in contrast, there was no significant consistency in heterogeneity change in either direction (increase or decrease). Nevertheless, comparing the consistency in aging and development, there was an apparent shift towards a consistent increase in aging – even if we cannot confidently report the genes that become significantly more heterogeneous with age across multiple datasets.

### Heterogeneity Trajectories

We next asked if there are specific patterns of heterogeneity change, *e.g.* increase only after a certain age. We used the genes with a consistent increase in heterogeneity with age during aging (*n* = 147) to explore the trajectories of heterogeneity change (Figure 4). Genes grouped with k-means clustering revealed three main patterns of heterogeneity increase (Supplementary Table S2): i) genes in clusters 3 and 7 show noisy but *a steady increase* throughout aging, ii) genes in clusters 4, 5 and 8 show *increase in early aging but a later slight decrease*, revealing a reversal (up-down) pattern, and iii) the remaining genes in cluster 1, 2 and 6 *increase in heterogeneity dramatically after the age of 60*. We next asked if these genes have any consistent heterogeneity change pattern in development (Supplementary Fig. S11), but most of the clusters showed no or only weak age-related changes during development. We also analyzed the accompanying changes in mean expression levels for these clusters. Except for cluster 1, which shows a decrease in expression level at around the age of 60 and then shows a dramatic increase, all clusters show a steady scaled mean expression level at around zero, *i.e.* different genes in a cluster show different patterns (Supplementary Fig. S12).

**Figure 4.**
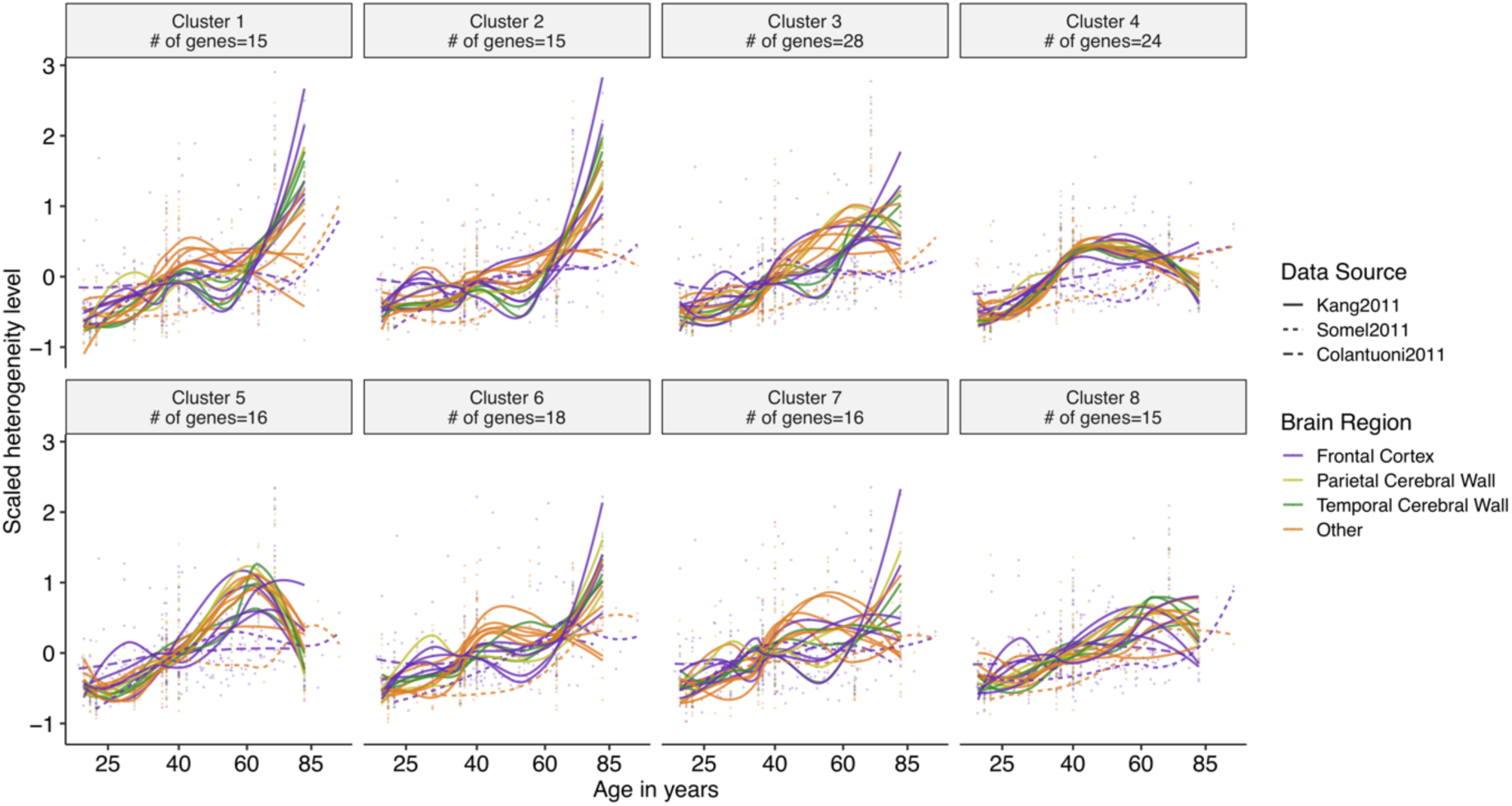
Clusters of genes showing a consistent heterogeneity increase in aging (n = 147). Clustering was performed based on patterns of the change in heterogeneity, using the k-means clustering method (see Methods). The x- and y-axes show age and heterogeneity levels, respectively. Mean heterogeneity change for the genes in each cluster was drawn by spline curves. The colors and line-types of curves specify different brain regions and data sources, respectively.

We further tested the genes showing dramatic heterogeneity increase after the age of 60 (clusters 1, 2 and 6) for association with Alzheimer’s Disease, as the disease incidence increases after 60^25^ as well; however, we found no evidence for such an association (see Supplementary Fig. S13).

### Functional analysis

To examine the functional associations of heterogeneity changes with age, we performed gene set enrichment analysis using KEGG pathways^26^, Gene Ontology (GO) categories^27,28^, Disease Ontology (DO) categories^29^, Reactome pathways^30^, transcription factor (TF) targets (TRANSFAC)^31^, and miRNA targets (MiRTarBase)^32^. Specifically, we rank-ordered genes based on the number of datasets that show a consistent increase in heterogeneity and asked if the extremes of this distribution are associated with the gene sets that we analyzed. There was no significant enrichment for any of the functional categories and pathways for the consistent changes in development. The significantly enriched KEGG pathways for the genes that become consistently heterogeneous during aging included multiple KEGG pathways known to be relevant for aging, including the longevity regulating pathway, autophagy^33^, mTOR signaling^34^ and FoxO signaling^35^ (Figure 5a). Among the pathways with a significant association (listed in Figure 5a), only protein digestion and absorption, primary immunodeficiency, linoleic acid metabolism, and fat digestion and absorption pathways had negative enrichment scores, meaning these pathways were significantly associated with the genes having the least number of datasets showing an increase. However, it is important to note that this does not mean these pathways have a decrease in heterogeneity as the distribution of consistent heterogeneity levels is skewed (Figure 3c, lower panel). We also calculated if the KEGG pathways that we identified are particularly enriched in any of the heterogeneity trajectories we identified. Although we lack the necessary power to test the associations statistically due to small number of genes, we saw that i) group 1, which showed a stable increase in heterogeneity, is associated more with the metabolic pathways and mRNA surveillance pathway, ii) group 2, which showed first an increase and a slight decrease at later ages, is associated with axon guidance, mTOR signaling, and phospholipase D signaling pathways, and iii) group 3, which showed a dramatic increase after age of 60, is associated with autophagy, longevity regulating pathway and FoxO signaling pathways. The full list is available as Supplementary Figure S14.

**Figure 5.**
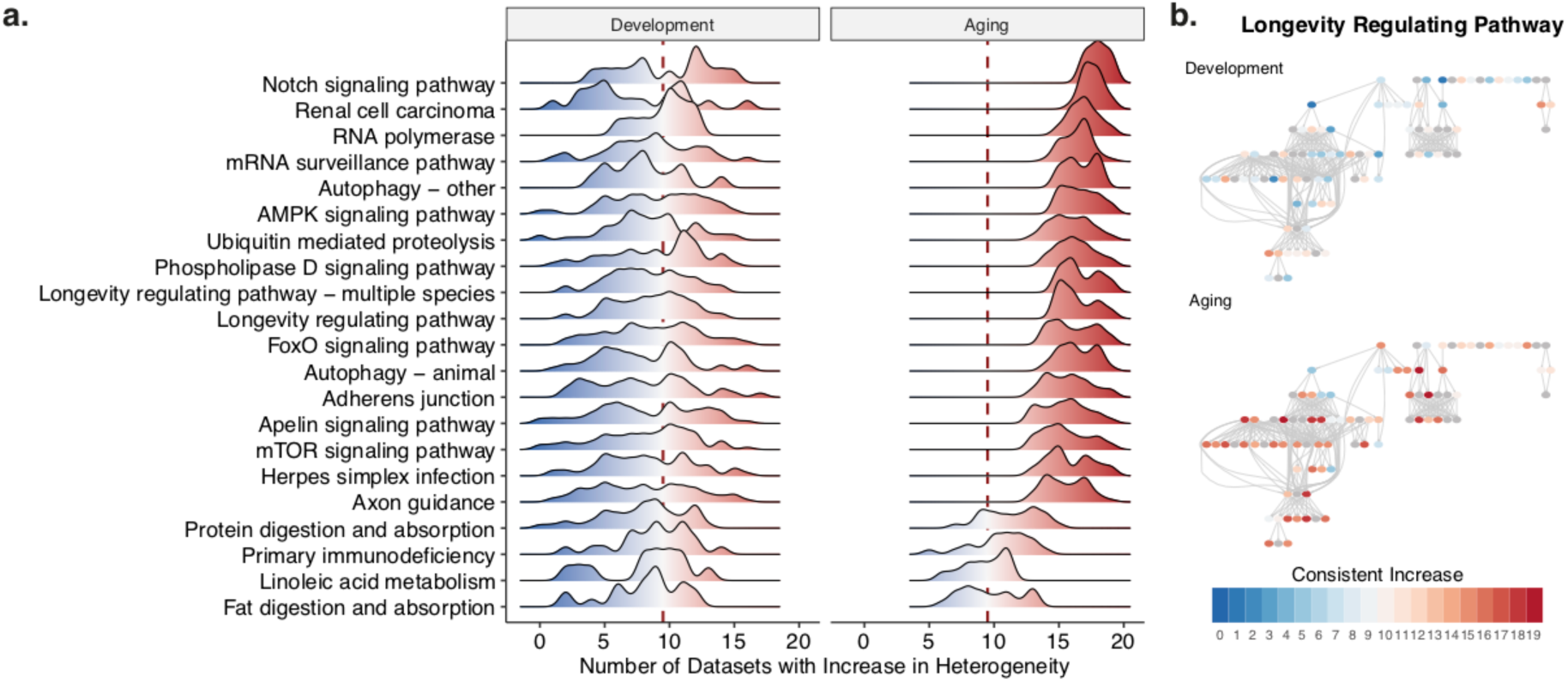
Functional analysis of consistent heterogeneity changes. (a) Distribution of consistent heterogeneity increase for the significantly enriched KEGG pathways, in development and aging. x- and y-axes show the number of datasets with a consistent increase and the density for each significant pathway, respectively. The dashed red line shows x = 9.5, which is the middle point for 19 datasets, representing no tendency to increase or decrease. Values higher than 9.5, shown with red color, indicate an increase in heterogeneity while values lower than 9.5, shown with blue color, indicate a decrease in heterogeneity and the darkness shows the consistency in change across datasets. b) The longevity regulating pathway (KEGG Pathway ID: hsa04211), exemplifying the distribution of the genes (circles), their heterogeneity across datasets (color – the same color scheme as panel (a)), and their relationship in the pathway (edges). More detailed schemes for all significant pathways with the gene names are given as SI.

The distribution of consistent heterogeneity in development and aging also showed a clear difference. The pathway scheme for the longevity regulating pathway (Figure 5b), colored based on the number of datasets with a consistent increase, shows how particular genes compare between development and aging. The visualizations for all significant pathways, including the gene names are given in the Supplementary Information. Other significantly enriched gene sets, including GO, Reactome, TF and miRNA sets, are included as Supplementary Tables S3-10. In general, while the consistent heterogeneity changes in development did not show any enrichment (except for miRNAs, see Supplementary Table S10), we detected a significant enrichment for the genes that become more heterogeneous during aging, with the exception that Disease Ontology terms were not significantly associated with consistent changes in either development or aging. The gene sets included specific categories such as autophagy and synaptic functions as well as broad functional categories such as regulation of transcription and translation processes, cytoskeleton or histone modifications. We also performed GSEA for each dataset separately and confirmed that these pathways show consistent patterns in aging (Supplementary Figs. S15-S19). There were 30 significantly enriched transcription factors, including *EGR* and *FOXO*, and 99 miRNAs (see Supplementary Table S8-9 for the full list). We also asked if the genes that become more heterogenous consistently across datasets are known aging-related genes, using the GenAge Human gene set^36^, but did not find a significant association (Supplementary Fig. S20).

It has been reported that the total number of distinct regulators of a gene (apart from its specific regulators) is correlated with gene expression noise^37^. Accordingly, we asked if the total number of transcription factors (TFs) or miRNAs regulating a gene might be related to the heterogeneity change with age (Figure 6). We calculated the correlations between the total number of regulators and the heterogeneity changes and found a mostly positive (18 / 19 for miRNA and 15 / 19 for TFs), and higher correlation between change in heterogeneity and the number of regulators in aging (*p* = 0.007 for miRNA and *p* = 0.045 TFs). We further tested the association while controlling for the expression changes in development and aging since regulation of expression changes during development could confound a relationship. However, we found that the pattern is mainly associated with the genes that show a decrease in expression during aging, irrespective of their expression during development (Supplementary Fig. S21).

**Figure 6.**
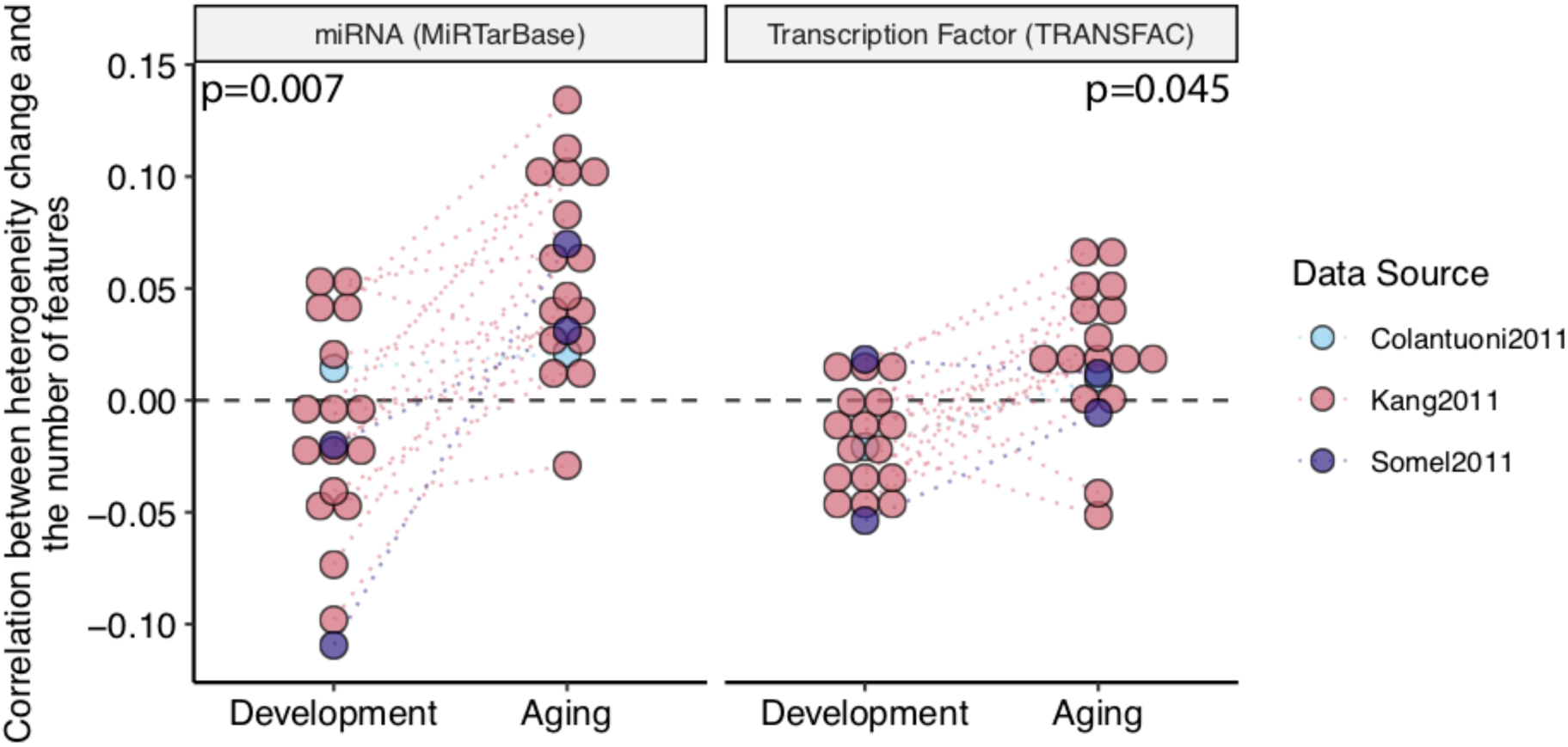
Correlation between the change in heterogeneity and number of transcriptional regulators, i.e. miRNA and transcription factors. Each point represents a dataset, and the color shows the data source. p-values are calculated using a permutation test. The dashed line at y = 0 shows zero correlation.

We further tested if genes with a consistent heterogeneity increase in aging are more central in the protein interaction network using STRING database^38^. Using multiple cutoffs and repeating the analysis, we observed a higher degree of interactions for the genes with increasing heterogeneity (Supplementary Fig. S22).

Johnson and Dong et al. previously compiled a list of traits that are age-related and have been sufficiently tested for genome-wide associations (*n* = 39)^39^. Using the genetic associations for GWAS Catalog traits, we tested if there are significantly enriched traits for the consistent changes in heterogeneity during aging (Supplementary Table S11). Although there was no significant enrichment, all these age-related terms had positive enrichment scores, *i.e.* they all tended to include genes that consistently become more heterogeneous with age during aging.

Using cell-type specific transcriptome data generated from FACS-sorted cells in mouse brain^40^, we also analyzed if there is an association between genes that become heterogeneous with age and cell-type specific genes, which could be expected if brain cell-type composition progressively varied among individuals with age. Although there was an overlap with oligodendrocytes and myelinated oligodendrocytes, there was no significant enrichment (which could be attributed to low power due to small overlap between aging and cell-type specific expression datasets) (Supplementary Fig. S23).

## Discussion

Aging is characterized by a gradual decrease in the ability to maintain homeostatic processes, which leads to functional decline, age-related diseases, and eventually to death. This age-related deterioration, however, is thought as not a result of expression changes in a few individual genes, but rather as a consequence of an age-related alteration of the whole genome, which could be a result of an accumulation of both epigenetic and genetic errors in a stochastic manner^23,41^. This stochastic nature of aging impedes the identification of conserved age-related changes in gene expression from a single dataset with a limited number of samples.

In this study, we examined 19 gene expression datasets compiled from three independent studies to identify the changes in gene expression heterogeneity with age. While all datasets have samples representing the whole lifespan, we separated postnatal development (0 to 20 years of age) and aging (20 to 98 years of age) by the age of 20, as this age is considered to be a turning-point in gene expression trajectories^13^. We implemented a regression-based method and identified genes showing a consistent change in heterogeneity with age, during development and aging separately. At the single gene level, we did not observe significant age-related heterogeneity change in most of the datasets, possibly due to insufficient statistical power due to small sample sizes and the subtle nature of the phenomenon. We hence took advantage of a meta-analysis approach and focused on consistent signals among datasets, irrespective of their effect sizes and significance. Although this approach fails to capture patterns that are specific to individual brain regions, it identifies genes that would otherwise not pass the significance threshold due to insufficient power. Furthermore, we demonstrated that our method is robust to noise and confounding effects within individual datasets.

By analyzing age-related gene expression changes, we first observed that there are more significant and more similar changes during development than aging. Additionally, genes showing significant change during aging tended to decrease in expression (Figure 1). These results can be explained by the accumulation of stochastic detrimental effects during aging, leading to a decrease in expression levels^2^. Our initial analysis of gene expression changes suggested a higher heterogeneity between aging datasets.

We next focused on age-related heterogeneity change between individuals and found a significant increase in age-related heterogeneity during aging, compared to development. Notably, increased heterogeneity is not limited to individual brain regions, but a consistent pattern across different regions during aging. We found that age-related heterogeneity change is more consistent among aging datasets, which may reflect an underlying systemic mechanism. Further, a larger number of genes showed more significant heterogeneity changes during aging than in development, and the majority of these genes tended to have more heterogeneous expression.

It was previously proposed that somatic mutation accumulations^2,41–43^ and epigenetic regulations^44^ might be associated with transcriptome instability. While Enge et al. and Lodato et al. suggested that genome-wide substitutions in single cells are not so common as to influence genome stability and cause transcriptional heterogeneity at the cellular level^23,45^, epigenetic mechanisms may be relevant. Although we cannot test age-related somatic mutation accumulation and epigenetic regulation in this study, an alternative mechanism might be related to transcriptional regulation, which is considered to be inherently stochastic^46^. Several studies demonstrated that variation in gene expression is positively correlated with the number of TFs controlling gene’s regulation^37^. We also found that genes with a higher number of regulators and a decrease in expression during aging become more heterogeneous. Further, significantly enriched TFs include early growth response (*EGF*), known to be regulating the expression of many genes involved in synaptic homeostasis and plasticity, and *FOXO* TFs, which regulate stress resistance, metabolism, cell cycle arrest and apoptosis. Together with these studies, our results support that transcriptional regulation may be associated with age-related heterogeneity increase during aging and may have important functional consequences in brain aging.

We next confirmed that observed increase in heterogeneity was not a result of low statistical power (Supplementary Fig. S1) or a technical artifact (Figure 3b, Supplementary Figs. S24-S25). Specifically, we tested whether increased heterogeneity during aging can be a result of the mean-variance relationship, but we found no significant effect that can confound our results. In fact, the mean-variance relationship in development and aging showed opposing profiles. We further analyzed this by grouping genes based on their expression in development and aging (Supplementary Fig. S24). The genes that decrease in expression both in development and aging showed the most opposing profiles in terms of the mean-variance relationship, which could suggest that the decrease in development are more coordinated and well-regulated whereas the decrease in aging occurs due to stochastic errors. Another potential confounder is the post-mortem interval (PMI), which is the time between death and sample collection. Since we do not have this data for all datasets we analyzed, we could not account for PMI in our model. However, using the list of genes previously suggested as associated with PMI^47^, we checked if the consistency among aging datasets could be driven by PMI. Only 2 PMI-associated genes were among the 147 that become consistently heterogeneous, and the distribution also suggested there is no significant relationship (Supplementary Fig. S25). We also confirmed that the increase in heterogeneity is not caused by outlier samples in datasets (Supplementary Fig. S26) or by the confound of sex with age (Supplementary Fig. S27).

One important limitation of our study is that we analyze microarray-based data. Since gene expression levels measured by microarray do not reflect an absolute abundance of mRNAs, but rather are relative expression levels, we were only able to examine relative changes in gene expression. A recent study analyzing single-cell RNA sequencing data from the aging *Drosophila* brain identified an age-related decline in total mRNA abundance^48^. It is also suggested that, in microarray studies, genes with lower expression levels tend to have higher variance^49^. In this context, whether the change in heterogeneity is a result of the total mRNA decay is an important question. As an attempt to see if the age-related increase in heterogeneity is dependent on the technology used to generate data, we repeated the initial analysis using RNA sequencing data for the human brain, generated by GTEx Consortium ^50^ (Supplementary Figs. S28-30). Nine out of thirteen datasets displayed more increase than decrease in heterogeneity during aging, consistent with 18/19 microarray datasets, while the remaining four datasets showed the opposite pattern (BA24, cerebellar hemisphere, cerebellum and substantia nigra). Unlike what we observed for the microarray datasets, the change in expression levels and heterogeneity were strongly positively correlated (Supplementary Fig. S30). Unfortunately, average expression levels and variation levels in RNA sequencing is challenging to disentangle. Thus, the biological relevance of the relationship between the age-related change in expression levels and expression heterogeneity still awaits to be studied through novel experimental and computational approaches. Nevertheless, RNA sequencing analysis also suggests an overall increase in age-related heterogeneity increase.

Another limitation is related to use of bulk RNA expression datasets, where each value is an average for the tissue. While it is important to note that our results indicate increased heterogeneity between individuals rather than cells, the fact that the brain is composed of different cell types raises the question if increased heterogeneity may be a result of changes in brain cell-type proportions. To explore the association between heterogeneity and cell-type specific genes, we used FACS-sorted cell type specific transcriptome dataset from mouse brain^40^. We only had nine genes that have consistent heterogeneity increase and are specific to one cell-type. Eight out of nine were highly expressed in oligodendrocytes, which is consistent with the results reported in our earlier work^24^. However, we did not observe any significant association between cell-type specific genes and heterogeneity (Supplementary Fig. S23).

Gene set enrichment analysis of the genes with increased heterogeneity with age revealed a set of significantly enriched pathways that are known to modulate aging, including longevity regulating pathway, autophagy, mTOR signaling pathway (Figure 5a). Furthermore, GO terms shared among these genes include some previously identified common pathways in aging and age-related diseases (Supplementary Figs. S16-18). We have also tested if these genes are associated with age-related diseases through GWAS, and although not significant, we found a positive association with all age-related traits defined in Johnson and Dong et al. Overall, these results indicate the effect of heterogeneity on pathways that modulate aging and may reflect the significance of increased heterogeneity in aging. Importantly, we identified genes that are enriched in terms related to neuronal and synaptic functions, such as axon guidance, neuron to neuron synapse, postsynaptic specialization, which may reflect the role of increased heterogeneity in synaptic dysfunction observed in the mammalian brain, which is considered to be a major factor in age-related cognitive decline^51^. We also observed genes that become more heterogeneous with age consistently across datasets are more central (*i.e.* have a higher number of interactions) in a protein-protein interaction network (Supplementary Fig. S22). Although this could mean the effect of heterogeneity could be even more critical because it affects hub genes, another explanation is research bias that these genes are studied more than others.

In summary, by performing a meta-analysis of transcriptome data from diverse brain regions we found a significant increase in gene expression heterogeneity during aging, compared to development. Increased heterogeneity was a consistent pattern among diverse brain regions in aging, while no significant consistency was observed across development datasets. Our results support the view of aging as a result of stochastic molecular alterations, whilst development has a higher degree of gene expression regulation. We also found that genes showing a consistent increase in heterogeneity during aging are involved in pathways important for aging and neuronal function. Therefore, our results demonstrate that increased heterogeneity is one of the characteristics of brain aging and is unlikely to be only driven by the passage of time starting from developmental stages.

## Methods

### Dataset collection

In this study, we performed re-analysis of publicly available transcriptome datasets to test age-related change in gene expression heterogeneity. All data collection in these previous studies were performed in accordance with relevant guidelines, regulations and approved experimental protocols, including informed consents for the use of samples for research from all donors or their next of kin.

#### Microarray datasets

Raw data used in this study were retrieved from the NCBI Gene Expression Omnibus (GEO) from three different sources (Supplementary Table S1). All three datasets consist of human brain gene expression data generated on microarray platforms. In total, we obtained 1017 samples from 298 individuals, spanning the whole lifespan with ages ranging from 0 to 98 years (Supplementary Fig. S1).

#### RNA sequencing dataset

We used the transcriptome data generated by the GTEx Consortium (v6p)^50^. We only used the samples with a death circumstance of 1 (violent and fast deaths due to an accident) and 2 (fast death of natural causes) on the Hardy Scale excluding individuals who died of illnesses. As we focus only on the brain, we used all 13 brain tissue data in GTEx. We thus analyzed 623 samples obtained from 99 individuals.

#### Separating datasets into development and aging datasets

To differentiate changes in gene expression heterogeneity during aging from those during development, we used the age of 20 to separate pre-adulthood from adulthood. It was shown that the age of 20 corresponds to the first age of reproduction in human societies^52^. Structural changes after the age of 20 in the human brain were previously linked to age-related phenotypes, specifically neuronal shrinkage and a decline in total length of myelinated fibers^3^. Earlier studies examining age-related gene expression changes in different brain regions also showed a global change in gene expression patterns after the age of 20^11,13,53^. Thus, consistent with these studies, we separated datasets using the age of 20 into development (0 to 20 years of age, *n* = 441) and aging (20 to 98 years of age, *n* = 569).

### Preprocessing

#### Microarray datasets

RMA correction (using the ‘oligo’ library in R)^54^ and log2 transformation were applied to Somel2011 and Kang2011 datasets. For the Colantuoni2011 dataset, as there was no public R package to analyze the raw data, we used the preprocessed data deposited in GEO, which had been loess normalized by the authors. We quantile normalized all datasets using the ‘preprocessCore’ library in R^55^. Notably, our analysis focused on consistent patterns across datasets, instead of considering significant changes within individual datasets. Since we don’t expect random confounding factors to be shared among datasets, we used quantile normalization to minimize the effects of confounders, and we treated consistent results as potentially a biological signal. We also applied an additional correction procedure for Somel2011 datasets, in which there was a batch effect influencing the expression levels, as follows: for each probeset (1) calculate mean expression (M), (2) scale each batch separately (to mean = 0, standard deviation = 1), (3) add M to each value. We excluded outliers given in Supplementary Table S1, through a visual inspection of the first two principal components for the probeset expression levels (same as in Dönertaş, Fuentealba Valenzuela, Partridge, & Thornton, 2018; Dönertaş et al., 2017). We mapped probeset IDs to Ensembl gene IDs 1) using the Ensembl database, through the ‘biomaRt’ library ^57^ in R for the Somel2011 dataset, 2) using the GPL file deposited in GEO for Kang2011, as probeset IDs for this dataset were not complete in Ensembl, and 3) using the Entrez gene IDs in the GPL file deposited in GEO for the Colantuoni2011 dataset and converting them into Ensembl gene IDs using the Ensemble database, through the “biomaRt” library in R. Lastly, we scaled expression levels for genes (to mean = 0, standard deviation = 1) using the ‘scale’ function in R. Age values of individuals in each dataset were converted to the fourth root of age (in days) to have a linear relationship between age and expression both in development and aging. However, we repeated the analysis using different age scales and confirmed that the results were quantitatively similar (Supplementary Fig. S3).

#### RNA sequencing dataset

The genes with median RPKM value of 0 were excluded from the dataset. The RPKM values provided in the GTEx data were log2 transformed and quantile-normalized. Similar to the microarray data, we excluded the outliers based on the visual inspection of the first and second principal components (Supplementary Table S1). In GTEx, ages are given as 10 year intervals. We therefore used the middle point of these age intervals to represent that individual’s age.

### Age-related expression change

We used linear regression to assess the relationship between age and gene expression. The model used in the analysis is:

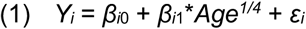

where Y_i_ is the scaled log2 expression level for the i^th^ gene, *β*_*i*_0 is the intercept, *β*_*i*_1 is the slope, and ε_i_ is the residual. We performed the analysis for each dataset (development and aging datasets separately) and considered the *β*_*1*_ value as a measure of change in expression. *p*-values obtained from the model were corrected for multiple testing according to Benjamini and Hochberg procedure^58^ by using ‘p.adjust’ function in R.

### Age-related heterogeneity change

In order to quantify the age-related change in gene expression heterogeneity, we calculated Spearman’s correlation coefficient (*ρ*). The correlations were calculated between the absolute values of residuals obtained from equation (1) and the fourth root of individual age. We regarded the absolute values of residuals as a measure of heterogeneity. Therefore, high positive correlation coefficients suggest that heterogeneity increases with age, whereas strong negative correlation implies heterogeneity decreases with age. *p*-values were calculated from the correlation analysis and corrected for multiple testing with the Benjamini and Hochberg method using the ‘p.adjust’ function in R. To compare heterogeneity changes in aging and development, we employed paired Wilcoxon test (‘wilcox.test’ in the R ‘stats’ package) in which we compared median heterogeneity changes in aging and development dataset pairs.

### Principal Component Analysis

We conducted principal component analysis on both age-related changes in expression (*β*) and heterogeneity (*ρ*) We followed a similar procedure for both analyses, in which we used the ‘prcomp’ function in R. The analysis was performed on a matrix containing *β* values (for the change in expression level) and *ρ* values (for the change in heterogeneity), for 11,137 commonly expressed genes for all 38 development and aging datasets. In each dataset, the estimates of expression change (*β*) or heterogeneity change (*ρ*) values were scaled for each dataset before calculating principal components.

### Permutation test

We performed a permutation test, taking into account the non-independence of samples across the Somel2011 and Kang2011 datasets, due to the fact that these datasets include multiple samples from the same individuals for different brain regions. We first randomly permuted ages among individuals, not samples, for 1,000 times in each data source, using the ‘sample’ function in R. Next, we assigned ages of individuals to corresponding samples and calculated age-related expression and heterogeneity change for each dataset, corresponding to different brain regions. For the tests related to the changes in gene expression with age, we used a linear model between gene expression levels and the randomized ages. In contrast, for the tests related to the changes in heterogeneity with age, we measured the correlation between the randomized ages and the absolute value of residuals from the linear model that is between expression levels and non-randomized ages for each gene. In this way, we preserved the relationship between age and expression, and we were able to ensure that our regression model was viable for calculating age-related heterogeneity change. Using expression and heterogeneity change estimates calculated using permuted ages, we tested (a) if the correlation of expression (and heterogeneity) change in aging is higher than in development datasets; (b) if the correlations of expression (and heterogeneity) change in development and in aging datasets are significantly higher than null expectation; (c) if the number of genes showing significant change in expression (and heterogeneity) is significantly higher in aging than in development datasets; (d) if the overall increase in age-related heterogeneity during aging is significantly higher than development; (e) if the observed consistency in heterogeneity increase is significantly different from expected. All tests using permuted ages were performed one-tailed. We also demonstrate that our permutation strategy is more stringent than random permutations in Supplementary Figure S10, giving the distributions calculated using both dependent permutations and random permutations.

To test the overall correlation within development or aging datasets for the changes in expression (*β*) and heterogeneity (*ρ*), we calculated median correlations among independent three subsets of datasets (one Kang2011, one Somel2011 and the Colantuoni2011 dataset), taking the median value calculated for each possible combination of independent subsets (16 × 2 × 1 = 32 combinations). Using 1,000 permutations of individuals’ ages, we generated an expected distribution for the median correlation coefficient for triples and compared these with the observed values, asking how many times we observe a higher value. We used this approach to calculate expected median correlation among development (and aging) datasets, because the number of independent pairwise comparisons are outnumbered by the number of dependent pairwise comparisons, causing low statistical power.

To further test the significance of the difference between correlations among development and aging datasets, we calculated the median difference in correlations between aging and development datasets for each permutation. We next constructed the null distribution of 1,000 median differences and calculated empirical *p*-values compering the observed differences with these null distributions. Next, to test the significance of the difference in the number of significantly changing genes between development and aging, we calculated the difference in the number of genes showing significant change between development and aging datasets for each permutation. Empirical *p*-values were computed according to observed differences. Likewise, to test if the overall increase in age-related heterogeneity during aging is significant compared to development, we computed median differences between median heterogeneity change values of each aging and development dataset, for each permutation, followed by an empirical *p*-value calculation to answer if the aging datasets have a higher increase in age-related heterogeneity.

### Expected heterogeneity consistency

Expected consistency in heterogeneity change was calculated from heterogeneity change values (*ρ*) measured using permuted ages. For each permutation, we first calculated the total number of genes showing consistent heterogeneity increase for N number of datasets (N = 0, …, 19). To test if observed consistency significantly differed from the expected, we compared observed consistency values to the distribution of expected numbers, by performing a one-sided test for the consistency in N number of datasets, N = 1, …, 19.

### Clustering

We used the k-means algorithm (‘kmeans’ function in R) to cluster genes showing consistent heterogeneity change (n=147) according to their heterogeneity profiles. We first took the subset of the heterogeneity levels (absolute value of the residuals from equation (1)) to include only the genes that show a consistent increase with age and then scaled the heterogeneity levels to the same mean and standard deviation. Since the number of samples in each dataset is different, just running k-means on the combined dataset would not equally represent all datasets. Thus, we first calculated the spline curves for scaled heterogeneity levels for each gene in each dataset (using the ‘smooth.spline’ function in R, with three degrees of freedom). We interpolated at 11 (the smallest sample size) equally distant age points within each dataset. Then we used the combined interpolated values to run the k-means algorithm with k = 8, a liberal choice, given the total number of genes being 147.

To test association of the clusters with Alzheimer’s Disease, we retrieved overall AD association scores of the 147 consistent genes (*n* = 40) from the Open Targets Platform^59^.

### Functional Analysis

We used the “clusterProfiler” package in R to run Gene Set Enrichment Analysis, using Gene Ontology (GO) Biological Process (BP), GO Molecular Function (MF), GO Cellular Compartment (CC), Reactome, Disease Ontology (DO), and KEGG Pathways. We performed GSEA on all gene sets with a size between 5 and 500, and we corrected the resulting *p*-values with the Benjamini and Hochberg correction method. To test if the genes with a consistent increase or decrease in their expression are associated with specific functions, we used the number of datasets with a consistent increase to run GSEA. Since we are running GSEA using number of datasets showing consistency, our data includes many ties, potentially making the ranking ambiguous and non-robust. In order to assess how robust our results are, we ran GSEA 1,000 times on the same data and counted how many times we observed the same set of KEGG pathways as significant (Supplementary Table S3). The lowest number among the pathways with a significant positive enrichment score was 962 out of 1,000 (Phospholipase D signaling pathway). Moreover, we repeated the same analysis using the heterogeneity change levels (*ρ*), instead of using the number of datasets with a consistent change, for each dataset to confirm the gene sets are indeed associated with the increase or decrease in heterogeneity (Supplementary Figs. S15-S19). We visualized the KEGG pathways using ‘KEGGgraph’ library in R and colored the genes by the number of datasets that show an increase.

We also performed an enrichment analysis of the transcription factors and miRNA to test if specific TFs or miRNAs regulate the genes that become more heterogeneous consistently. We collected gene-regulator association information using the Harmonizome database^60^, “MiRTarBase microRNA Targets” (12086 genes, 596 miRNAs) and “TRANSFAC Curated Transcription Factor Targets” (13216 genes, 201 TFs) sets. We used the ‘fgsea’ package in R, which allows GSEA on a custom gene set. We tested the association for each regulator with at least 10 and at most 500 targets. Moreover, we tested if the number of regulators is associated with the change in heterogeneity. We first calculated the correlation between heterogeneity change with age (or the number of datasets with an increase in expression heterogeneity) and the number of TFs or miRNAs regulating that gene, for aging and development separately. We repeated the analysis while accounting for the direction of expression changes in these periods (*i.e.* separating genes into down-down, down-up, up-down, and up-up categories based on their expression in development and aging, Supplementary Fig. S21). To test the difference in the correlations between aging and development, we used 1,000 random permutations of the number of TFs. For each permutation, we randomized the number of TFs and calculated the correlation between heterogeneity change (or the number of datasets with an increase in heterogeneity) and the randomized numbers. We then calculated the percentage of datasets where aging has a higher correlation than development. Using the distribution of percentages, we tested if the observed value is expected by chance.

### Protein-protein interaction network analysis

We downloaded all human protein interaction data from the STRING database (v11)^38^. Ensembl Peptide IDs are mapped to Ensembl Gene IDs using the “biomaRt” package in R. Here we aimed to test whether genes showing consistent increase in heterogeneity have a different number of interactors than other genes. For this we calculated the degree distributions for the genes that become consistently more heterogeneous with age and all remaining genes using different cutoffs for interaction confidence scores. In order to calculate the significance of the difference, we i) calculated the number of interactors (degree) for each gene, ii) for 10,000 times, randomly sampled k genes from all interactome data (k = number of genes that become heterogeneous with age across all datasets and have interaction information in STRING database, after filtering for a cutoff), iii) calculated the median of degree for each sample. We then calculated an empirical *p*-value by asking how many of these 10,000 samples we see a median degree that is equivalent to or higher than our original value. The number of genes and interactions after each cutoff are given in Supplementary Figure S22.

### Cell-type specificity analysis

Using FACS-sorted cell-type specific transcriptome data from the mouse brain^40^, we checked if there is any overlap between genes that become heterogeneous with age and cell-type specific genes. We downloaded the dataset from the GEO database (GSE9566) and preprocessed it by performing: i) RMA correction using the ‘affy’ package in R^61^, ii) log2 transformation, iii) quantile normalization using the ‘preprocessCore’ package in R^55^, iv) mapping probeset IDs to first mouse genes, and then human genes. We only included genes that have one to one orthologs in humans, after filtering out probesets that map to multiple genes. We defined cell-type specific genes by calculating the effect size (Cohen’s D) for each gene and cell type and identifying genes that have an effect size higher than or equal to 2 as specific to that cell type. At this cutoff, there was no overlap between cell type specific gene lists. To test for association between heterogeneity and cell type specificity, we used the Fisher’s exact test using the R ‘fisher.test’ function.

## Supporting information

Supplementary Material - Pathway results

## Code Availability

All analysis was performed using R and the code to calculate heterogeneity changes with age is available as an R package ‘hetAge’, documented at https://mdonertas.github.io/hetAge/. “ggplot2”^62^ and “ggpubr”^63^ R libraries were used for the visualization.

## Data availability

We performed re-analysis of the raw data that we downloaded from the GEO database (GSE30272, GSE25219, GSE22569, GSE18069) and GTEx data portal. All results generated in this study are available as Supplementary Tables and all summary statistics are available in the BioStudies database (http://www.ebi.ac.uk/biostudies) under accession number S-BSST273.

## Author Contributions

H.M.D. conceived and designed the study with the contributions from M.S., and J.M.T.. U.I. and H.M.D. analyzed the data. U.I. and H.M.D. interpreted the results and wrote the manuscript with the contributions from M.S. and J.M.T. All authors read, revised and approved the final version of this manuscript.

## Acknowledgements

The authors thank Hamit Izgi, Matias Fuentealba Valenzuela, Dr. Daniel K. Fabian, and Prof Linda Partridge for helpful discussions. H.M.D. is a member of Darwin College, University of Cambridge.

