## Supplementary Material for "Gene expression heterogeneity during brain development and aging: temporal changes and functional consequences"

**Supplementary Tables**

**Table S1:** The list of human brain gene expression datasets analyzed in the study.

**Table S2**: Genes showing consistent heterogeneity increase (*n* = 147) across all 19 datasets and the trajectory clusters they belong to.

**Table S3:** KEGG Pathways that are significantly associated (FDR corrected $p \leq 0.1$) with consistent heterogeneity change in aging.

**Table S4:** Gene Ontology Biological Process Categories that are significantly associated (FDR corrected $p \leq 0.1$) with consistent heterogeneity change in aging.

**Table S5:** Gene Ontology Molecular Function Categories that are significantly associated (FDR corrected $p \leq 0.1$) with consistent heterogeneity change in aging.

**Table S6:** Gene Ontology Cellular Component Categories that are significantly associated (FDR corrected $p \leq 0.1$) with consistent heterogeneity change in aging.

**Table S7:** Reactome Pathways that are significantly associated (FDR corrected $p \leq 0.1$) with consistent heterogeneity change in aging.

**Table S8:** Transcription factors that are significantly associated (FDR corrected $p \leq 0.1$) with consistent heterogeneity change in aging.

**Table S9:** miRNAs that are significantly associated (FDR corrected $p \leq 0.1$) with consistent heterogeneity change in aging.

**Table S10:** miRNAs that are significantly associated (FDR corrected $p \leq 0.1$) with consistent heterogeneity change in development.

**Table S11:** GSEA results showing the association between the aging-related traits (Johnson et al. 2015) and consistent heterogeneity change in aging.

### Supplementary Figures

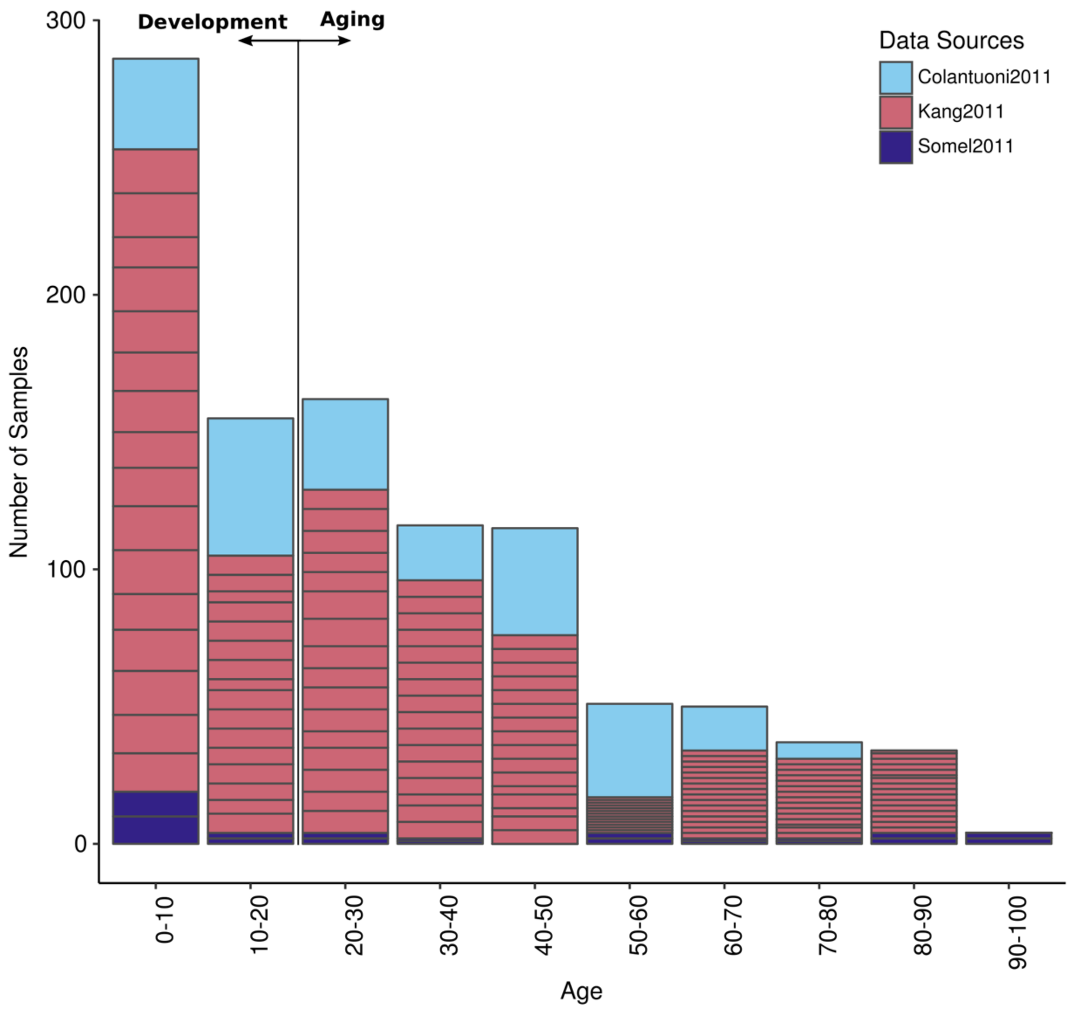


**Figure S1:** The distribution of ages for the samples used in the analysis. x- and y-axes show age intervals and number of samples included, respectively. The horizontal line reflects the separation point of development and aging (age of 20). Data sources are indicated with different colors.

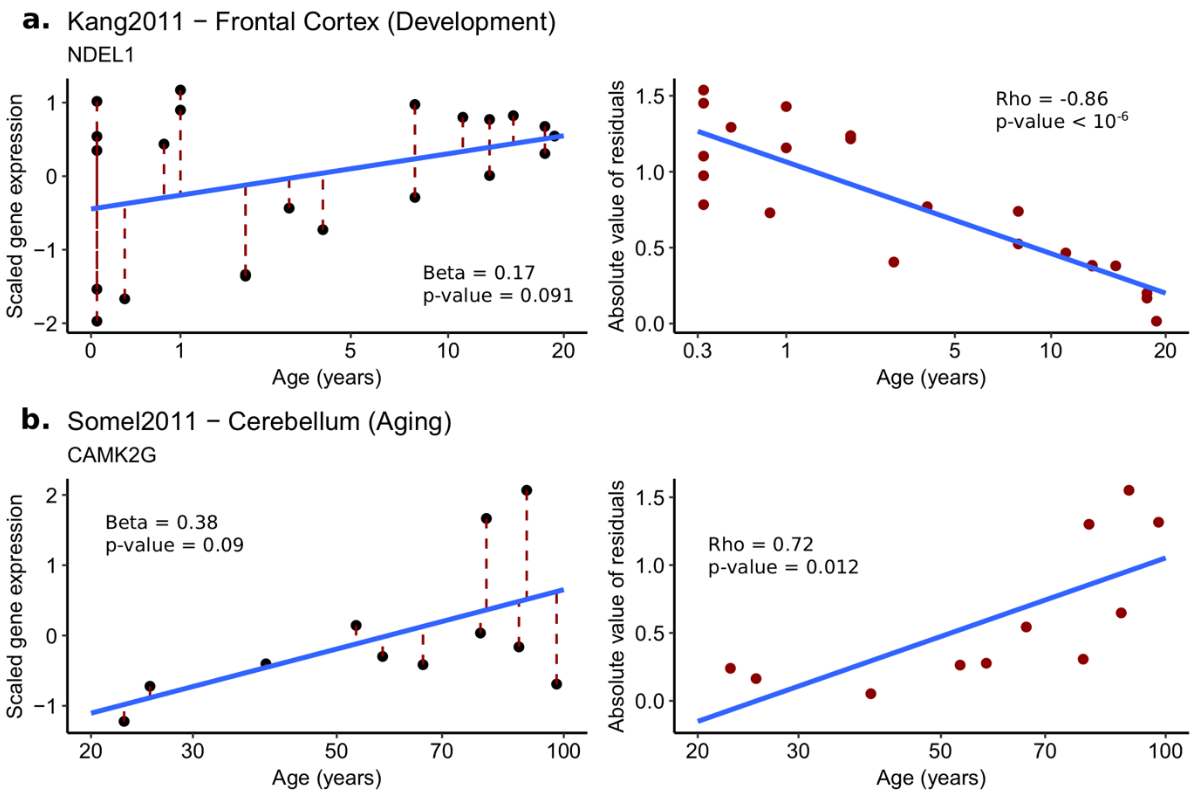


**Figure S2:** Demonstration of the method used to calculate age-related changes in expression level and heterogeneity of two genes during development (a) and aging (b). *Left panels:* Age-related expression change is quantified by a linear model using scaled gene expression values (y-axis) and the fourth root of ages (x-axis). Beta values and raw *p*-values are indicated on the figure. The blue line is drawn based on the linear model, and the red dotted lines represent residuals. Right panels: corresponding scatterplots for the absolute values of residuals (y-axis) and fourth root of age (x-axis). Rho and raw *p*-values are calculated based on Spearman's correlation test. The blue line is drawn based on a linear model between the absolute value of residuals and the fourth root of age, only for demonstration purposes.

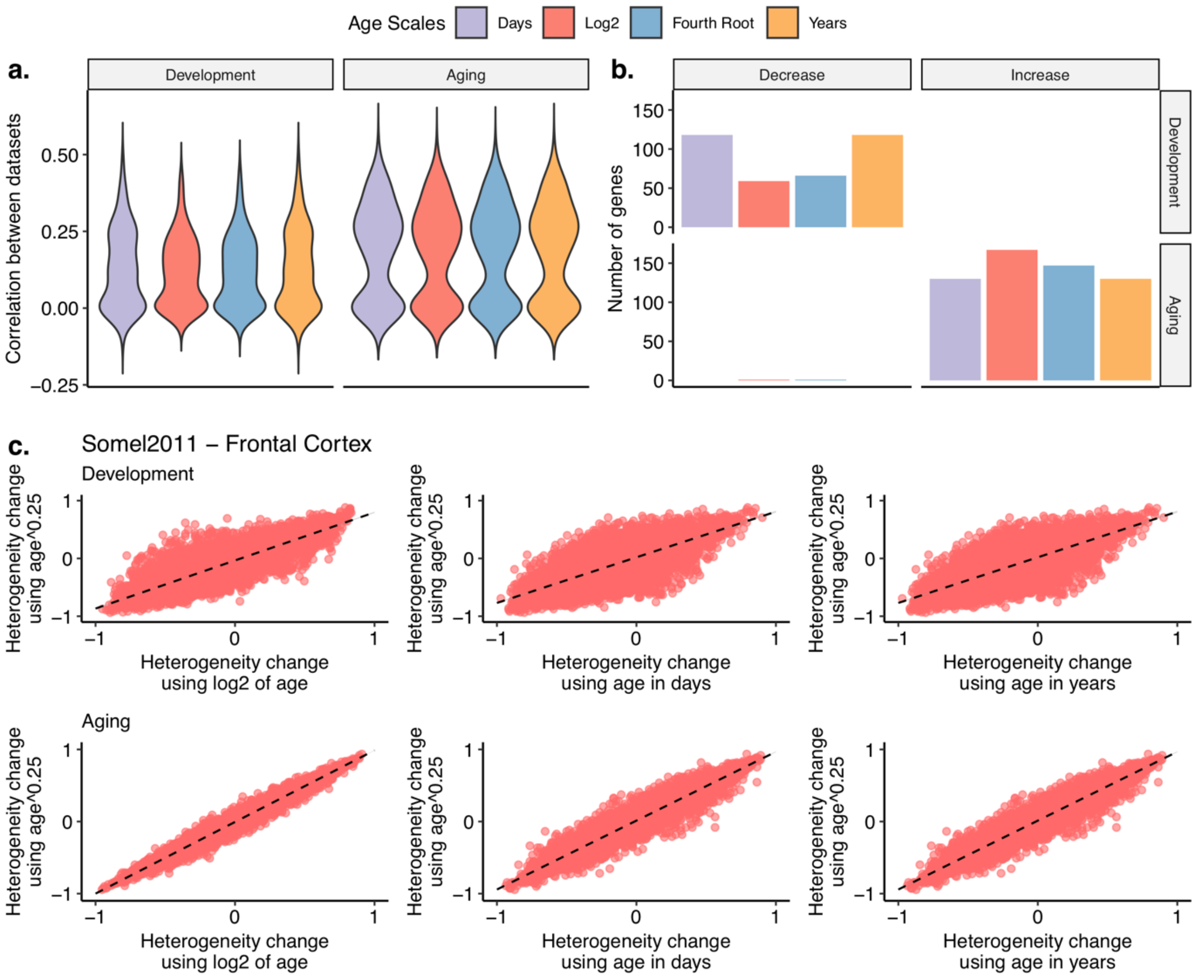


**Figure S3:** Different age scales yields consistent results. We performed downstream analyses using age in days, age in years, log2 of age in days, and fourth root of age in days. (a) Distribution of correlation coefficients between age-related heterogeneity changes across development and aging datasets (y-axis), using different age scales (x-axis). (b) The number of genes showing consistent heterogeneity change across all 19 datasets (y-axis), using different age scales. (c) Scatterplots of age-related heterogeneity change values of 11,137 genes from one example dataset (Somel2011_PFC) calculated using the fourth root of age in days (y-axis) and different scales including log2 of age in days, age in days and age in years (x-axis) during development (upper panel) and aging (lower panel).

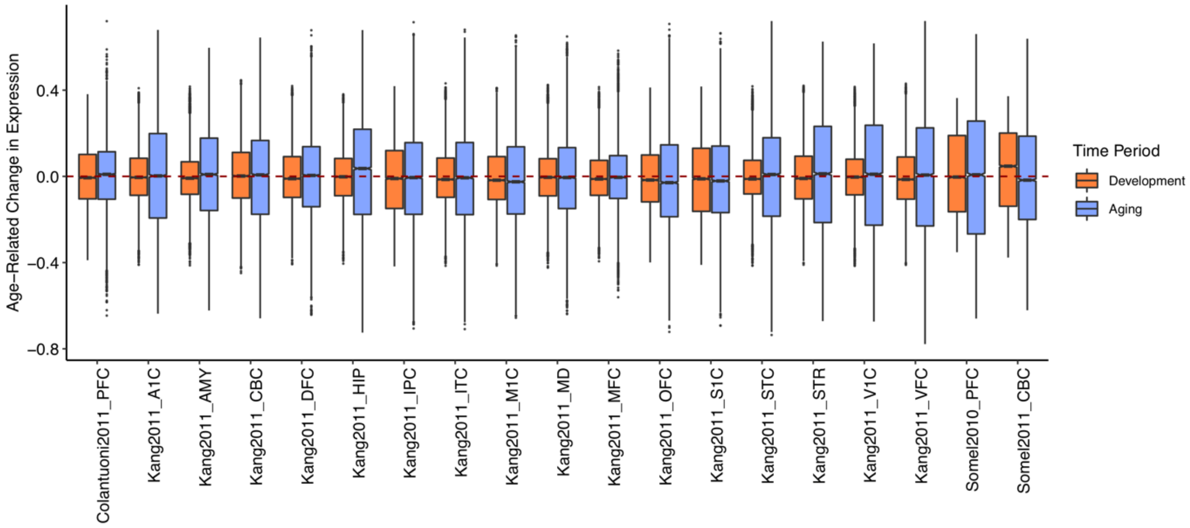


**Figure S4:** Distribution of age-related change in expression values (Beta values) for each dataset (x-axis) during development and aging. Beta values (y-axis) were computed by linear models using scaled expression values and the fourth root of age for each gene in each dataset during development (orange) and aging (blue) separately (see Figure S2).

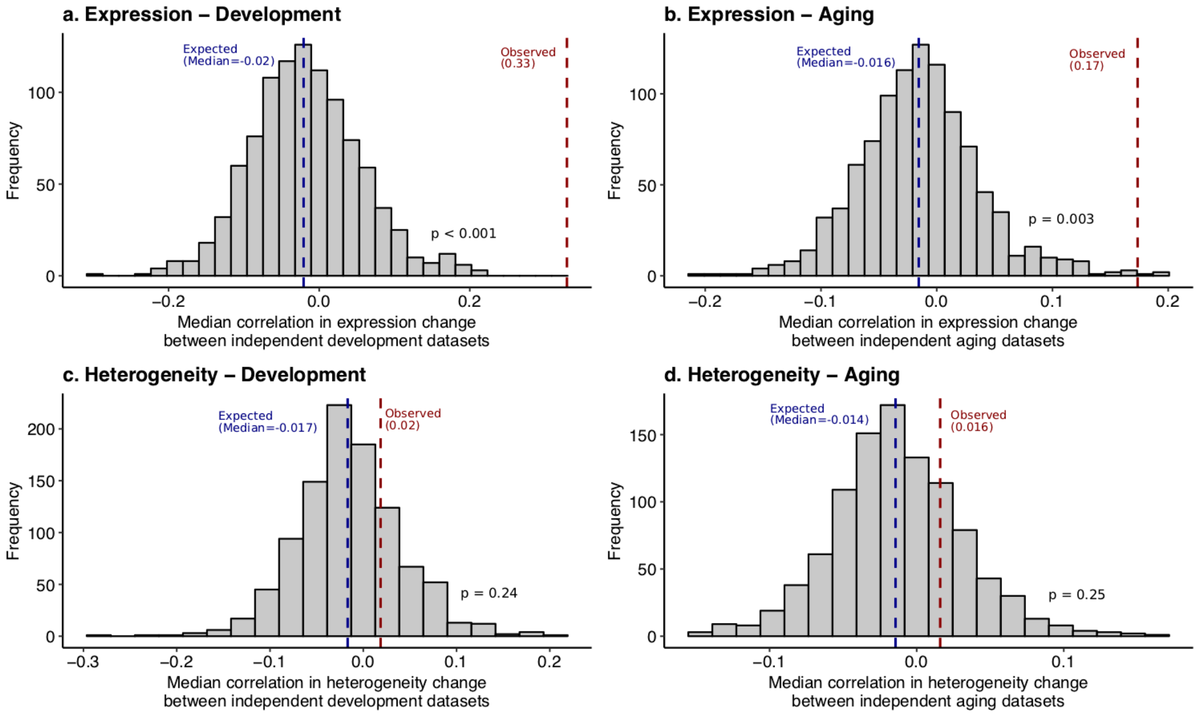


**Figure S5:** Permutation test results for dataset correlations of expression and heterogeneity change during development and aging. We constructed the distributions by calculating median correlation between all possible three independent datasets from three data sources for all permutations. Similarly, the observed values were calculated as a median value for all three independent dataset combinations from three data sources (i.e. the median value of: Median Correlation(Kang2011_A1C, Somel2011_PFC, Colantuoni2011_PFC), Median Correlation(Kang2011_AMY, Somel2011_CBC, Colantuoni2011_PFC), …). (a, b) Permutation test results for the significance of observed correlation in gene expression change during development (a) and aging (b). (c, d) Permutation test results for the significance of observed correlation in heterogeneity change during development (c) and aging (d).

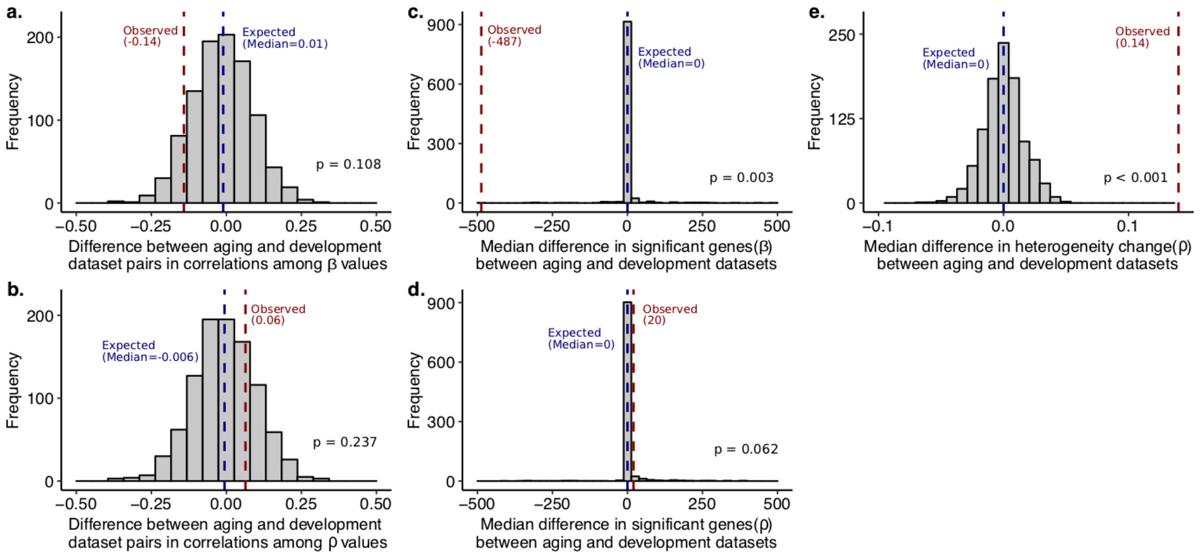


**Figure S6:** (a, b) Permutation test results for differences between dataset correlations during development and aging. Distributions were constructed using the median differences between correlations among aging datasets and development datasets for 1000 permutations, i.e. negative values imply higher correlation among development datasets compared to aging datasets, while positive values indicate higher correlation during aging. (a) Distribution of median differences between correlations among age-related expression changes (β values) among aging datasets and development datasets. (b) The same distribution for heterogeneity changes (ρ values). (c, d) Permutation test results for the median difference in the number of genes showing significant change between aging and development datasets, in terms of (c) expression change and (d) heterogeneity change. For each permutation, *p*-values were corrected for multiple testing using FDR method. (e) Permutation test result for the median heterogeneity change differences between aging and development datasets. Associated *p*-values are from one-tailed tests.

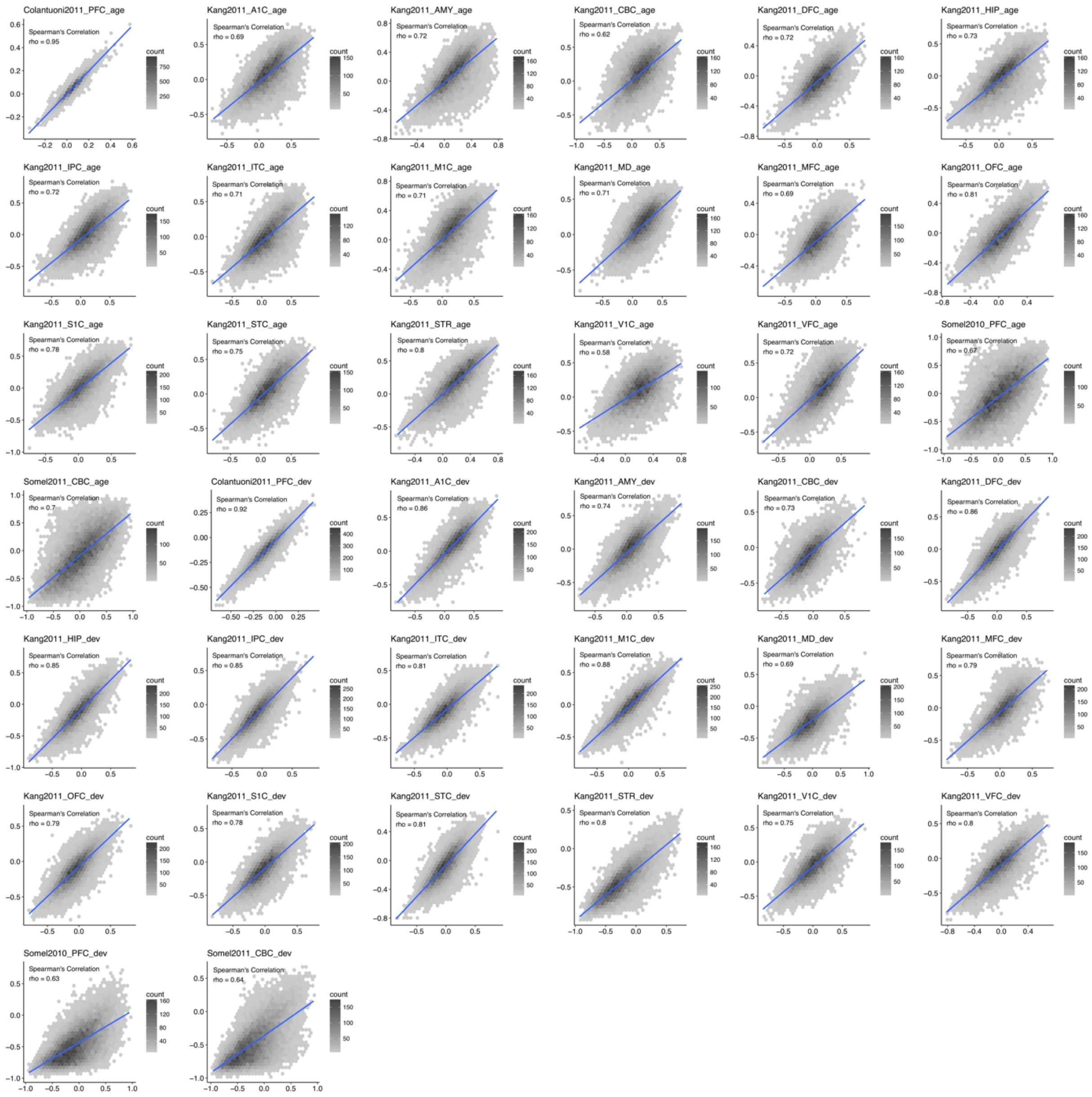


**Figure S7:** Hexagonal density maps for the changes in heterogeneity based on the residuals from a linear model (x-axis) and loess regressions (y-axis). The color intensity shows the number of genes in that hexagonal bin, where darker color shows more genes. The blue line is drawn using linear regression, for demonstration purposes. The correlations between the values are calculated using Spearman's correlation test.

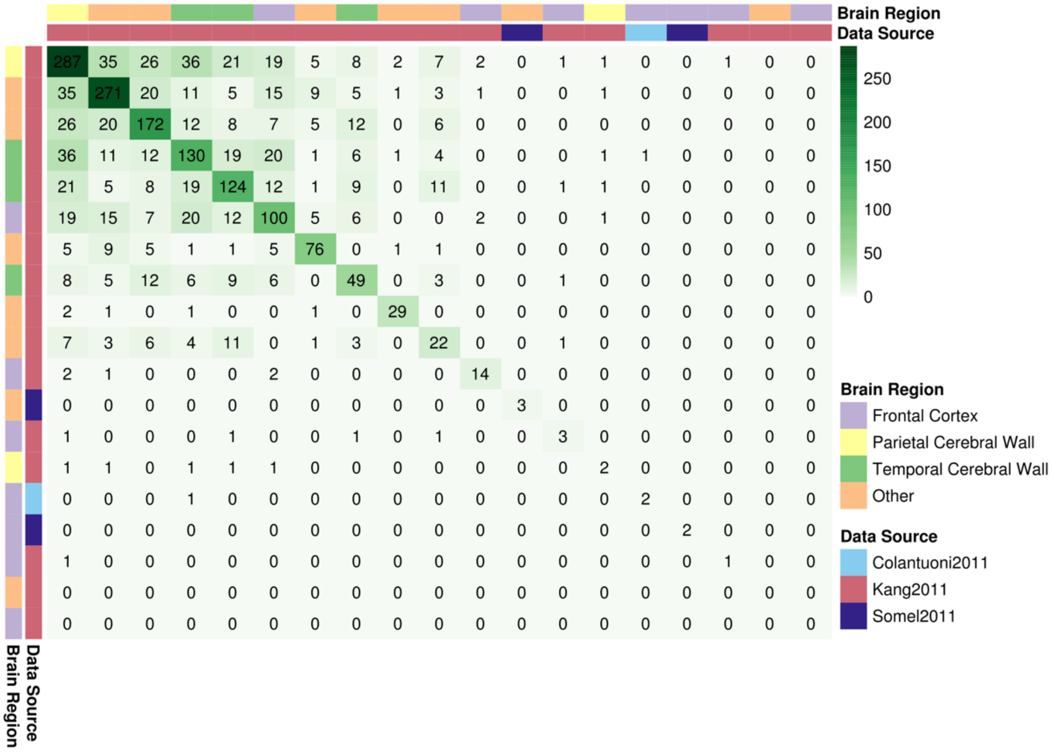


**Figure S8:** Overlaps between datasets for the genes showing a significant increase in heterogeneity. *p*-values are computed using Spearman correlation test between the absolute value of residuals and fourth root of ages, followed by FDR correction. The color intensity of cells reflects an increased number of overlapping genes among two corresponding datasets, while the numbers show the exact numbers.

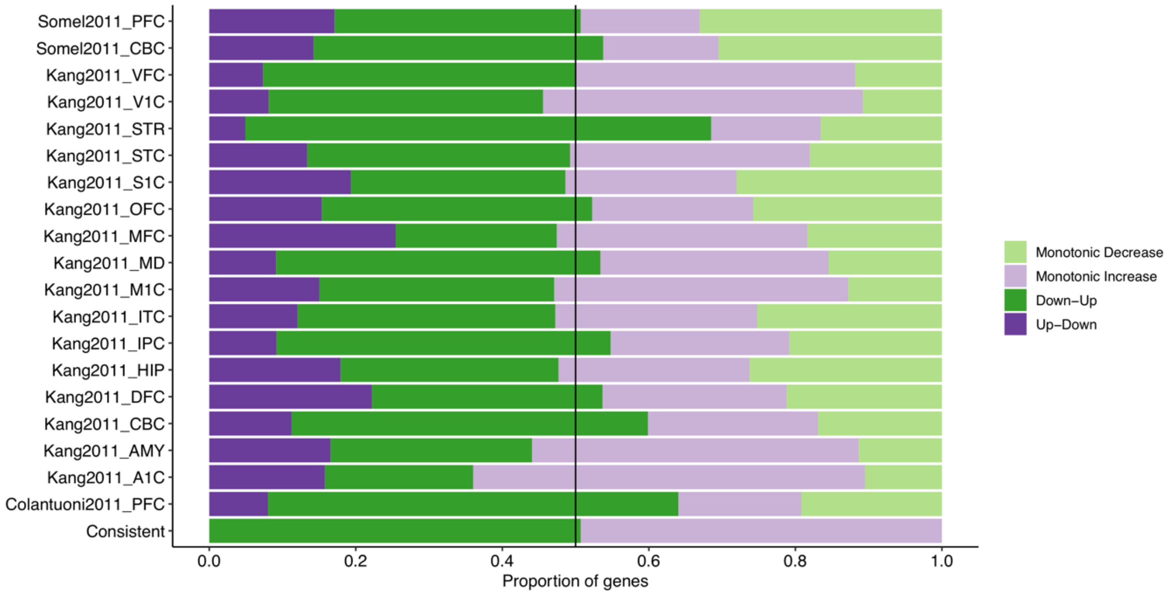


**Figure S9:** The proportion of different trends in age-related heterogeneity change in each dataset and among the genes showing a consistent increase across aging datasets ($n$ = 147). No effect size or significance cutoff was used. Up-down: increase in development & decrease in aging; down-up: decrease in development & increase in aging; monotonic increase: increase in development and aging; monotonic decrease: decrease in development and aging.

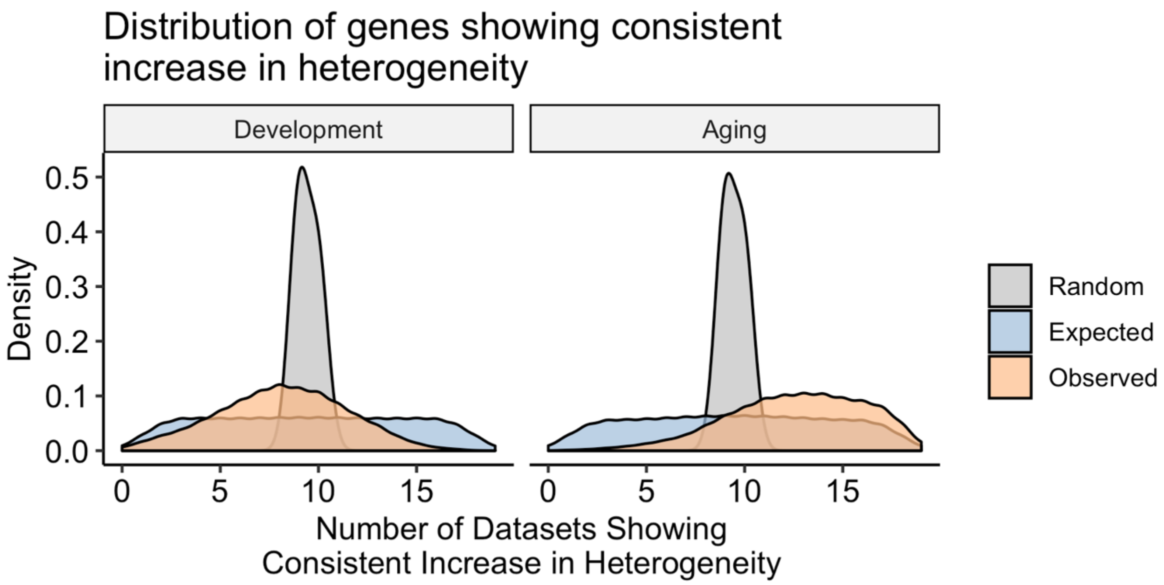


**Figure S10:** Random (expectation calculated with independent permutations), expected (based on permutations taking dataset dependency into account) and observed consistency in the heterogeneity change across datasets in development and aging. This plot corresponds to Figure 3c, but here we also include the expectation based on independent permutations ('Random'), and demonstrate that it is less stringent.

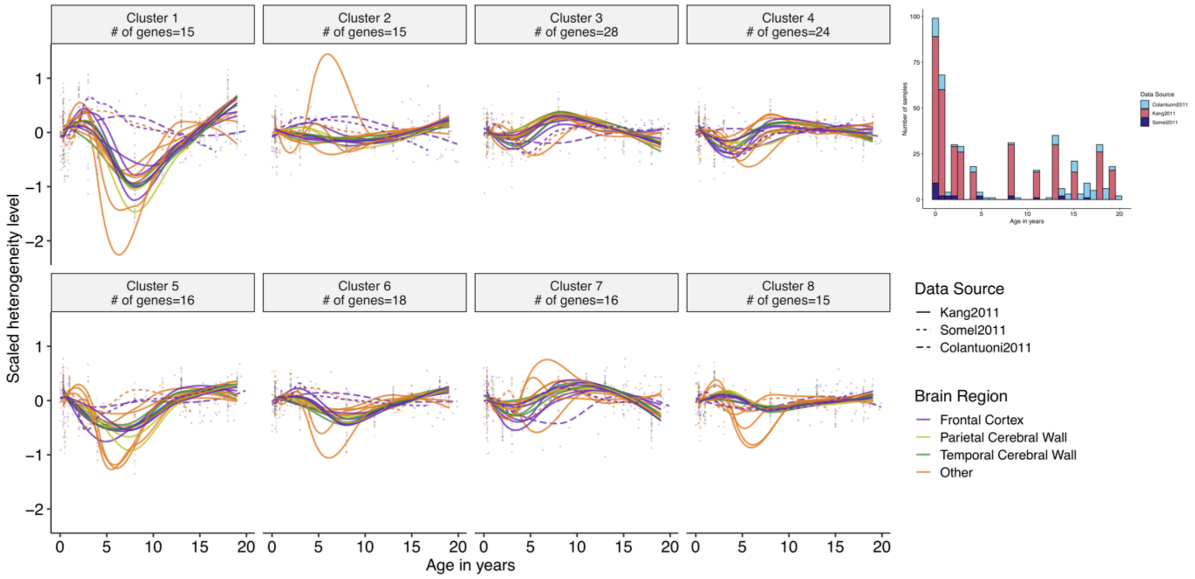


**Figure S11:** Heterogeneity trajectories in development for the genes that were clustered according to their heterogeneities in aging, using k-means clustering method (in Figure 4). The x-axis shows the age and the y-axis shows scaled heterogeneity level (residuals from linear model). Spline curves represent mean age-related heterogeneity changes of genes in each cluster, from each dataset and brain region. The colors and line-types of curves specify different brain regions and data sources, respectively. Different pattern observed between age of 5 to 8 could be biologically relevant but it is important to note that the number of samples in this age range is low. The age distribution of samples in development datasets is given as a bar plot in upper right corner.

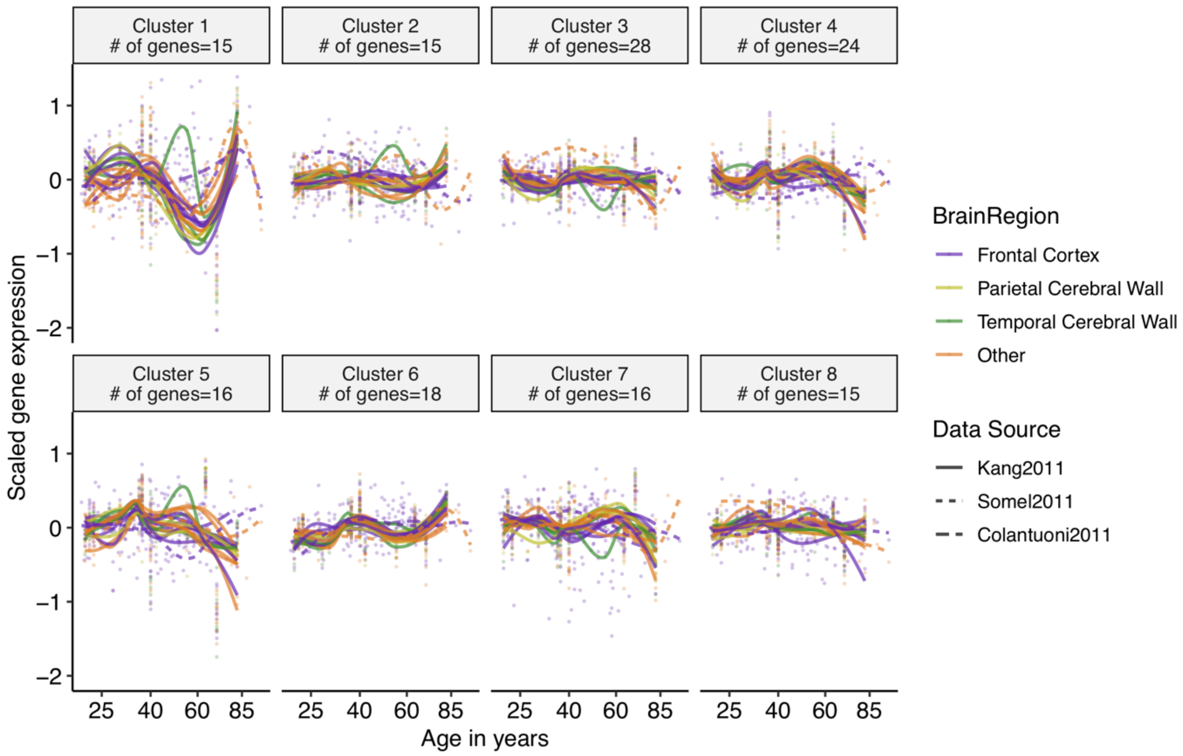


**Figure S12:** Gene expression trajectories of the genes that were clustered according to their heterogeneities, using k-mean clustering method (in Figure 4). The x-axis shows the age on the fourth root scale, and the y-axis shows scaled gene expression values. Spline curves represent mean age-related expression changes of genes in each cluster, from each dataset and brain region. The colors and line-types of curves specify different brain regions and data sources, respectively.

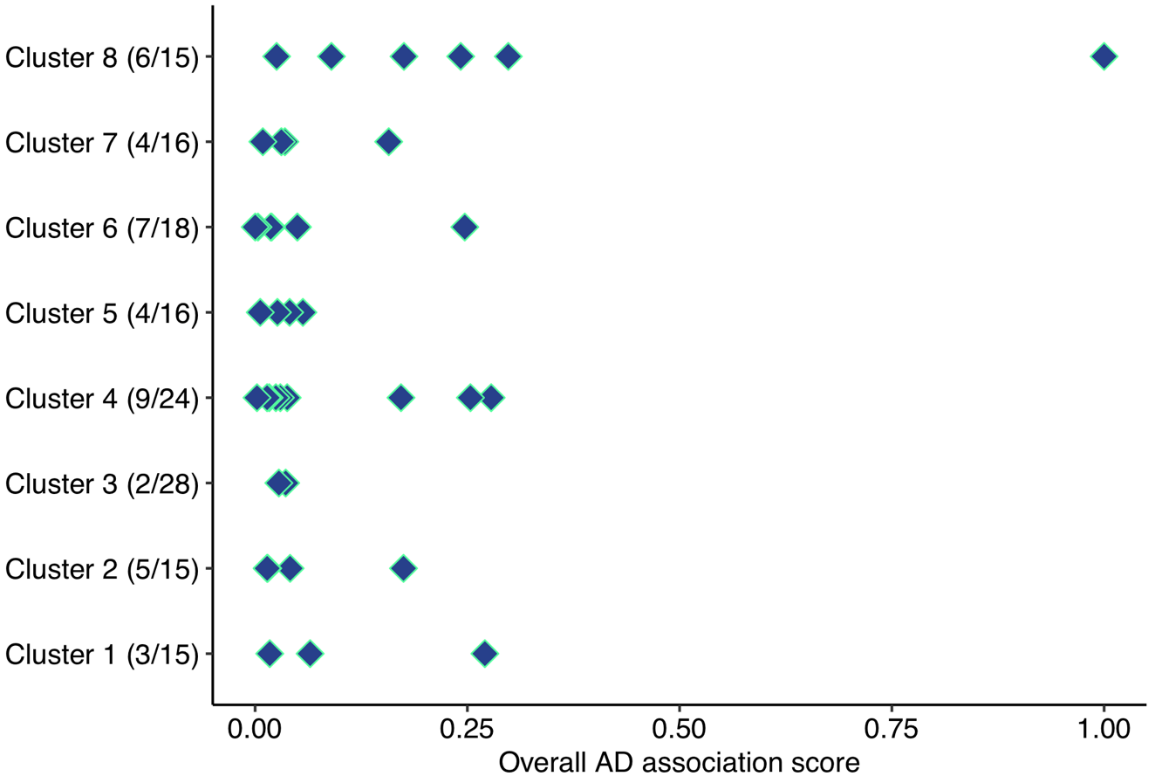


**Figure S13:** Association between the Alzheimer's-related genes and the genes that consistently become more heterogeneous in aging (*n* = 147) and belong to different heterogeneity trajectories (Figure 4). The x-axis shows overall score for the association with Alzheimer's Disease, while the y-axis shows different clusters in Figure 4. Numbers in the parenthesis on the y-axis reflect the proportion of the genes in clusters found to be in association with AD (40/147, overall).

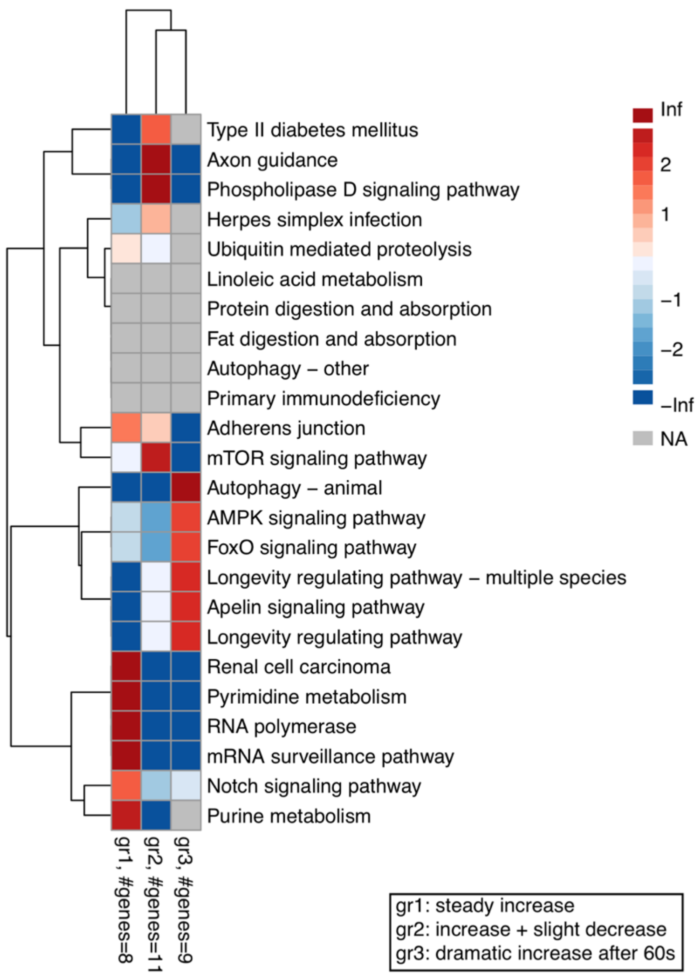


**Figure S14:** Heatmap showing the association between different heterogeneity trajectories and KEGG pathways that are significantly associated with a consistent change in heterogeneity during aging. Heterogeneity trajectories are based on the definitions in the main text and i) gr1 includes clusters 3 and 7, ii) gr2 includes clusters 4, 5, and 8, and iii) gr3 includes clusters 1, 2, and 6. The colors represent the Odd's ratio, where red shows enrichment and blue shows depletion.

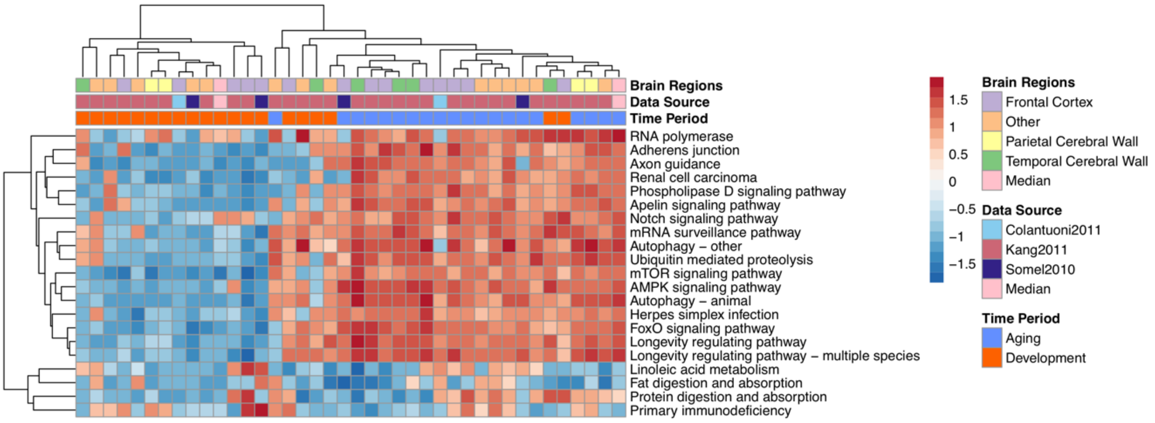


**Figure S15:** A heatmap of the normalized enrichment scores for KEGG pathways that are significantly associated with consistent heterogeneity in aging (rows) and datasets (columns). The color represents normalized enrichment scores; red: positive, blue: negative and the darkness specifies how strong the association is.

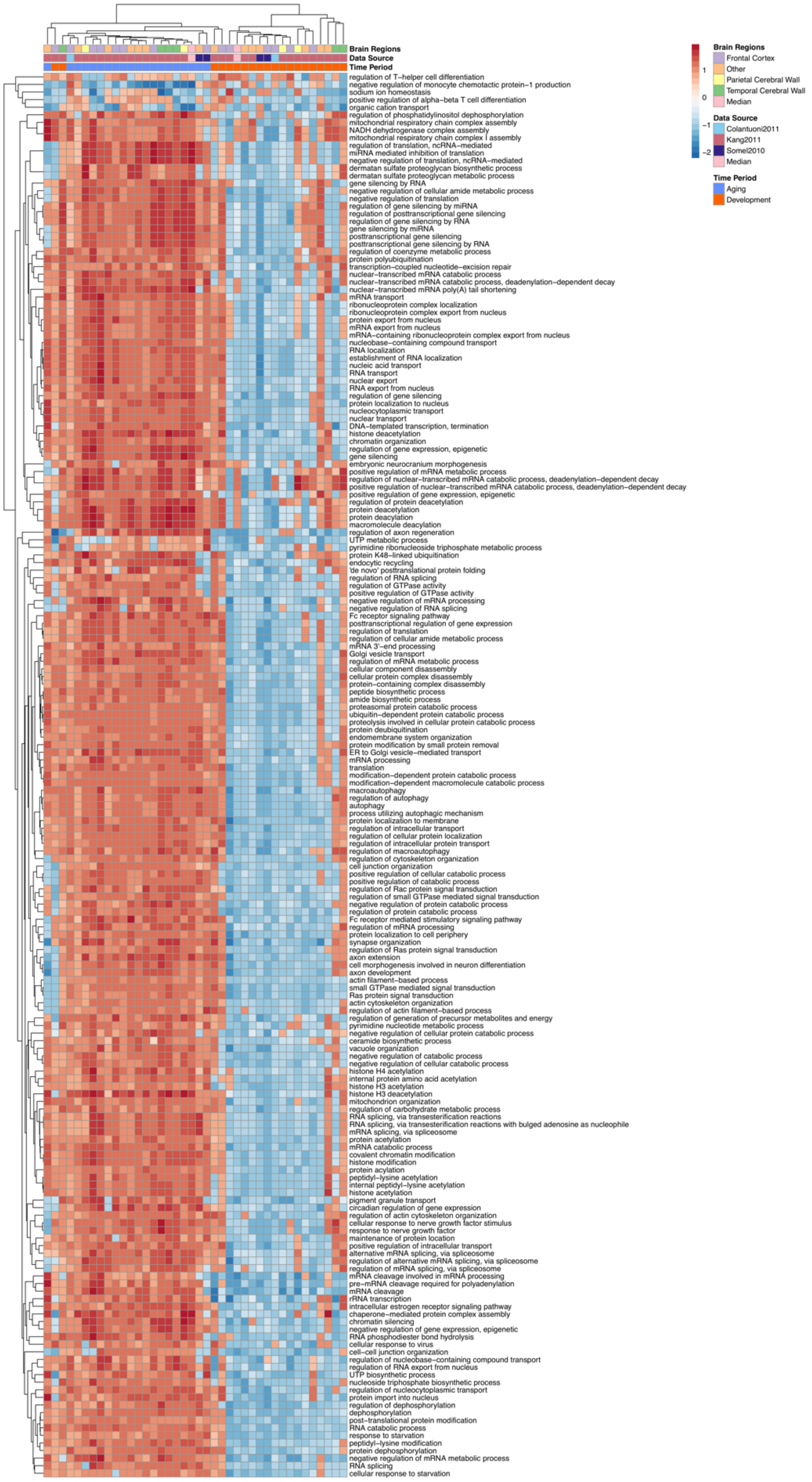


**Figure S16:** The same as Figure S15 but for Gene Ontology Biological Process categories.

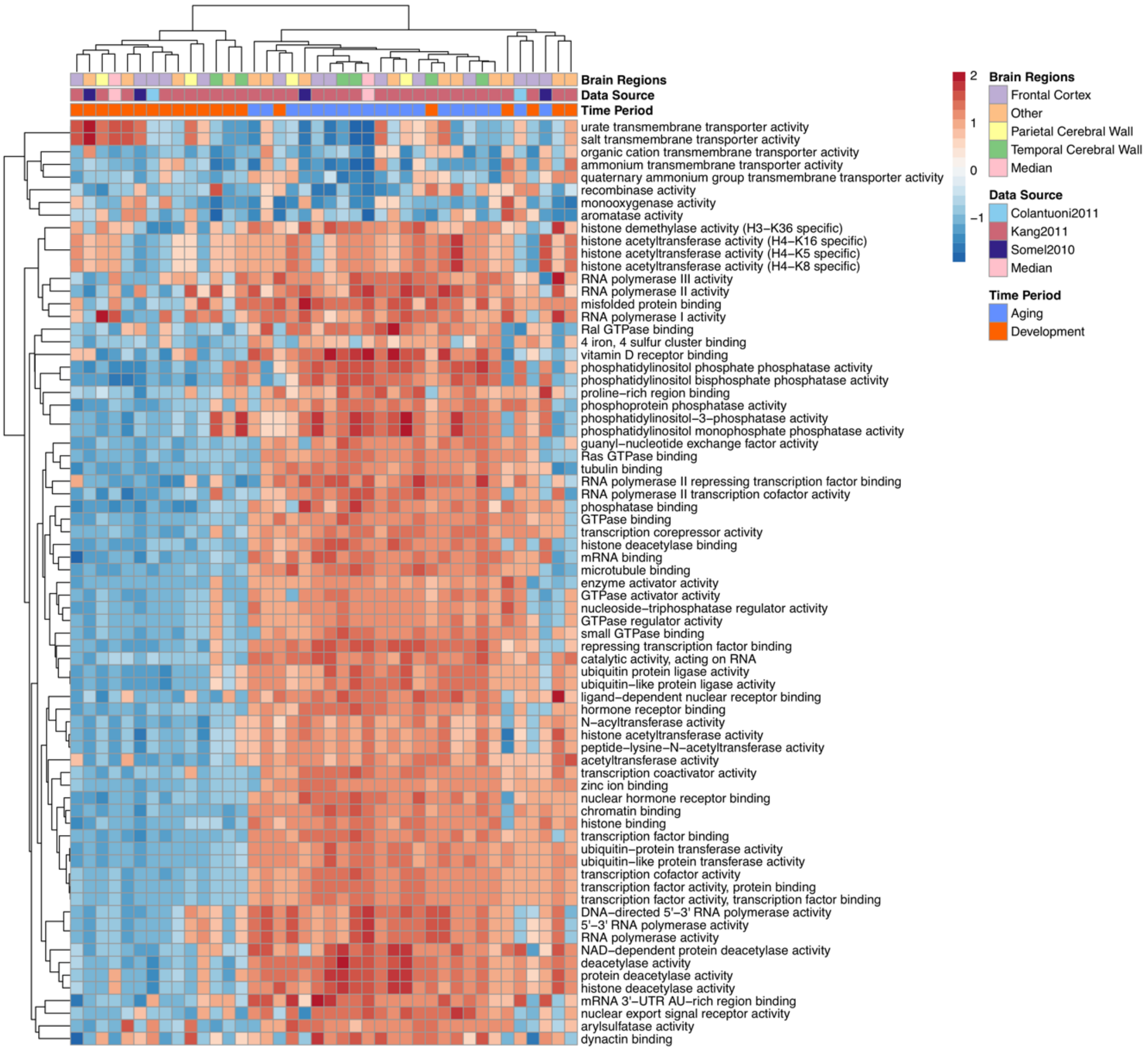


**Figure S17:** The same as Figure S15 but for Gene Ontology Molecular Function categories.

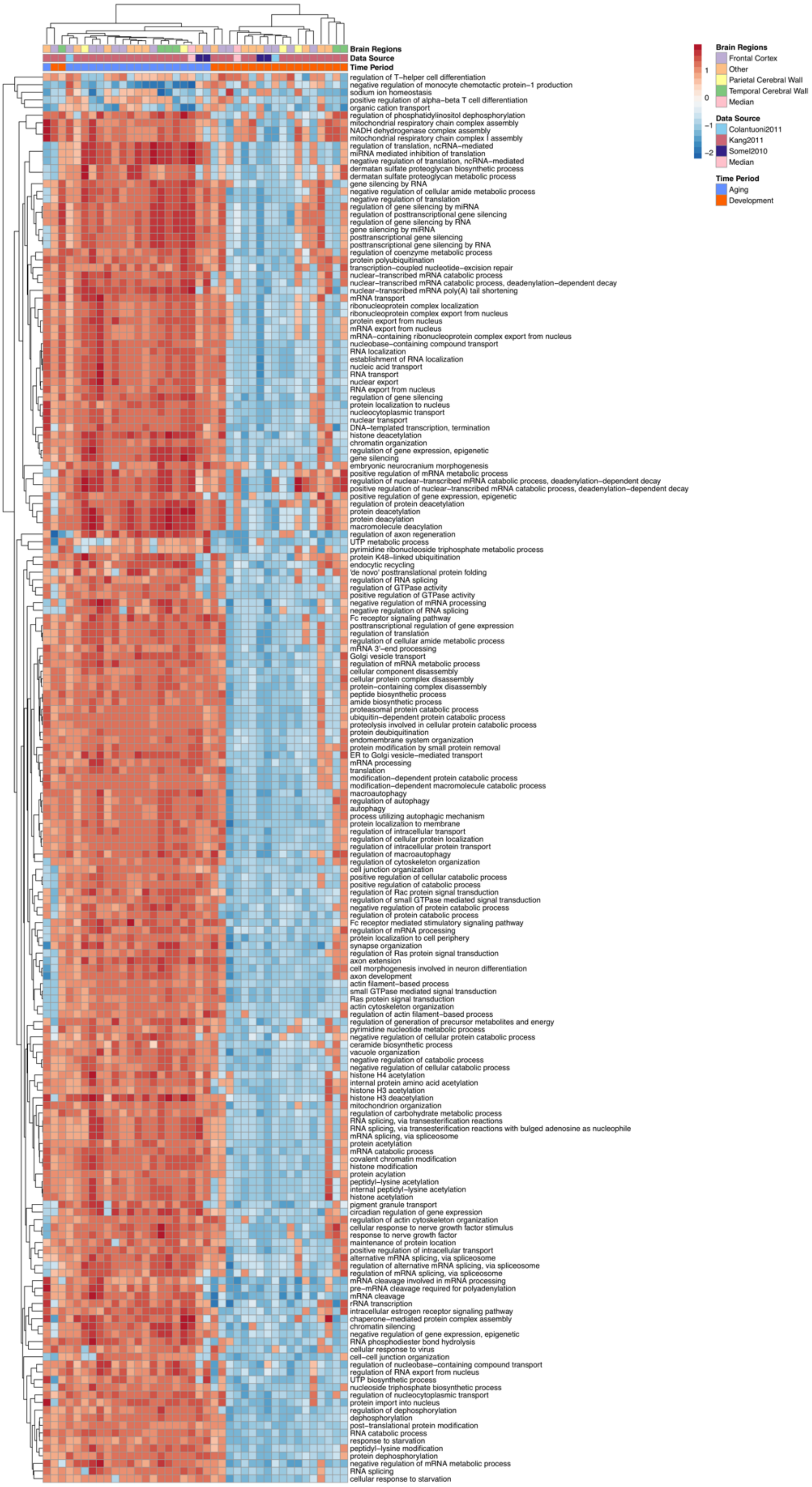


**Figure S18:** The same as Figure S15 but for Gene Ontology Cellular Component categories.

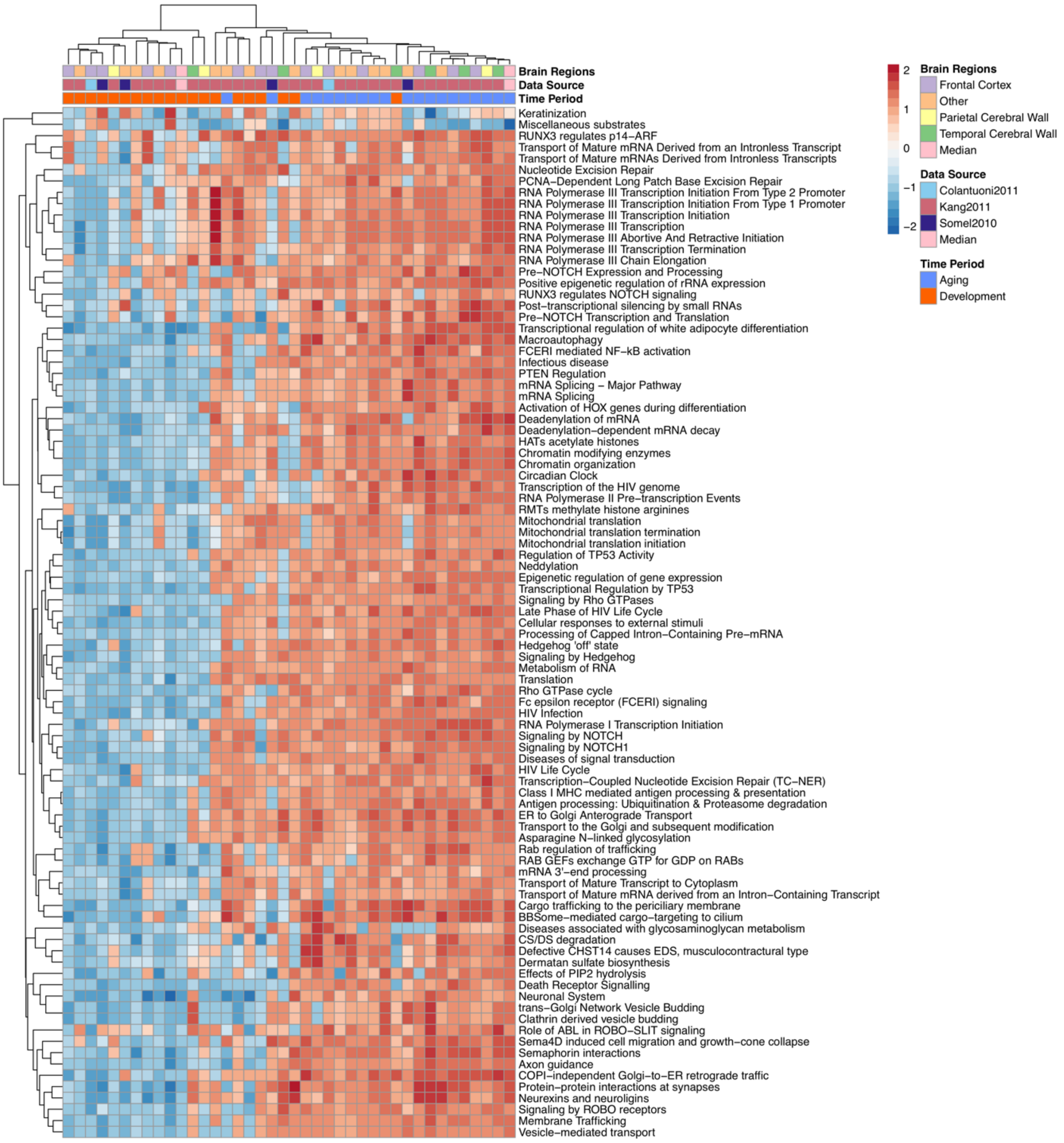


**Figure S19:** The same as Figure S15 but for Reactome pathways.

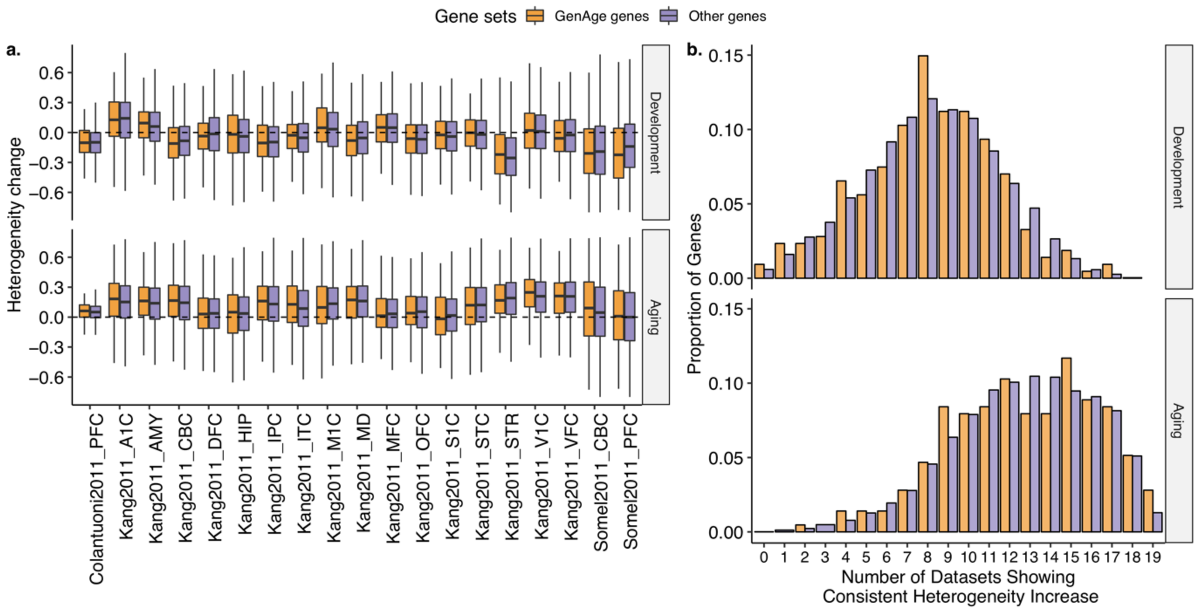


**Figure S20:** Association between GenAge human gene set (*n* = 214) and age-related heterogeneity. (a) Boxplots show age-related heterogeneity changes (ρ values) (y-axis) of genes from GenAge gene set (orange) and the remaining genes (purple) in different datasets (x-axis) during aging (lower panel) and development (upper panel). (b) Consistency in age-related heterogeneity increase in genes from GenAge gene set (orange) and the other genes (purple) in aging (lower panel) and development (upper panel). The x-axis shows the number of datasets among which genes show consistent heterogeneity increase, while the y-axis shows the proportions of the number of genes.

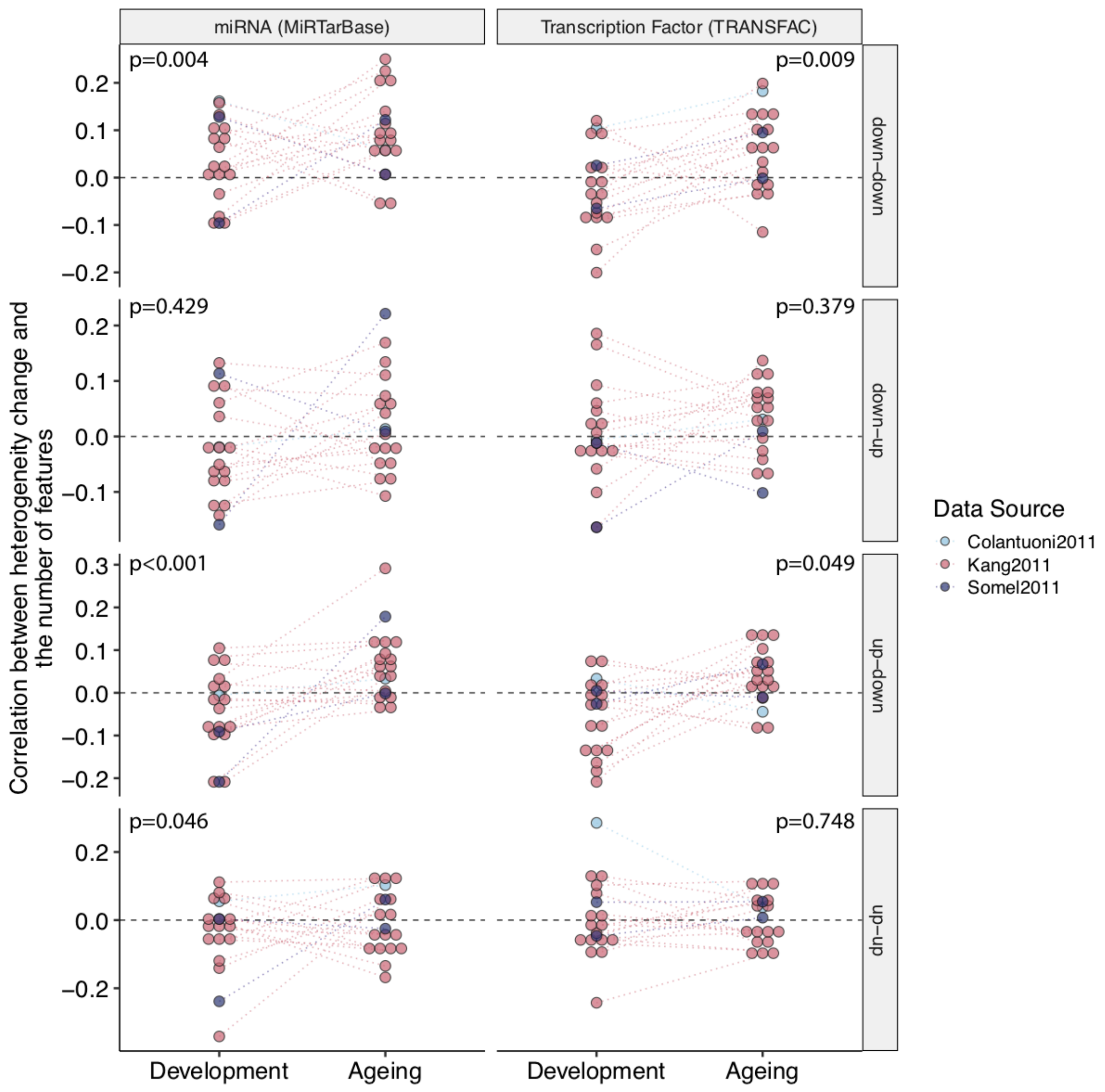


**Figure S21:** Correlation between the change in heterogeneity and number of transcriptional regulators, i.e. miRNA and transcription factors. Each point represents a dataset, and the color shows the data source. *p*-values are calculated using a permutation test. The dashed line at y = 0 shows zero correlation. Genes were divided into four sets based on the change in their expression level in development and aging, e.g. “down-down” includes genes with decreased expression in both development and aging, whereas “down-up” includes genes with decreasing expression level in development and then increase in aging.

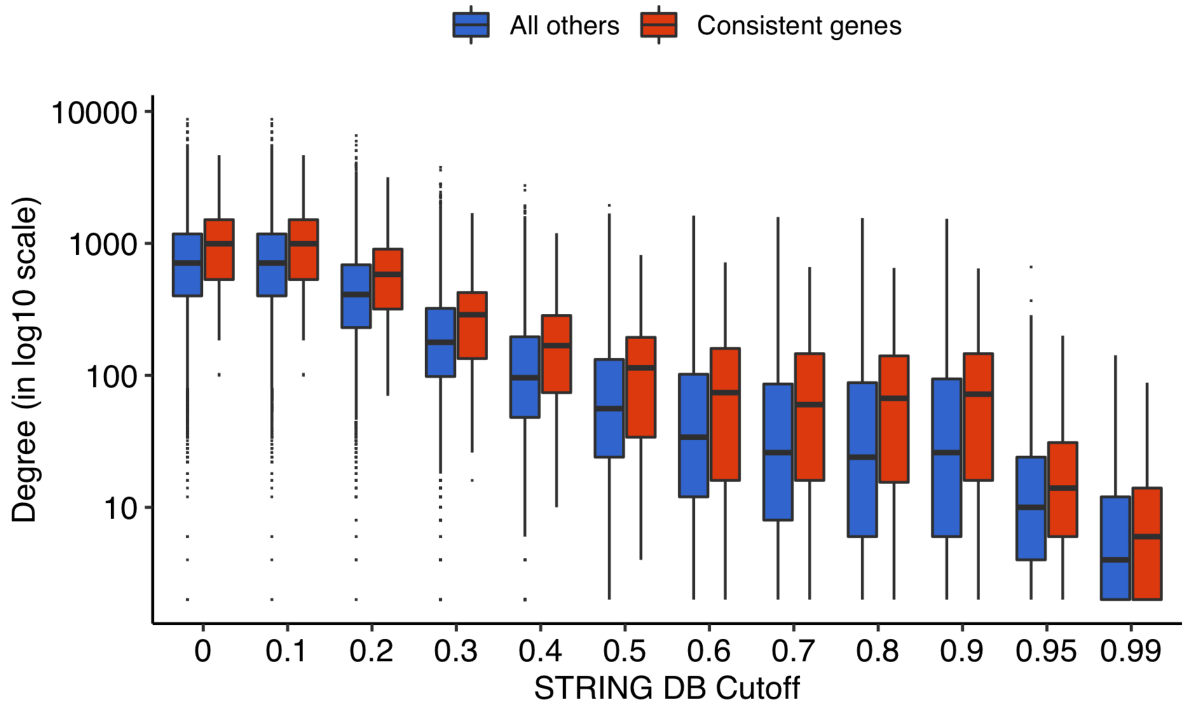


| Cutoff | Number of Genes | Number of Interactions | p | FDR |
|---|---|---|---|---|
| 0.00 | 11,016 | 4,920,316 | <0.0001 | <0.0001 |
| 0.10 | 11,016 | 4,920,316 | <0.0001 | <0.0001 |
| 0.20 | 11,015 | 2,918,350 | 0.0001 | 0.0001 |
| 0.30 | 11,015 | 1,376,452 | <0.0001 | <0.0001 |
| 0.40 | 11,008 | 845,298 | <0.0001 | <0.0001 |
| 0.50 | 10,955 | 578,970 | <0.0001 | <0.0001 |
| 0.60 | 10,665 | 438,686 | <0.0001 | <0.0001 |
| 0.70 | 9,881 | 353,362 | <0.0001 | <0.0001 |
| 0.80 | 8,588 | 305,128 | <0.0001 | <0.0001 |
| 0.90 | 7,386 | 270,942 | 0.0001 | 0.0001 |
| 0.95 | 5,800 | 58,190 | 0.0235 | 0.0256 |
| 0.99 | 3,167 | 17,660 | 0.3989 | 0.3989 |

**Figure S22:** Degree distributions in protein-protein interaction database (STRING) for consistent genes (red) and all others (blue). The y-axis show degree (number of interactors) in log scale. The x-axis shows different cutoffs for interaction confidence used to filter STRING database. The color represents two sets of genes; red: consistent genes that show consistent increase in heterogeneity across all aging datasets, blue: all other genes. Interaction degree is significantly higher in consistent genes across all cutoffs except for 0.99 (permutation test). Details, including the number of genes, interactions and *p*-values, are given as a table.

| Cell Type | Odds Ratio | FDR | Total Cell–type | Total Heterogeneus | a | b | c | d | p |
|---|---|---|---|---|---|---|---|---|---|
| OLs | 5.537 | 0.179 | 51 | 9 | 2 | 49 | 7 | 953 | 0.072 |
| Myelin_OLs | 3.854 | 0.179 | 348 | 9 | 6 | 342 | 3 | 660 | 0.071 |
| OPCs | 0.265 | 0.357 | 322 | 9 | 1 | 321 | 8 | 681 | 0.286 |
| Neurons | 0.000 | 0.205 | 272 | 9 | 0 | 272 | 9 | 730 | 0.123 |
| Astrocytes | 0.000 | 1.000 | 18 | 9 | 0 | 18 | 9 | 984 | 1.000 |

**Figure S23:** Cell-type specificity analysis for genes that become heterogenous with age across all aging datasets. Fisher's exact test is used to calculate the association (Odds Ratio) and *p*-value. FDR corrected *p*-value (FDR column) is also given.

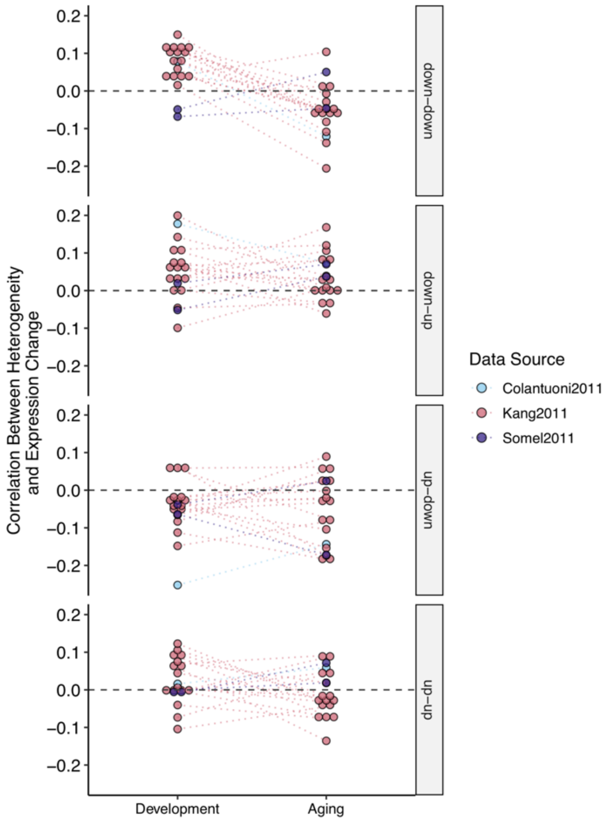


**Figure S24:** The relationship between the age-related changes in gene expression (β values) and changes in heterogeneity (ρ values). The y-axis shows the Spearman's correlation coefficients, while different periods are in the x-axis. Facets show different trends in expression changes: up-down: up-regulation in development and down-regulation in aging; down-up: down-regulation in development and up-regulation in aging; up-up: up-regulation in development and up-regulation in aging; down-down: down-regulation in development and down-regulation in aging.

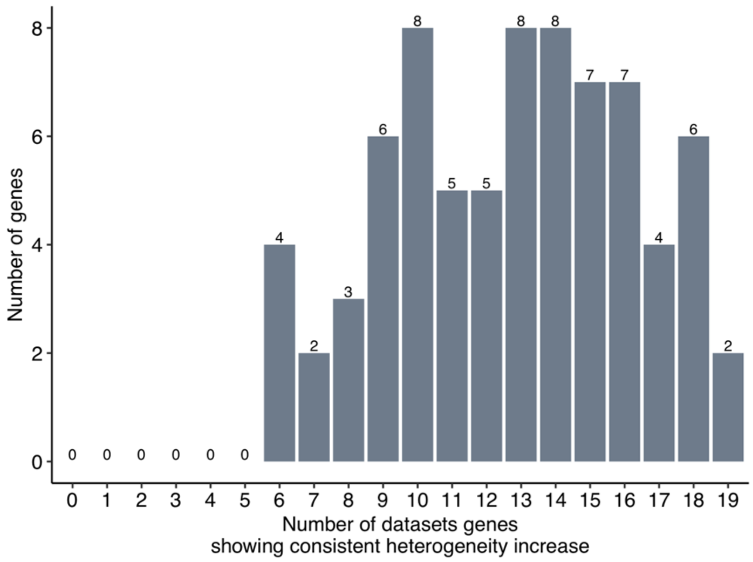


**Figure S25:** Analysis of the consistency in aging-related heterogeneity of the post-mortem interval (PMI)-associated genes. The x-axis shows the number of datasets in which genes were showing a consistent increase in age-related heterogeneity during aging, while y-axis reflects the number of genes. There were 105 previously identified PMI associated genes in the human cerebral cortex (see main text), 75 of which were included in our analyses. We asked if these PMI associated genes show more increase in heterogeneity, and found that only 2 of 147 consistent genes (i.e. genes showing an increase in heterogeneity in 19 out of 19 aging datasets) were associated with PMI.

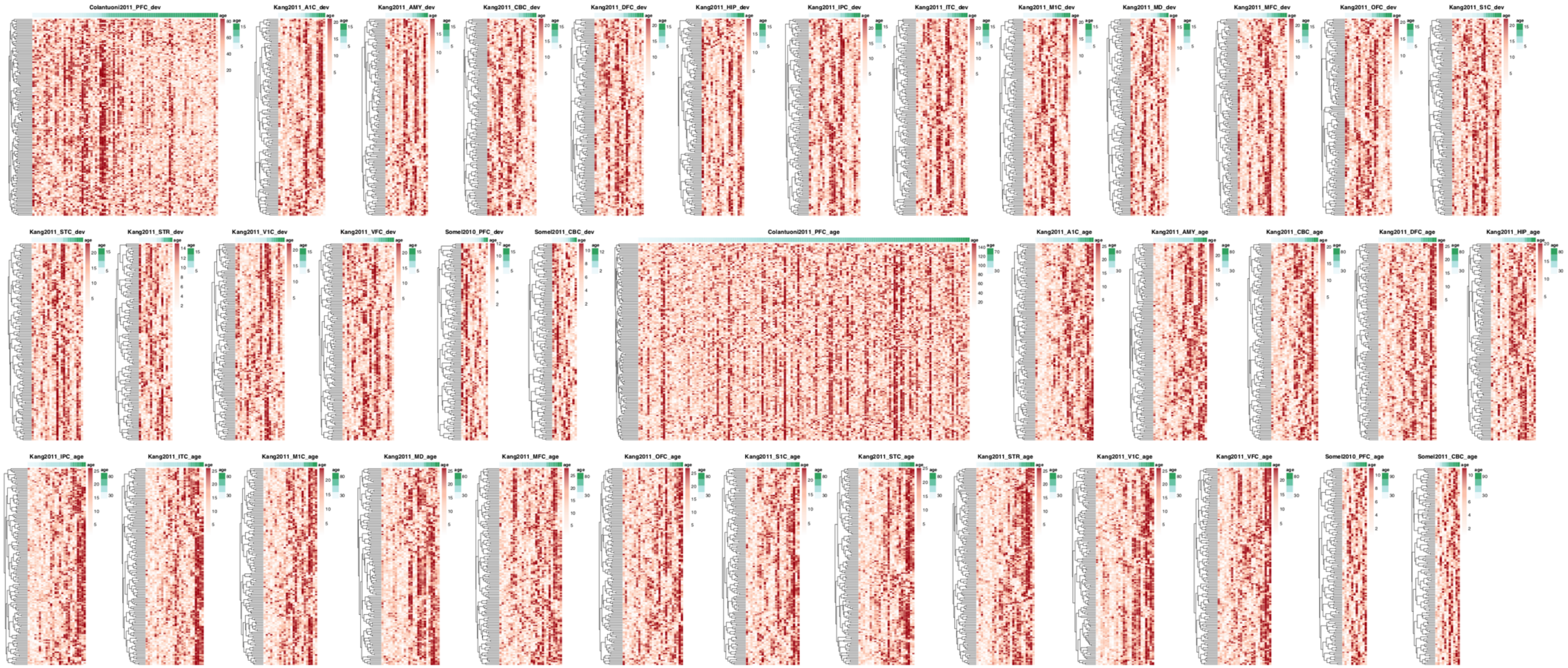


**Figure S26:** Heatmaps for 38 datasets showing the heterogeneity levels (residuals) for each individual (columns) in 147 genes that show consistent increase across all aging datasets (rows). The color shows the rank of individuals with respect to their heterogeneity for that particular gene. Darker colors show individuals with the highest heterogeneity for that particular gene. Genes are clustered using hierarchical clustering and individuals are ordered by age (darker green in column annotation showing older ages).

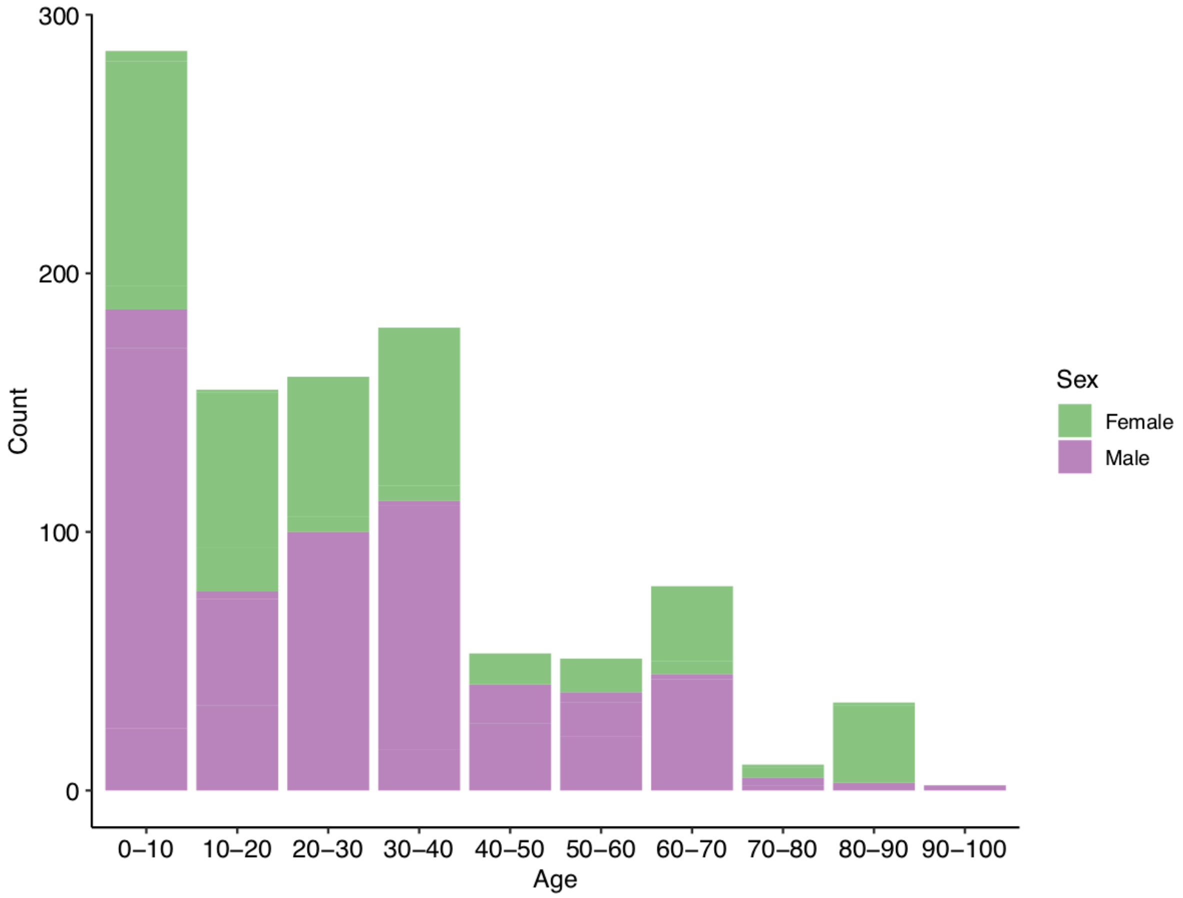


**Figure S27:** The proportions of sexes of samples used in the analysis. The x- and y-axes show age intervals and number of samples, respectively. The colors specify the proportion of male (purple) and female (green) samples.

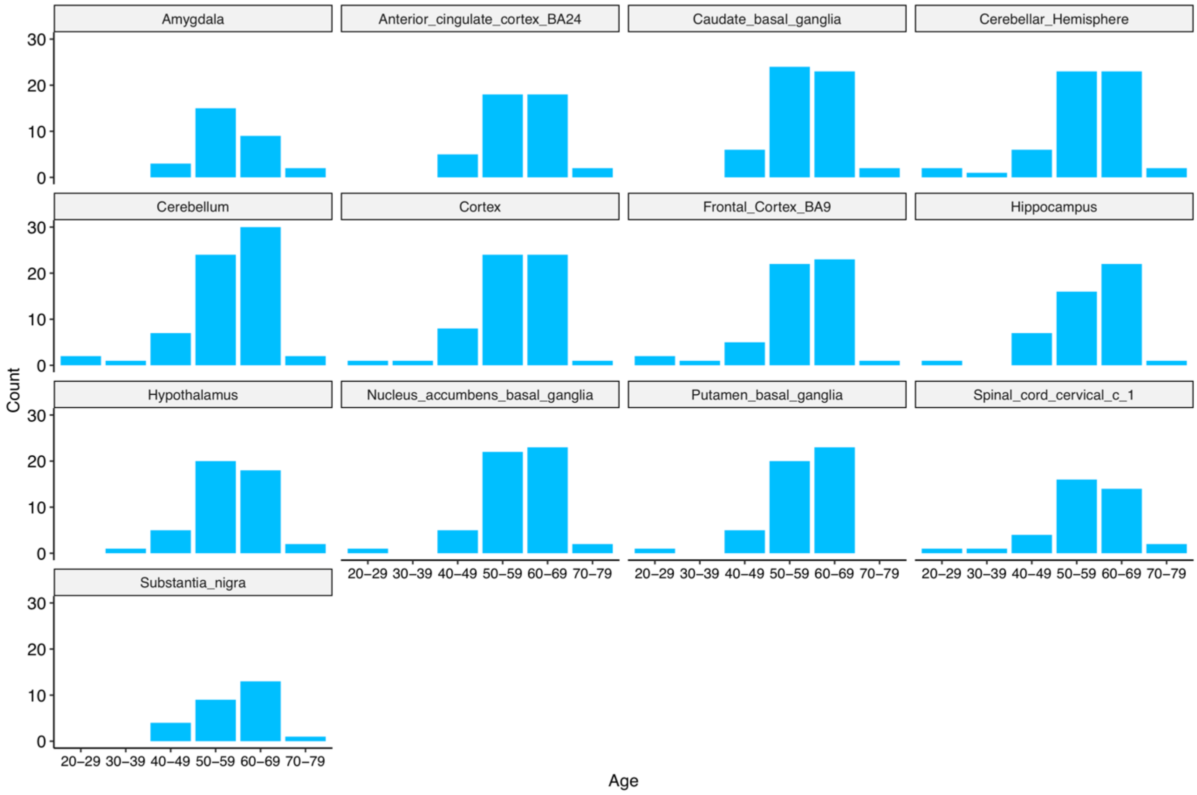


**Figure S28:** Age distribution of the individuals used in GTEx RNA-Seq datasets.

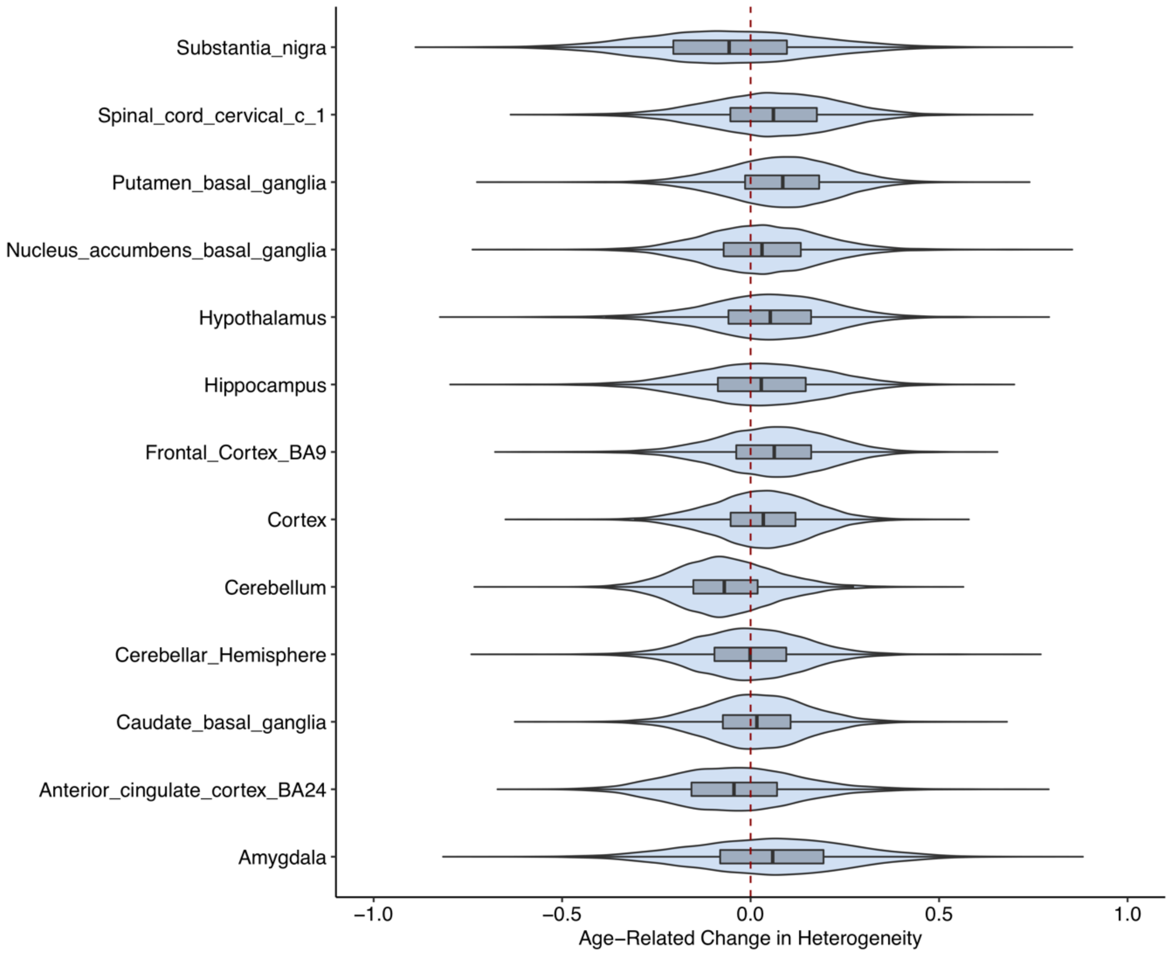


**Figure S29:** Distribution of age-related changes in heterogeneity (rho values) (x-axis) in different GTEx datasets, corresponding to different brain regions in (y-axis). The red dotted vertical line (at x = 0) reflects no change in heterogeneity.

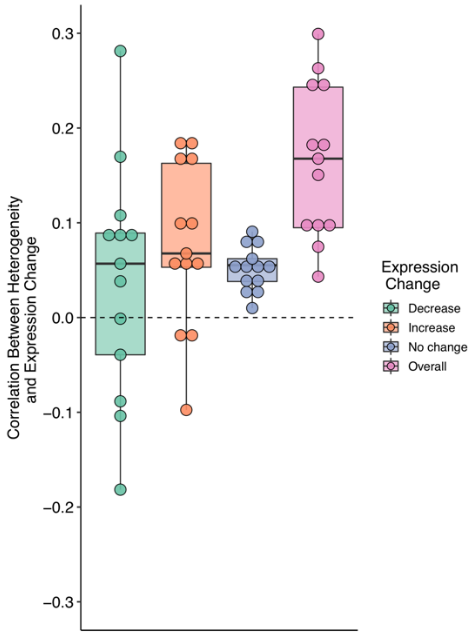


**Figure S30:** The relationship between expression and heterogeneity changes in GTEx datasets. The y-axis shows Spearmen correlation coefficients calculated between age-related expression changes ($\beta$ values) and heterogeneity changes ($\rho$ values) separately for each dataset. Different colors indicate the direction of expression change. Genes with the beta values within the range of -0.1 and 0.1 were assumed to be no change. The overall includes all the genes irrespective of the direction of their expression change.
