## Supplementary Material - Pathway results for "Gene expression heterogeneity during brain development and aging: temporal changes and functional consequences"

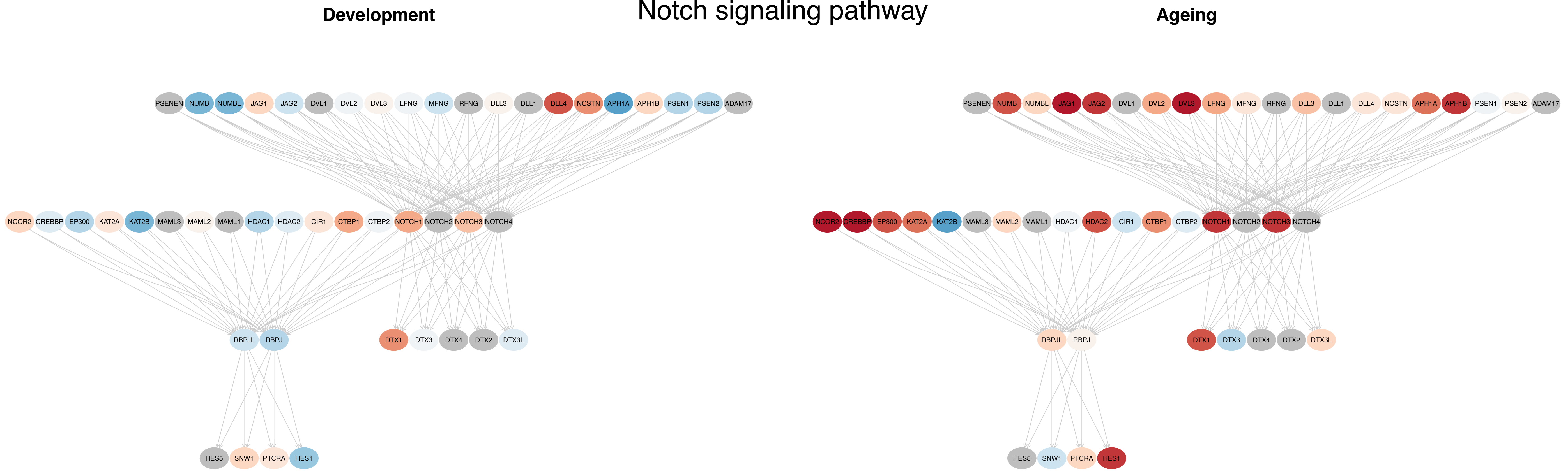

Notch signaling pathway
Development
Ageing
PSENEN NUMB NUMBL JAG1 JAG2 DVL1 DVL2 DVL3 LFNG MFNG RFNG DLL3 DLL1 DLL4 NCSTN APH1A APH1B PSEN1 PSEN2 ADAM17
NCOR2 CREBBP EP300 KAT2A KAT2B MAML3 MAML2 MAML1 HDAC1 HDAC2 CIR1 CTBP1 CTBP2 NOTCH1 NOTCH2 NOTCH3 NOTCH4
RBPJL RBPJ DTX1 DTX3 DTX4 DTX2 DTX3L
HES5 SNW1 PTCRA HES1
PSENEN NUMB NUMBL JAG1 JAG2 DVL1 DVL2 DVL3 LFNG MFNG RFNG DLL3 DLL1 DLL4 NCSTN APH1A APH1B PSEN1 PSEN2 ADAM17
NCOR2 CREBBP EP300 KAT2A KAT2B MAML3 MAML2 MAML1 HDAC1 HDAC2 CIR1 CTBP1 CTBP2 NOTCH1 NOTCH2 NOTCH3 NOTCH4
RBPJL RBPJ DTX1 DTX3 DTX4 DTX2 DTX3L
HES5 SNW1 PTCRA HES1

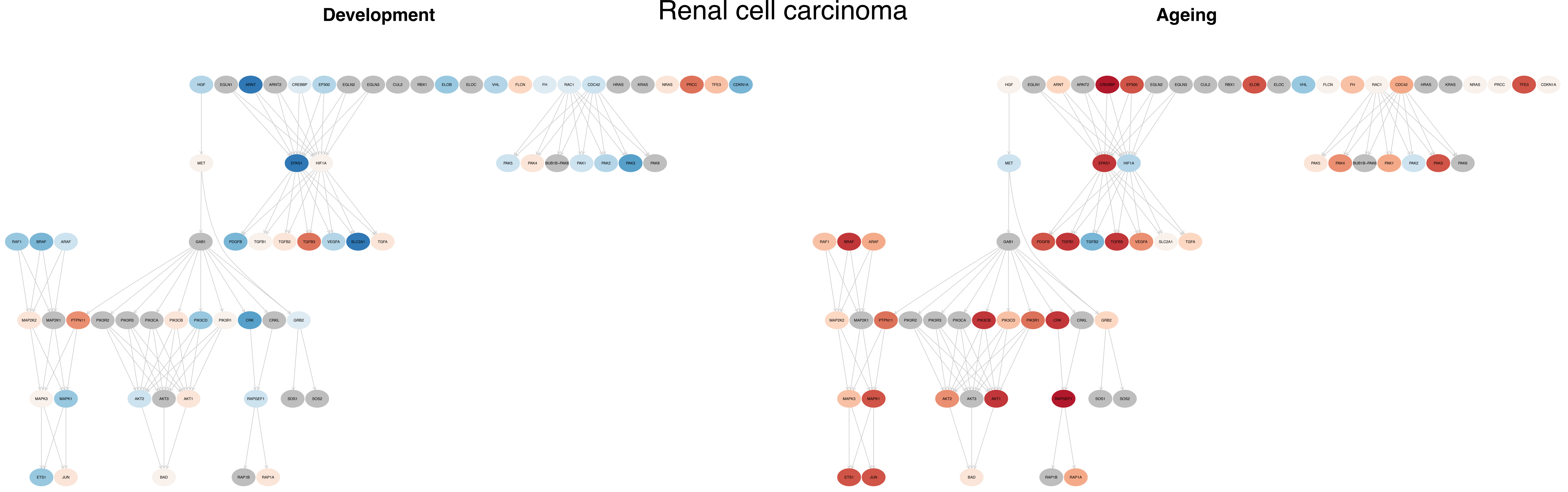

Renal cell carcinoma
Development
Ageing
HGF
EGLN1
ARNT
ARNT2
CREBBP
EP300
EGLN2
EGLN3
CUL2
RBX1
ELOB
ELOC
VHL
FLCN
FH
RAC1
CDC42
HRAS
KRAS
NRAS
PRCC
TFE3
CDKN1A
MET
EPAS1
HIF1A
PAK5
PAK4
BUB1B-PAK6
PAK1
PAK2
PAK3
PAK6
RAF1
BRAF
ARAF
GAB1
PDGFB
TGFB1
TGFB2
TGFB3
VEGFA
SLC2A1
TGFA
MAP2K2
MAP2K1
PTPN11
PIK3R2
PIK3R3
PIK3CA
PIK3CB
PIK3CD
PIK3R1
CRK
CRKL
GRB2
MAPK3
MAPK1
AKT2
AKT3
AKT1
RAPGEF1
SOS1
SOS2
ETS1
JUN
BAD
RAP1B
RAP1A

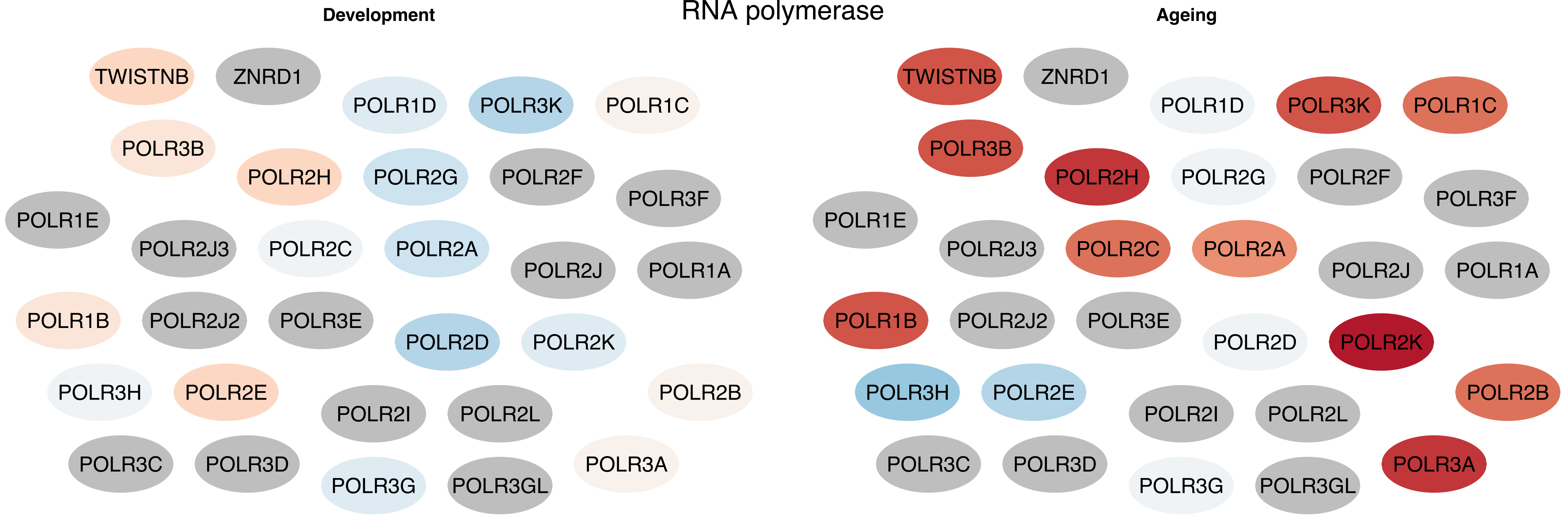
RNA polymerase
Development
TWISTNB
ZNRD1
POLR1D
POLR3K
POLR1C
POLR3B
POLR2H
POLR2G
POLR2F
POLR3F
POLR1E
POLR2J3
POLR2C
POLR2A
POLR2J
POLR1A
POLR1B
POLR2J2
POLR3E
POLR2D
POLR2K
POLR3H
POLR2E
POLR2I
POLR2L
POLR2B
POLR3C
POLR3D
POLR3G
POLR3GL
POLR3A
Ageing
TWISTNB
ZNRD1
POLR1D
POLR3K
POLR1C
POLR3B
POLR2H
POLR2G
POLR2F
POLR3F
POLR1E
POLR2J3
POLR2C
POLR2A
POLR2J
POLR1A
POLR1B
POLR2J2
POLR3E
POLR2D
POLR2K
POLR3H
POLR2E
POLR2I
POLR2L
POLR2B
POLR3C
POLR3D
POLR3G
POLR3GL
POLR3A

### mRNA surveillance pathway

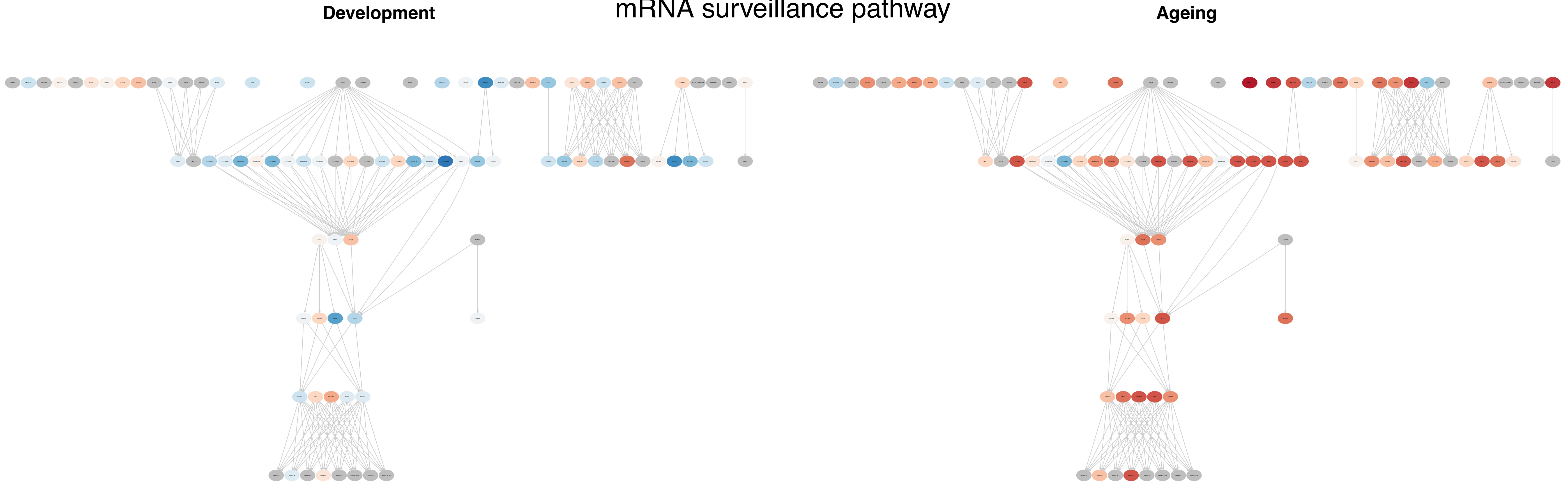

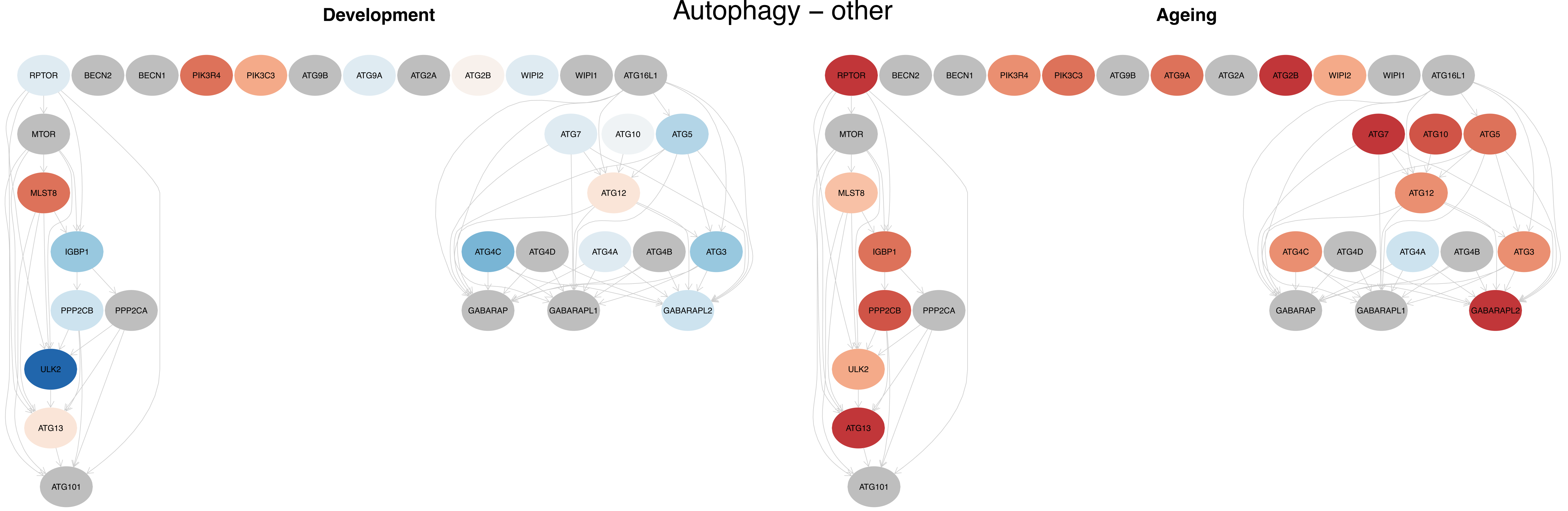
Autophagy – other
Development
Ageing
RPTOR
BECN2
BECN1
PIK3R4
PIK3C3
ATG9B
ATG9A
ATG2A
ATG2B
WIPI2
WIPI1
ATG16L1
MTOR
MLST8
IGBP1
PPP2CB
PPP2CA
ULK2
ATG13
ATG101
ATG7
ATG10
ATG5
ATG12
ATG4C
ATG4D
ATG4A
ATG4B
ATG3
GABARAP
GABARAPL1
GABARAPL2

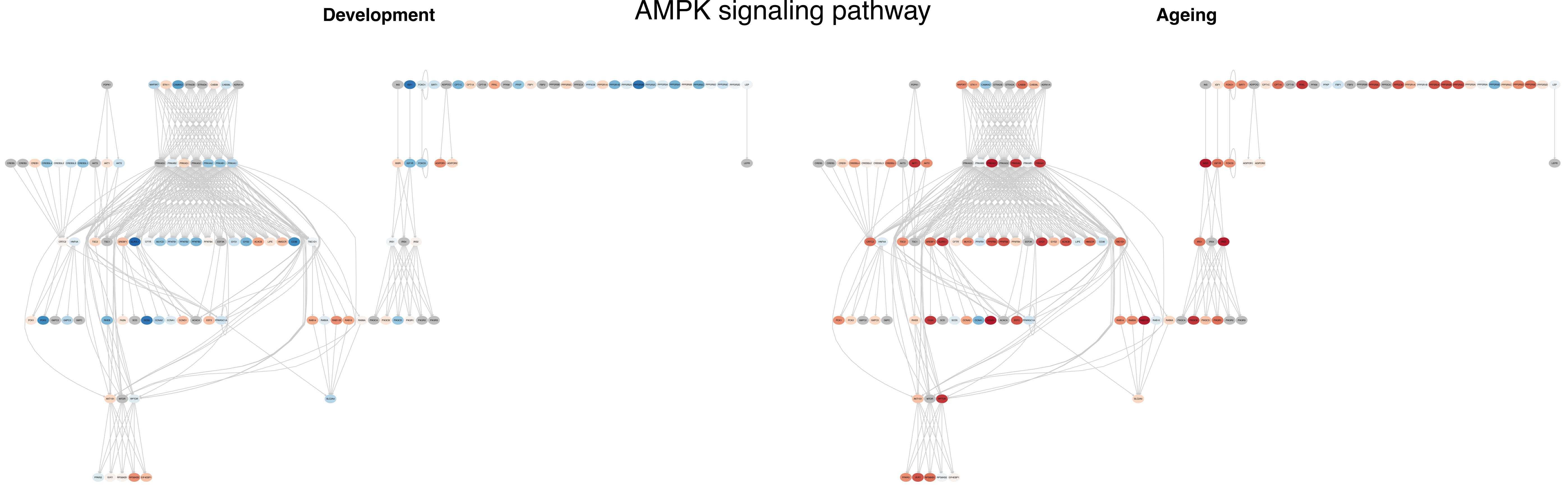
AMPK signaling pathway
Development
Ageing

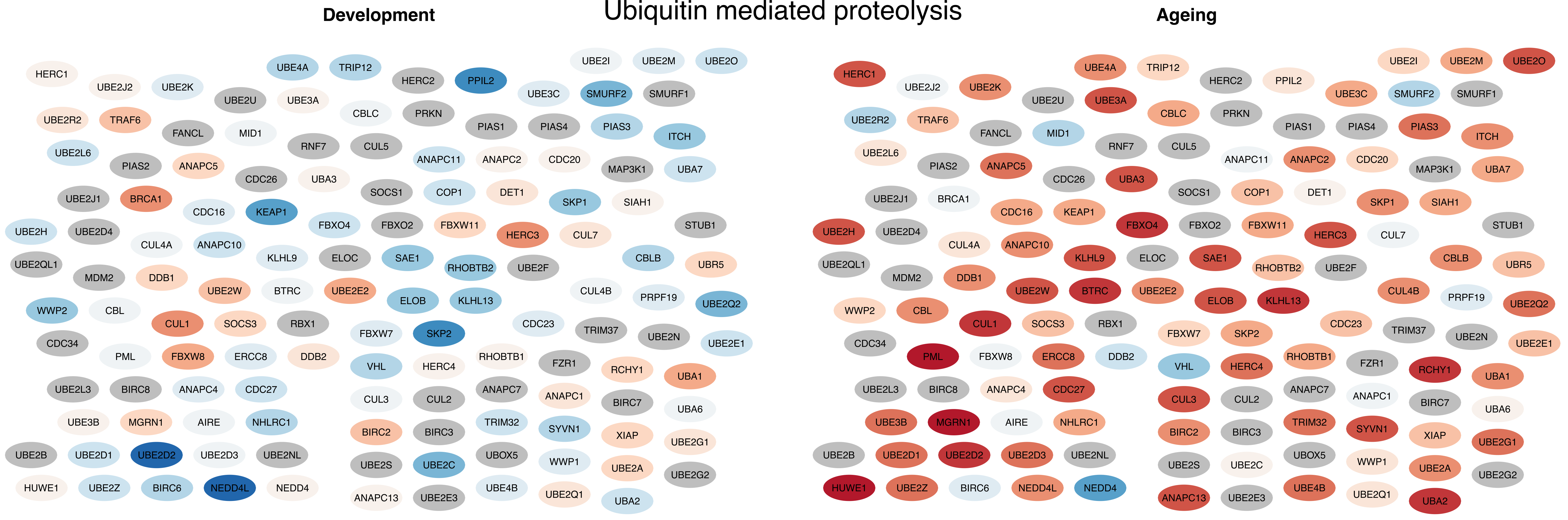
Ubiquitin mediated proteolysis
Development
Ageing
HERC1
UBE2J2
UBE2K
UBE4A
TRIP12
UBE2U
UBE3A
HERC2
PPIL2
UBE2I
UBE2M
UBE2O
UBE3C
SMURF2
SMURF1
UBE2R2
TRAF6
FANCL
MID1
CBLC
PRKN
PIAS1
PIAS4
PIAS3
ITCH
UBE2L6
RNF7
CUL5
ANAPC11
ANAPC2
CDC20
MAP3K1
UBA7
PIAS2
ANAPC5
CDC26
UBA3
UBE2J1
BRCA1
SOCS1
COP1
DET1
SKP1
SIAH1
CDC16
KEAP1
FBXO4
FBXO2
FBXW11
HERC3
CUL7
STUB1
UBE2H
UBE2D4
CUL4A
ANAPC10
KLHL9
ELOC
SAE1
RHOBTB2
UBE2F
CBLB
UBR5
UBE2QL1
MDM2
DDB1
UBE2W
BTRC
UBE2E2
ELOB
KLHL13
CUL4B
PRPF19
UBE2Q2
WWP2
CBL
CUL1
SOCS3
RBX1
FBXW7
SKP2
CDC23
TRIM37
UBE2N
UBE2E1
CDC34
PML
FBXW8
ERCC8
DDB2
VHL
HERC4
RHOBTB1
FZR1
RCHY1
UBA1
UBE2L3
BIRC8
ANAPC4
CDC27
CUL3
CUL2
ANAPC7
ANAPC1
BIRC7
UBA6
UBE3B
MGRN1
AIRE
NHLRC1
TRIM32
SYVN1
XIAP
BIRC2
BIRC3
UBE2G1
UBE2B
UBE2D1
UBE2D2
UBE2D3
UBE2NL
UBOX5
WWP1
UBE2S
UBE2C
UBE2A
UBE2G2
HUWE1
UBE2Z
BIRC6
NEDD4L
NEDD4
UBE4B
UBE2Q1
ANAPC13
UBE2E3
UBA2

#### Phospholipase D signaling pathway

**Development** **Ageing**

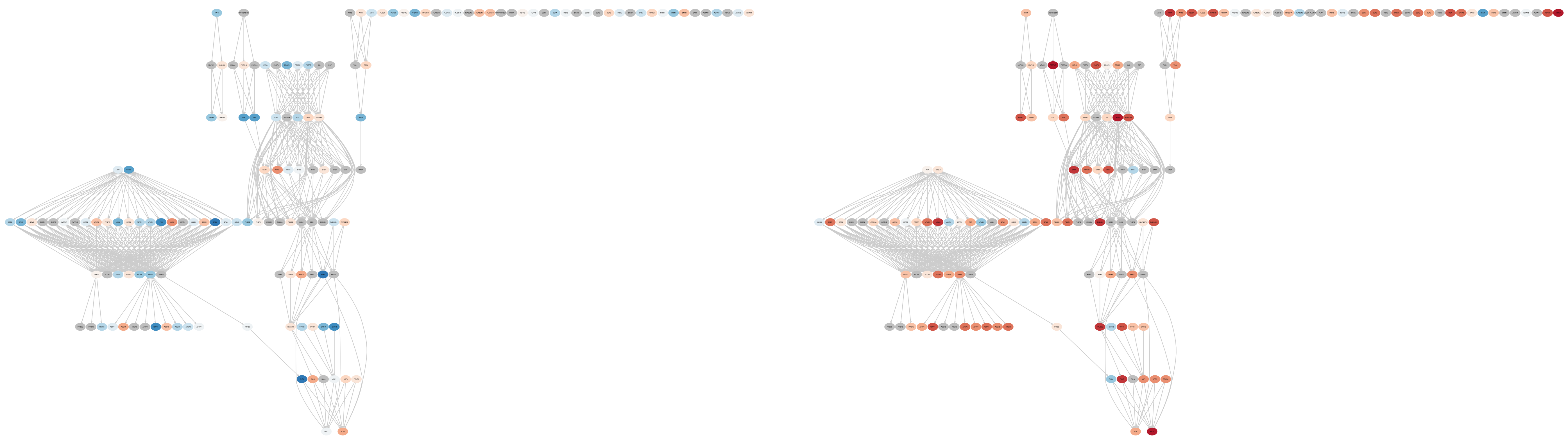

### Longevity regulating pathway – multiple species

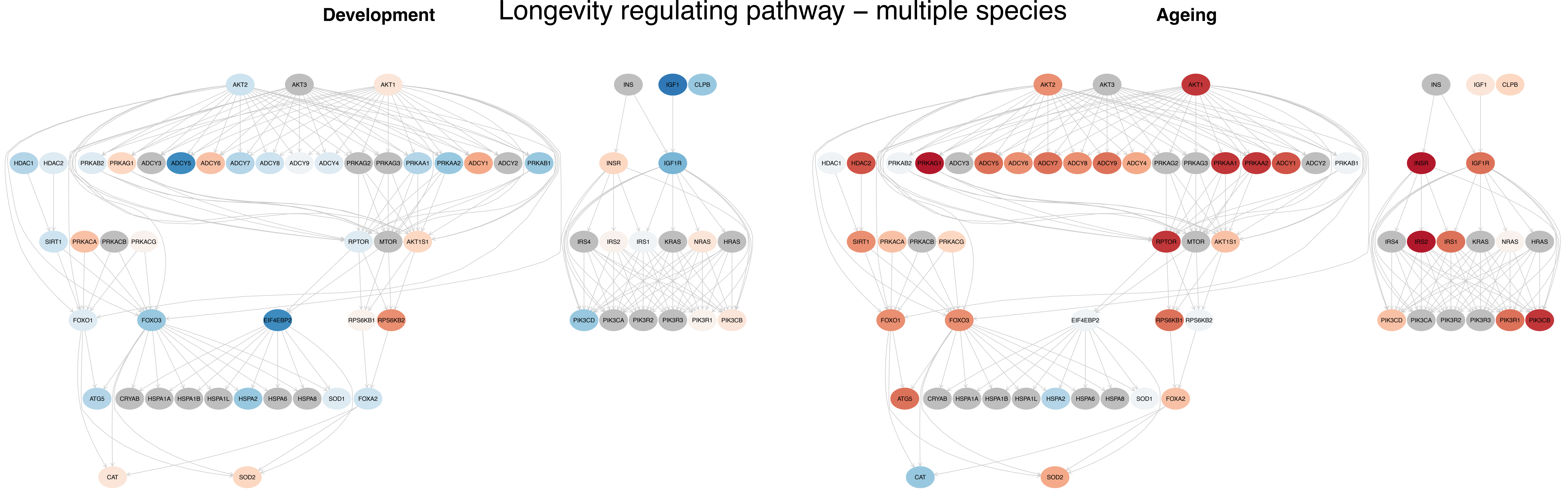

### Longevity regulating pathway

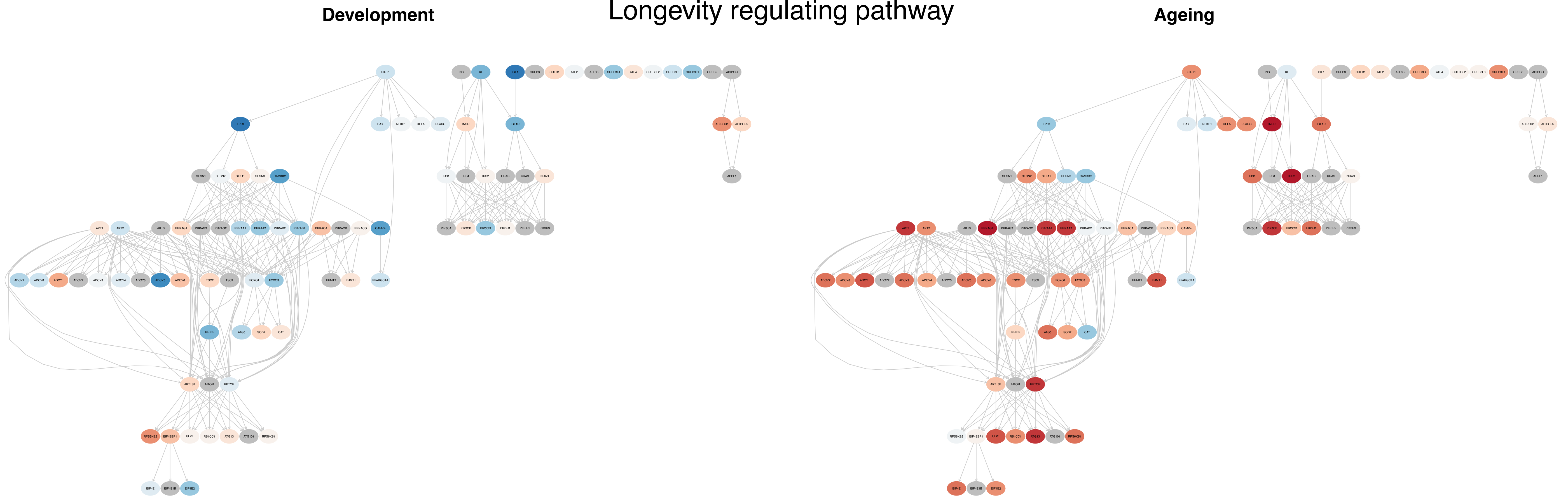

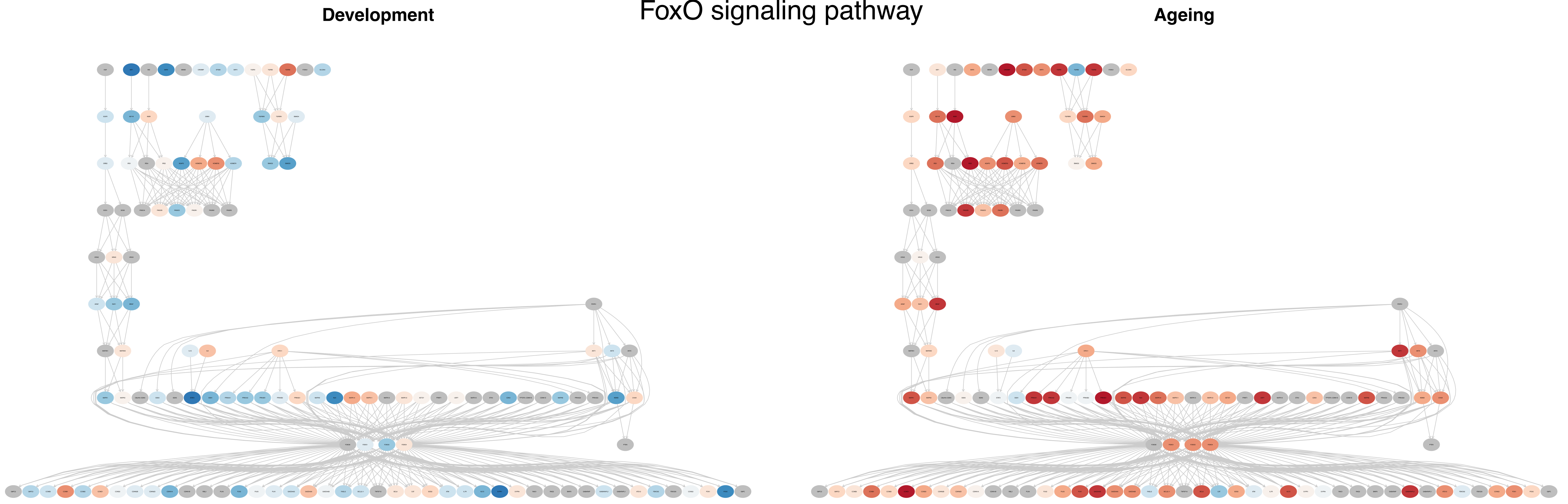
FoxO signaling pathway
Development
Ageing

### Autophagy – animal

**Development** **Ageing**

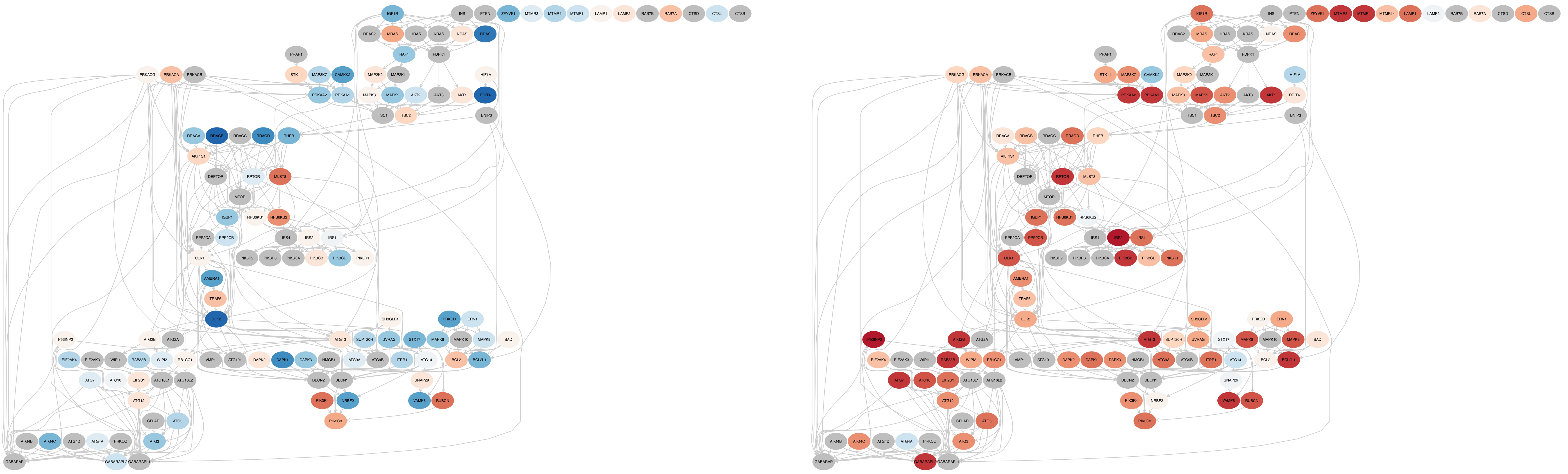

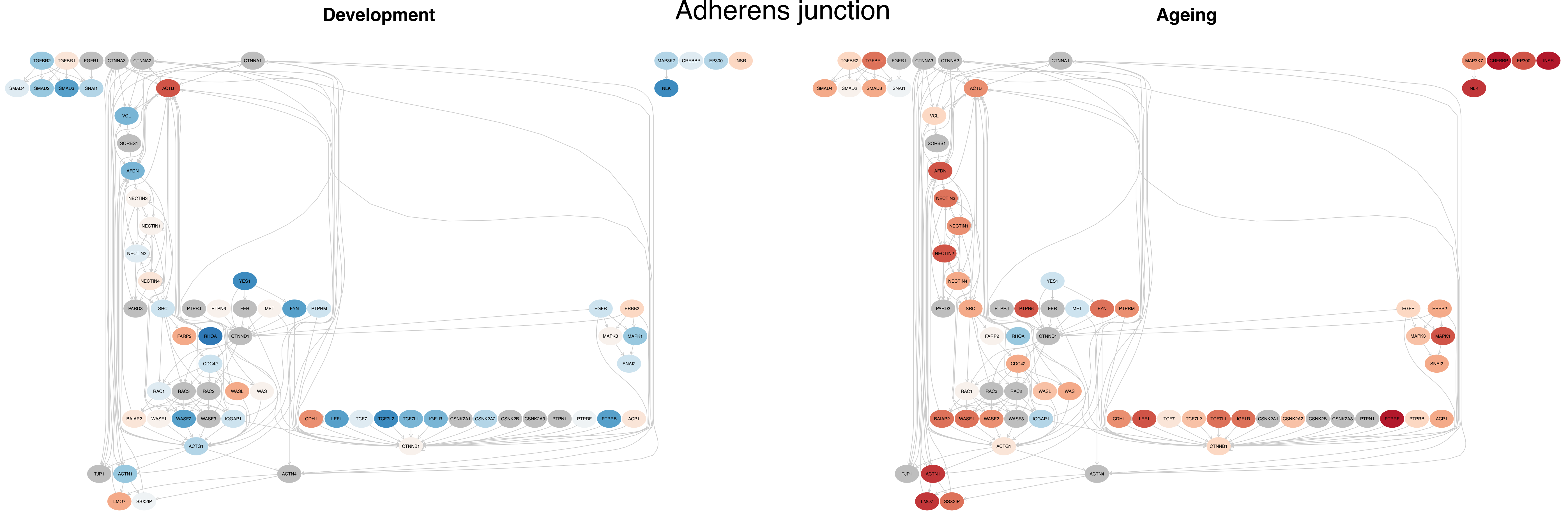

Adherens junction
Development
Ageing
TGFBR2
TGFBR1
FGFR1
CTNNA3
CTNNA2
CTNNA1
MAP3K7
CREBBP
EP300
INSR
SMAD4
SMAD2
SMAD3
SNAI1
ACTB
NLK
VCL
SORBS1
AFDN
NECTIN3
NECTIN1
NECTIN2
NECTIN4
YES1
PARD3
SRC
PTPRJ
PTPN6
FER
MET
FYN
PTPRM
EGFR
ERBB2
FARP2
RHOA
CTNND1
MAPK3
MAPK1
CDC42
SNAI2
RAC1
RAC3
RAC2
WASL
WAS
BAIAP2
WASF1
WASF2
WASF3
IQGAP1
CDH1
LEF1
TCF7
TCF7L2
TCF7L1
IGF1R
CSNK2A1
CSNK2A2
CSNK2B
CSNK2A3
PTPN1
PTPRF
PTPRB
ACP1
ACTG1
CTNNB1
TJP1
ACTN1
ACTN4
LMO7
SSX2IP

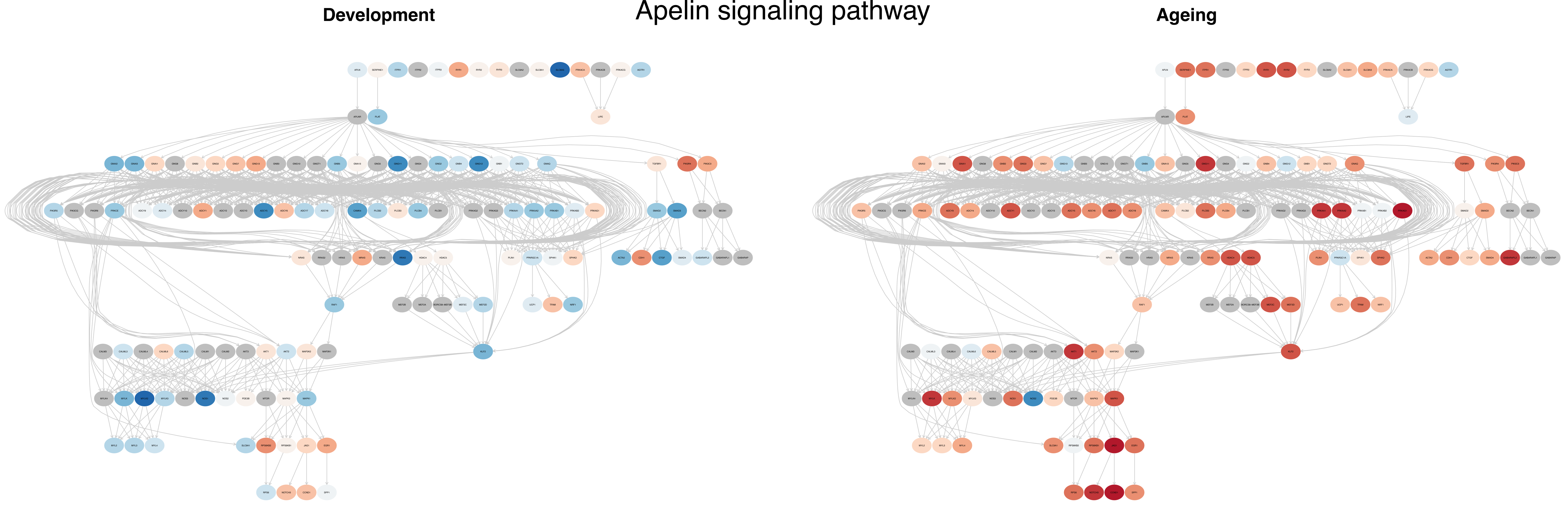
Development
Apelin signaling pathway
Ageing

### mTOR signaling pathway

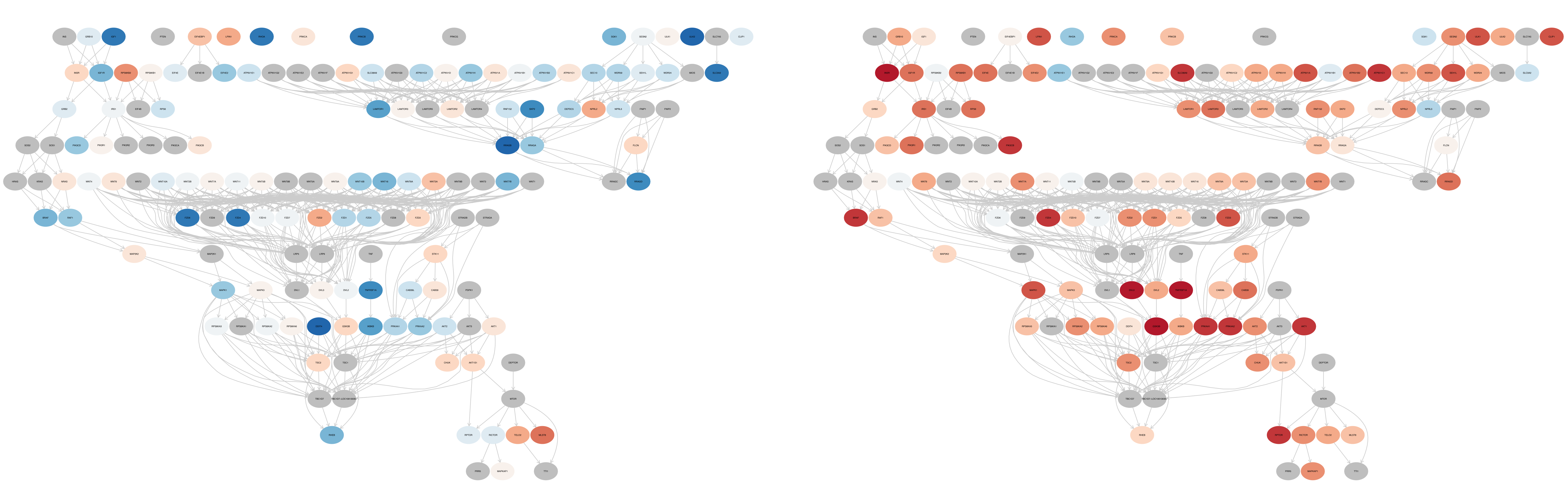

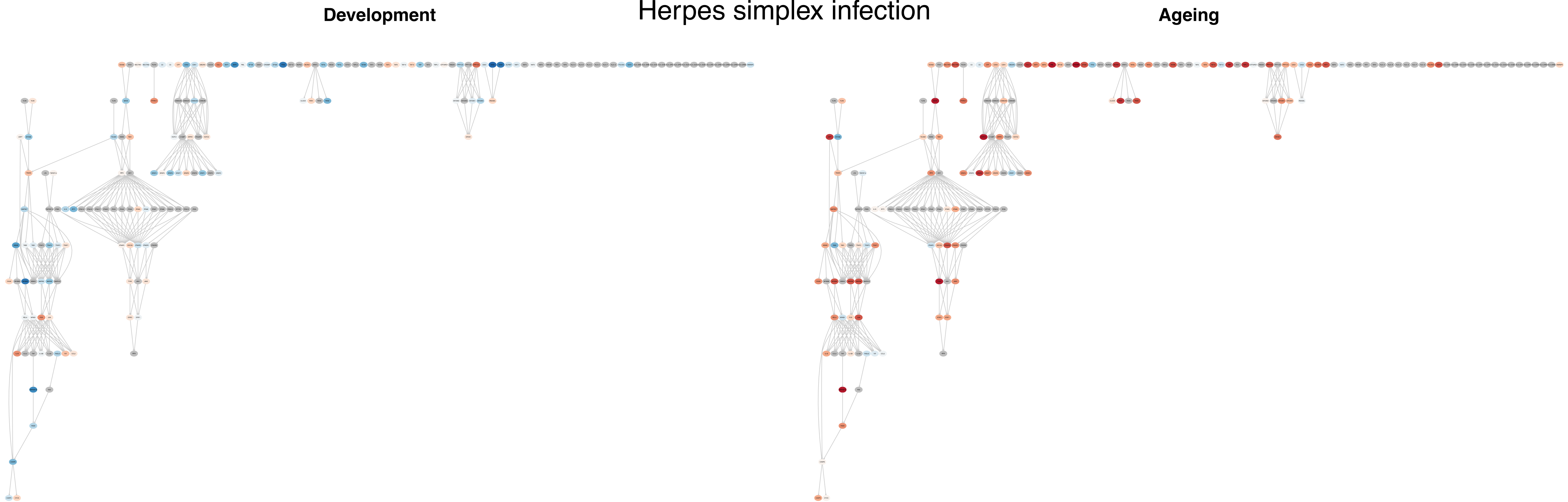
Herpes simplex infection
Development
Ageing

### Axon guidance

**Development** | **Ageing**

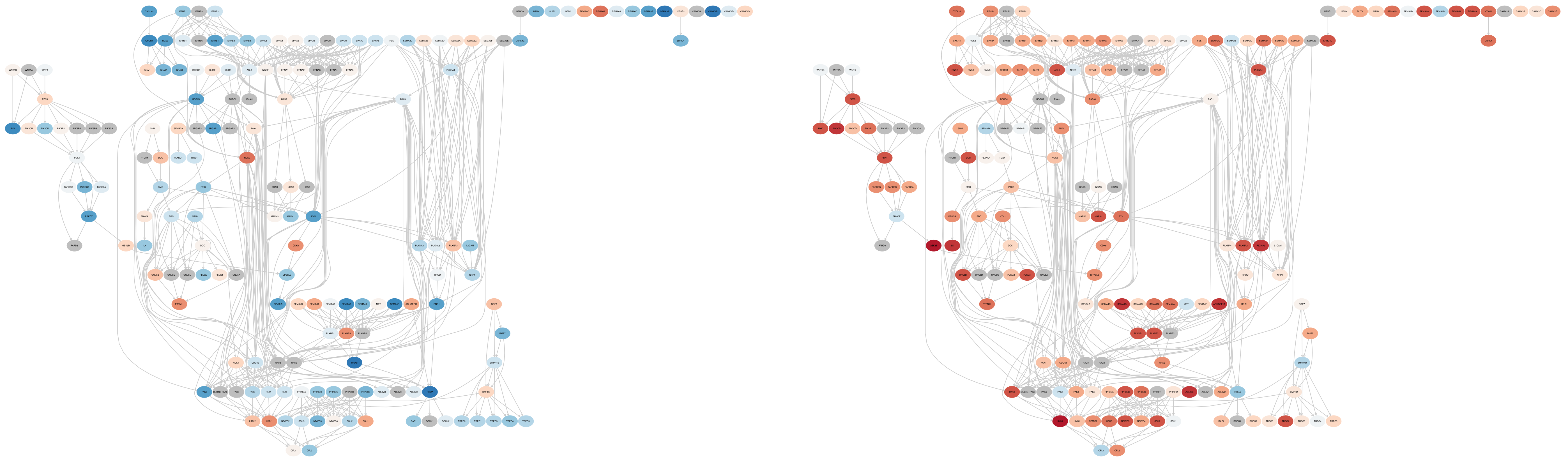

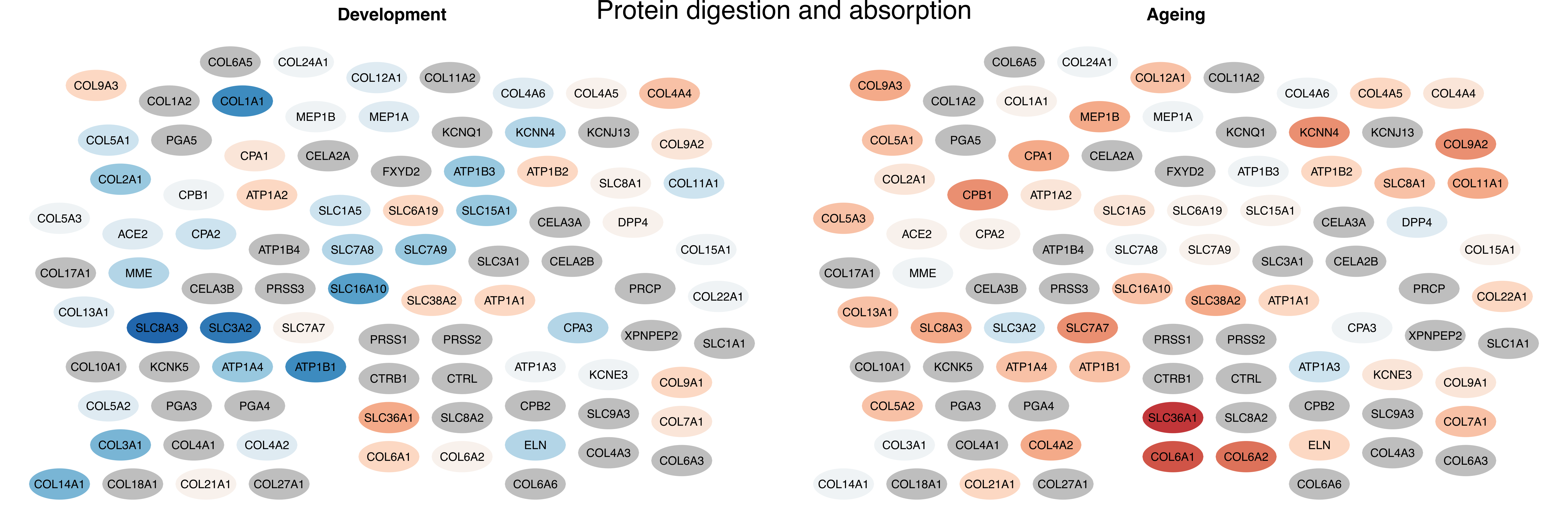

Protein digestion and absorption
Development
COL6A5
COL24A1
COL12A1
COL11A2
COL9A3
COL1A2
COL1A1
COL4A6
COL4A5
COL4A4
MEP1B
MEP1A
KCNQ1
KCNN4
KCNJ13
COL5A1
PGA5
CPA1
CELA2A
COL9A2
FXYD2
ATP1B3
ATP1B2
COL2A1
SLC8A1
COL11A1
CPB1
ATP1A2
SLC1A5
SLC6A19
SLC15A1
COL5A3
CELA3A
DPP4
ACE2
CPA2
ATP1B4
SLC7A8
SLC7A9
COL15A1
SLC3A1
CELA2B
COL17A1
MME
CELA3B
PRSS3
SLC16A10
PRCP
COL22A1
SLC38A2
ATP1A1
COL13A1
SLC8A3
SLC3A2
SLC7A7
CPA3
XPNPEP2
PRSS1
PRSS2
SLC1A1
COL10A1
KCNK5
ATP1A4
ATP1B1
ATP1A3
KCNE3
CTRB1
CTRL
COL9A1
COL5A2
PGA3
PGA4
CPB2
SLC9A3
SLC36A1
SLC8A2
COL7A1
COL3A1
COL4A1
COL4A2
ELN
COL4A3
COL6A1
COL6A2
COL6A3
COL14A1
COL18A1
COL21A1
COL27A1
COL6A6
Ageing
COL6A5
COL24A1
COL12A1
COL11A2
COL9A3
COL1A2
COL1A1
COL4A6
COL4A5
COL4A4
MEP1B
MEP1A
KCNQ1
KCNN4
KCNJ13
COL5A1
PGA5
CPA1
CELA2A
COL9A2
FXYD2
ATP1B3
ATP1B2
COL2A1
SLC8A1
COL11A1
CPB1
ATP1A2
SLC1A5
SLC6A19
SLC15A1
COL5A3
CELA3A
DPP4
ACE2
CPA2
ATP1B4
SLC7A8
SLC7A9
COL15A1
SLC3A1
CELA2B
COL17A1
MME
CELA3B
PRSS3
SLC16A10
PRCP
COL22A1
SLC38A2
ATP1A1
COL13A1
SLC8A3
SLC3A2
SLC7A7
CPA3
XPNPEP2
PRSS1
PRSS2
SLC1A1
COL10A1
KCNK5
ATP1A4
ATP1B1
ATP1A3
KCNE3
CTRB1
CTRL
COL9A1
COL5A2
PGA3
PGA4
CPB2
SLC9A3
SLC36A1
SLC8A2
COL7A1
COL3A1
COL4A1
COL4A2
ELN
COL4A3
COL6A1
COL6A2
COL6A3
COL14A1
COL18A1
COL21A1
COL27A1
COL6A6

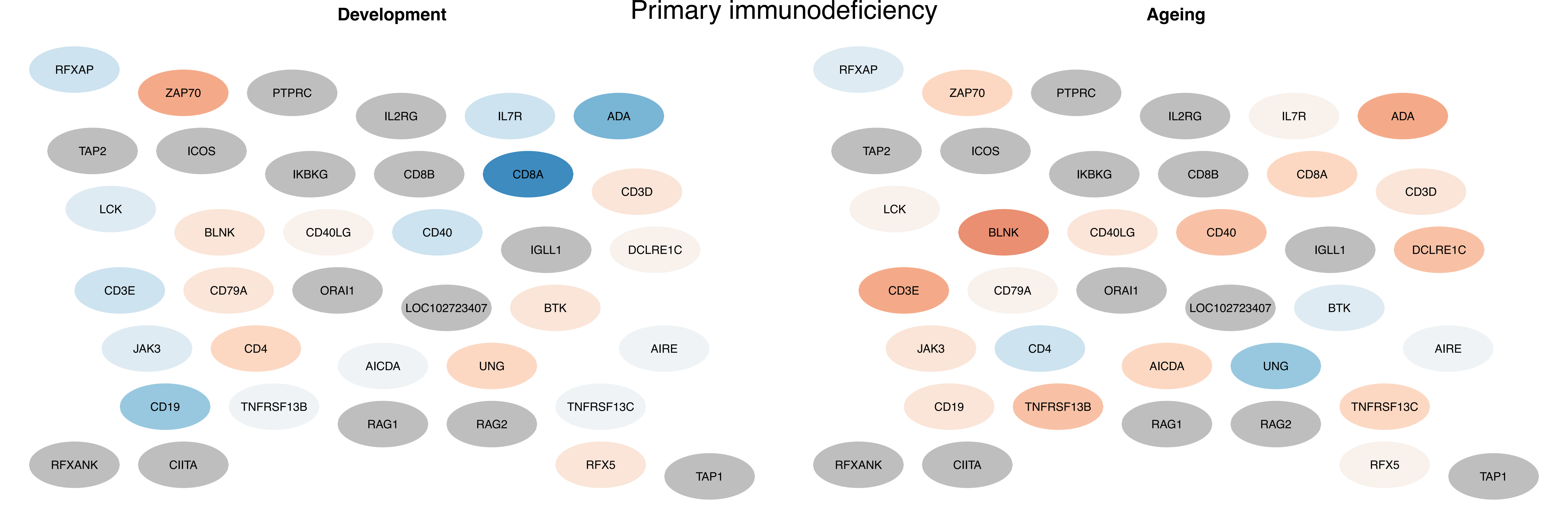
Primary immunodeficiency
Development
Ageing
RFXAP
ZAP70
PTPRC
IL2RG
IL7R
ADA
TAP2
ICOS
IKBKG
CD8B
CD8A
CD3D
LCK
BLNK
CD40LG
CD40
IGLL1
DCLRE1C
CD3E
CD79A
ORAI1
LOC102723407
BTK
JAK3
CD4
AICDA
UNG
AIRE
CD19
TNFRSF13B
RAG1
RAG2
TNFRSF13C
RFXANK
CIITA
RFX5
TAP1

### Linoleic acid metabolism

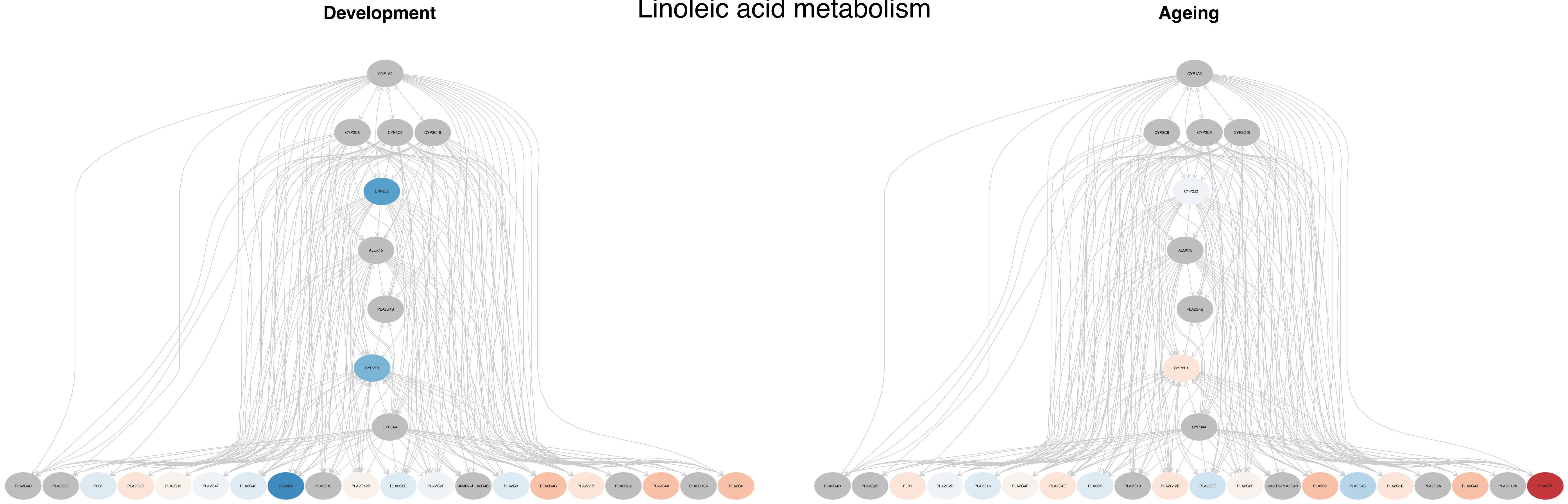

### Fat digestion and absorption

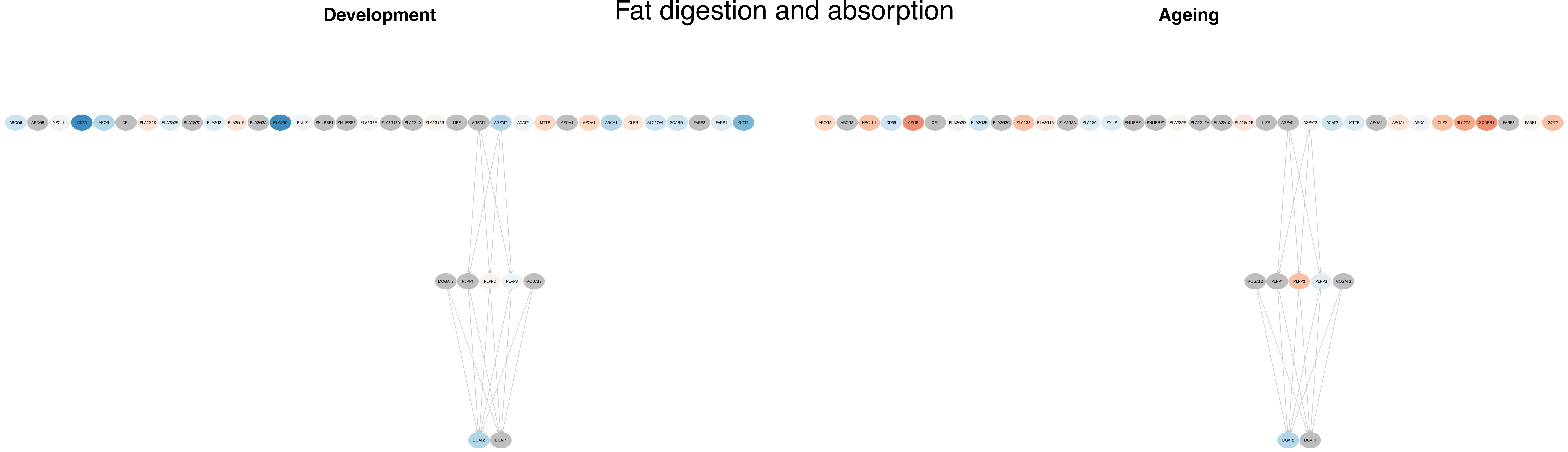
